# Evolutionary diversification of the nuclear pore complex in *Entamoeba histolytica* reveals conserved and lineage-specific nucleoporins

**DOI:** 10.64898/2026.09.13.750810

**Authors:** Huda Amilina, Herbert J. Santos, Kenichiro Imai, Tomoyoshi Nozaki

## Abstract

Nuclear pore complexes (NPCs) are the gateways for macromolecular exchange between the nucleus and cytoplasm. Although NPC architecture is broadly conserved across eukaryotes, substantial lineage-specific diversification has emerged, particularly among divergent protists. Here, we investigated the NPC of *Entamoeba histolytica*, an evolutionarily divergent amoebozoan and the causative agent of amebiasis. Using the FG-repeat nucleoporin EhNup98-like as bait in affinity purification coupled with mass spectrometry, we identified various associated proteins, including a previously uncharacterized candidate nucleoporin with similarity to Nup53/Nup35. Reciprocal proteomic analysis of EhNup53-like further recovered a broader repertoire of candidate NPC components, including proteins corresponding to conserved nucleoporins as well as several uncharacterized nuclear pore-associated proteins. Structural analyses indicated that this EhNup53-like protein retains a conserved RNA recognition motif (RRM)-like fold despite extensive primary-sequence divergence, while displaying lineage-specific features, including an expanded repeat-rich region and truncation of the C-terminal region typically associated with Nup155 binding. Together, these findings expand the known candidate nucleoporins in *E. histolytica* and reveal NPC-associated networks that combines recognizable conserved components with extensively remodeled and poorly characterized proteins. Our study highlights the evolutionary plasticity of the NPC and provides a framework for understanding how nuclear pore architecture has diversified across deeply divergent eukaryotic lineages.

## Introduction

The emergence of the nucleus was a defining transition in eukaryotic evolution, creating a physical boundary between the genome and the cytoplasm and requiring dedicated mechanisms for macromolecular exchange. By the time of the Last Eukaryotic Common Ancestor (LECA), the nuclear envelope and nuclear pore complexes (NPCs) were already established features of eukaryotic cells, indicating that regulated nucleocytoplasmic transport arose early in eukaryotic evolution(1). As eukaryotes subsequently diversified, this ancient transport machinery was retained across lineages while its molecular composition underwent extensive evolutionary modification(2). Understanding how such an essential cellular machine tolerates this divergence provides a fundamental opportunity to examine the relationship between evolutionary conservation, molecular architecture, and cellular function.

NPCs are large multiprotein assemblies that form selective gateways through the nuclear envelope. In well-studied opisthokont model organisms, approximately 30 distinct nucleoporins (Nups), present in multiple copies, assemble into cytoplasmic filaments, outer and inner ring scaffolds, a central transport channel, and a nuclear basket(3,4). FG-repeat Nups occupy the central channel and generate a selective permeability barrier while providing transient interaction sites for soluble nuclear transport receptors. These structural and dynamic components allow rapid exchange of proteins and RNAs while maintaining nuclear compartmentalization(5,6).

Our current understanding of NPC architecture, however, is derived predominantly from fungi and metazoans. High-resolution structures from *Saccharomyces cerevisiae* and vertebrates have revealed striking similarities in the overall organization of the pore, supporting the concept of a conserved architectural framework(7). Yet opisthokonts represent only one branch of the vast diversity of eukaryotes. Studies of divergent protists increasingly show that conservation of NPC function does not necessarily require conservation of its apparent molecular composition(8). In protozoan parasites such as *Trypanosoma brucei*, for example, the nuclear basket contains highly divergent components, whereas canonical transmembrane Nups have not been readily identified in *Plasmodium* species(9,10). Such findings suggest that the NPC is evolutionary more plastic than implied by classical model systems. This evolutionary plasticity highlights that the essential eukaryotic cellular machines can undergo extensive molecular divergence while preserving their core architectural and functional properties.

Extensive sequence divergence can obscure evolutionary relationships among nucleoporins even when their structural organization or functional position within the NPC remains conserved(11). FG-repeat Nups provide useful entry points into such poorly characterized NPCs because their characteristic sequence features and transport-related functions can remain recognizable even when overall primary-sequence conservation is limited(12). Nup98 is especially informative in this context. Its FG-repeat region contributes to the selective transport barrier and interacts with nuclear transport receptors, while Nup98 also participates in mRNA export and gene regulation through interactions with factors including Rae1 and TAP/Mex67(13,14). In many eukaryotes, Nup98 additionally contains a conserved C-terminal nucleoporin-2 domain belonging to the peptidase S59 family, which mediates autoproteolytic processing(15). These combined features provide molecular signatures through which highly divergent Nup98-related proteins can be identified and used as starting points for investigating otherwise poorly defined nuclear pores.

*Entamoeba histolytica* provides a particularly informative system in which to examine NPC divergence(8). This unicellular parasite belongs to Amoebozoa, a major eukaryotic lineage that diverged deeply from the opisthokont organisms in which NPC structure has been most extensively characterized. *E. histolytica* is also the causative agent of human amebiasis, yet fundamental aspects of its nuclear organization remain poorly understood. Despite its requirement for regulated nucleocytoplasmic transport, the molecular composition of *E. histolytica* NPC is largely unknown(16). Comparative sequence analysis have failed to identify clear orthologs of several canonical nucleoporins(17). This apparent absence may reflect extensive primary-sequence divergence that obscures conserved structural relationships, lineage-specific loss or remodelling of canonical components, or the recruitment of previously uncharacterized proteins to perform equivalent NPC functions. The *E. histolytica* NPC therefore offers an opportunity to investigate how a deeply divergent eukaryote preserves the organization and function of an essential nuclear transport machinery despite substantial molecular divergence.

Here, we used a conserved Nup98-like protein as an entry point to investigate NPC organization in *E. histolytica*. By integrating comparative sequence and structural analysis with biochemical characterization, we characterized EhNup98-like protein and defined its associated protein network. This network contained 10 candidate homologs of established NPC components, including Rae1, Nup62, Nup93, Nup155, and Nup205(17), together with EHI_026310, a previously uncharacterized protein that emerged as a highly divergent Nup53/Nup35-related nucleoporin that we selected for further characterization. Our findings reveal a conserved molecular relationship embedded within a highly divergent amoebozoan NPC and demonstrate that core features of nuclear pore complex organization can persist despite extensive remodelling of nucleoporin sequences. More broadly, they illustrate how essential eukaryotic cellular machines can preserve fundamental architectural and functional principles while evolving markedly different molecular implementations.

## Results

### In silico characterization and domain architecture of EhNup98-like protein

To define the molecular features of the putative *E. histolytica* Nup98 homolog (EhNup98-like hereinafter; AmoebaDB accession number: EHI_179400), we first examined its sequence and predicted structural organization. Comparison with human Nup98 (HsNup98; UniProt ID: P52948) showed that, despite substantial overall sequence divergence, EhNup98-like retains two characteristic features of canonical Nup98 proteins: an approximately 80-amino-acid N-terminal FG-repeat region containing multiple phenylalanine-glycine (FG) motifs, and a conserved C-terminal domain belonging to the Peptidase S59 family (Fig 1A). Alternative splicing of human Nup98 gives rise to six annotated protein isoforms in UniProt. Isoforms 1, 2, 5, and 6 represent Nup98–Nup96 precursor-type proteins containing the C-terminal Nup96 region, whereas isoforms 3 and 4 are shorter Nup98-specific forms lacking Nup96(18). The FG-repeat region of EhNup98-like is shorter and compositionally distinct from that of HsNup98, whereas much of the intervening sequence between the FG-repeat region and the C-terminal domain of EhNup98-like is predicted to be intrinsically disordered like HsNup98. In addition, a recognizable Gle2-binding sequence (GLEBS) motif, which mediates interaction of canonical Nup98 proteins with the mRNA export factor Rae1/Gle2(13), was not identified in EhNup98-like.

**Fig 1.**
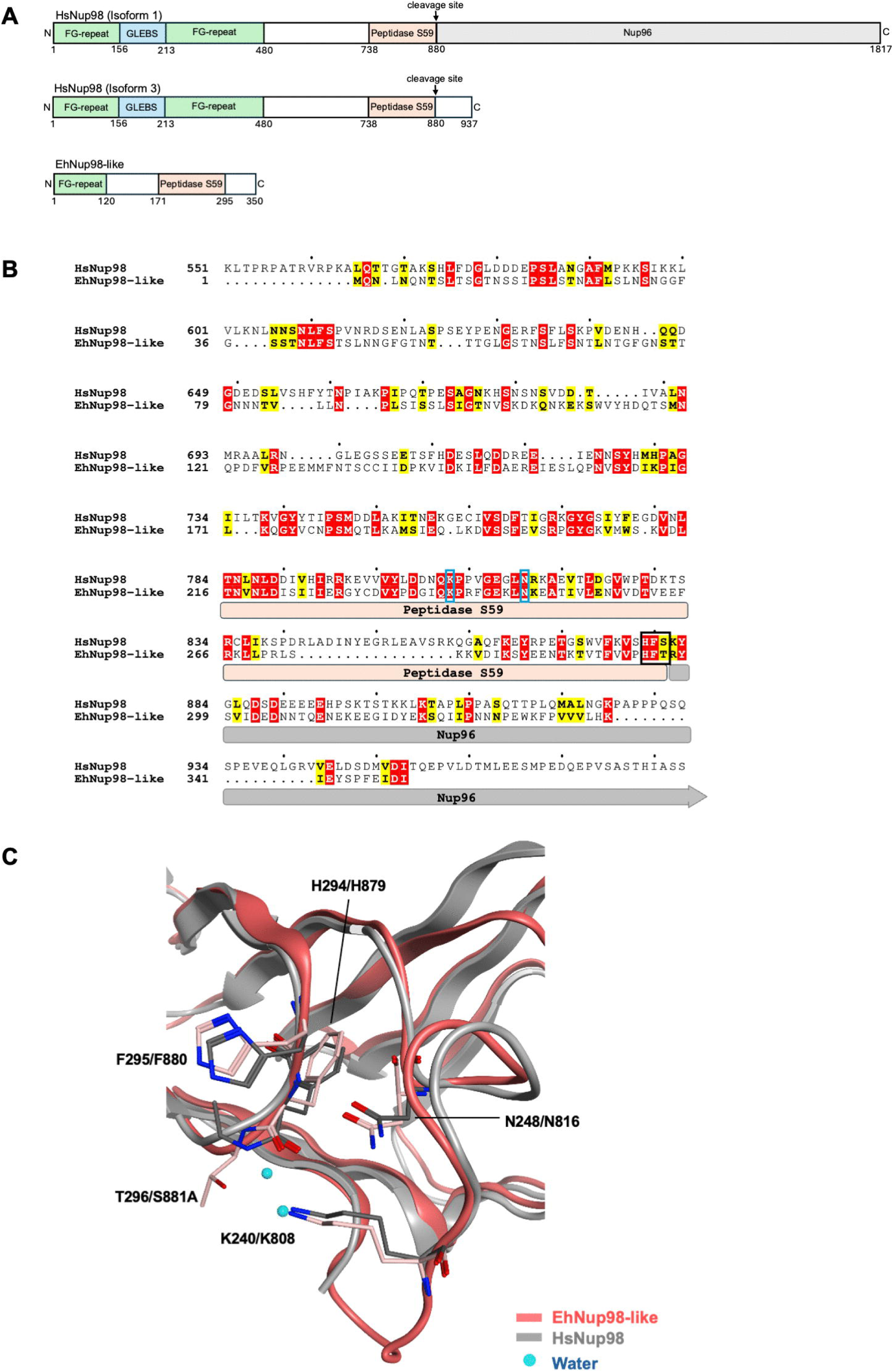
Domain architecture and structural conservation of *E. histolytica* Nup98-like. **(A)** Schematic representation of the domain organization of human Nup98 (HsNup98) and *E. histolytica* Nup98-like (EhNup98-like). FG-repeat regions and the conserved C-terminal Peptidase S59 domain are indicated, along with predicted unstructured regions. HsNup98 isoforms 1 and 3 are shown as representative Nup98-Nup96 precursor and Nup98-specific architectures, respectively. **(B)** Amino acid sequence alignment of the C-terminal Peptidase S59 domain highlighting the position of the catalytic HFS/T motif (black box). Conserved residues are shown as white letters on a red background, while residues with similar physicochemical properties are highlighted in yellow. K240 and N248 as catalytic residues are indicated in blue. Sequence alignment and visualization are performed using Clustal Omega and ESPript 3.2, respectively(19). **(C)** Homology model of the EhNup98-like autoproteolytic domain (pink) superimposed on the human Nup98 structure (grey, PDB ID: 2Q5X), showing spatial conservation of key catalytic residues (K240, N248, H294, and F295). Water molecules are shown as cyan spheres. Homology model of the EhNup98-like autoproteolytic domain was constructed by homology modeling based on human Nup98 structure.

Because autoproteolytic processing is a characteristic property of Nup98 proteins, we next examined conservation of the residues associated with this activity. Alignment of the C-terminal region revealed a conserved HFT motif corresponding to the autoproteolytic cleavage region of HsNup98 (Fig 1B, S1A Fig). Several residues surrounding this motif were also conserved, suggesting preservation of features associated with the Nup98 autoproteolytic domain despite extensive divergence elsewhere in the protein.

We next compared the predicted 3D structure of the EhNup98-like C-terminal domain with the experimentally determined structured of HsNup98. Structural superposition showed conservation of the overall fold of the peptidase S59 domain (Fig 1C). Residues K240, N248, H294, and F295 of EhNup98-like, corresponding to residues implicated in the Nup98 autoproteolytic mechanism, occupied similar spatial positions to their counterparts in HsNup98. In particular, the conserved H294-F295 residues were positioned within the predicted catalytic region, consistent with retention of the structural arrangement required for autoproteolytic processing.

To determine whether these features are conserved within the *Entamoeba* lineage, we compared EhNup98-like with homologs from *E. dispar*, *E. nutalli*, *E. moshkovskii*, and *E. invadens*. All five *Entamoeba* proteins retained a conserved C-terminal Peptidase S59 domain, including the characteristic HFT motif, despite substantial divergence in the surrounding sequences (S1B Fig). Thus, although the overall primary sequence of EhNup98-like is highly divergent, its FG-repeat architecture and autoproteolytic domain retain key molecular features characteristic of Nup98 proteins.

These findings support the identification of EHI_179400 as an *E. histolytica* Nup98-like protein and suggest that the structural basis underlying Nup98 autoproteolytic processing is conserved despite extensive sequence divergence.

### EhNup98-like localizes to the nuclear periphery and is enriched in nuclear fractions

To determine whether EhNup98-like associates with the nuclear pore complex in cells, we generated an N-terminal HA-tagged construct (HA-Nup98-like) and expressed it in *E. histolytica* trophozoites. The full-length HA-tagged protein has a predicted molecular weight of approximately 42 kDa. Immunoblot of whole-cell lysates with an anti-HA antibody detected a predominant band at approximately 37 kDa, close to the predicted size of the N-terminal product generated following autoproteolytic cleavage (∼35.5 kDa), together with a weaker upper band at approximately 42 kDa corresponding to the expected size of the full-length precursor (Fig 2A). This migration pattern is consistent with proteolytic processing of EhNup98-like and supports the predicted activity of its conserved C-terminal peptidase S59 domain. Cysteine synthase 1 (CS1) was used as a loading control(20).

**Fig 2.**
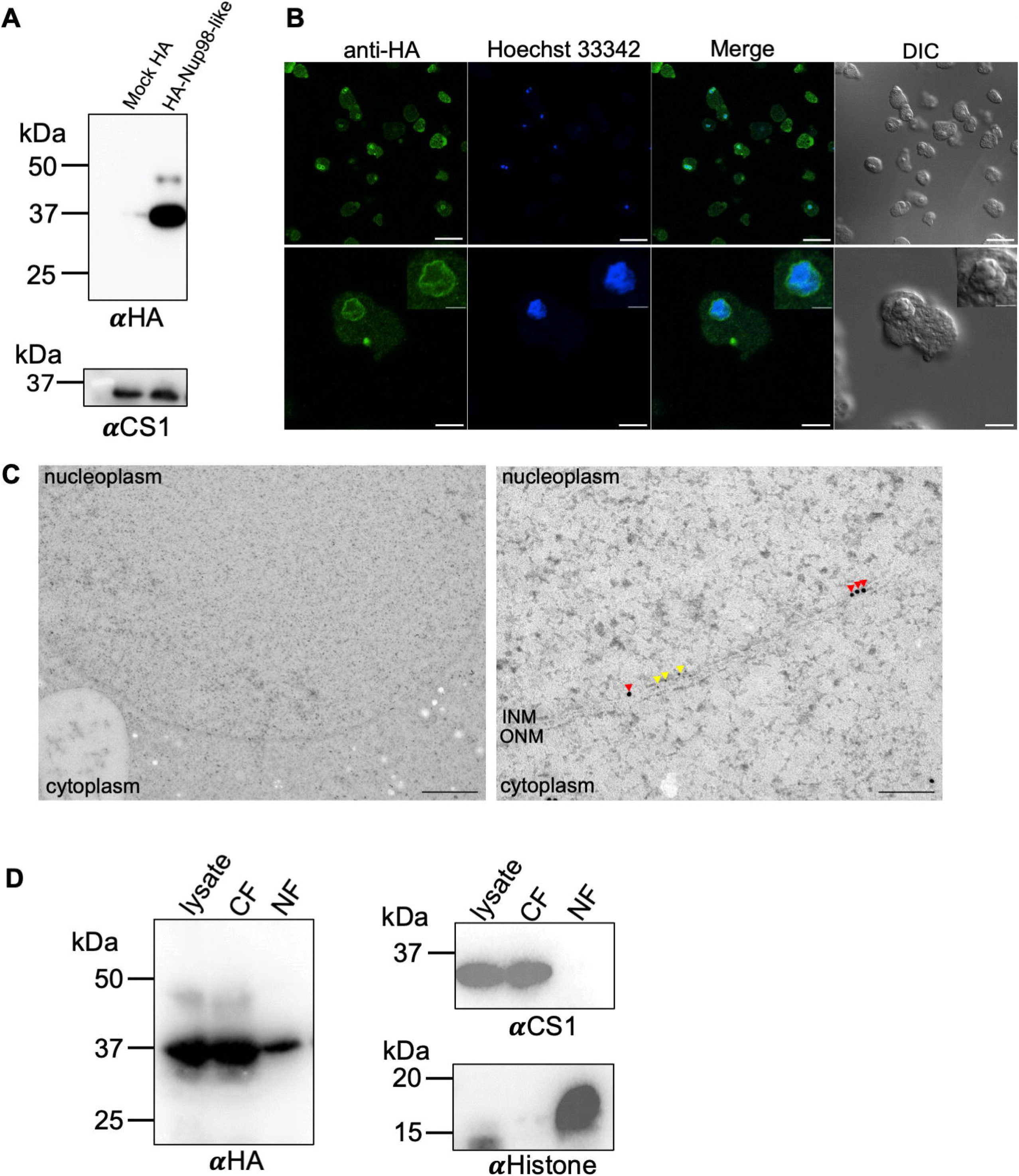
EhNup98-like localizes to the nuclear periphery and is enriched in nuclear fractions. **(A)** Immunoblot analysis of HA-Nup98-like expression in *E. histolytica* trophozoites. Whole-cell lysates probed with an anti-HA antibody reveals a predominant band at ∼37 kDa, corresponding to the processed N-terminal fragment, and a weaker band at ∼42 kDa, representing the full-length precursor. Cysteine synthase 1 (CS1) was used as a loading control. **(B)** Indirect immunofluorescence assay (IFA) of HA-Nup98-like localization. Cells were stained with anti-HA antibody and co-stained with Hoechst 33342 to visualize nuclei. Scale bar, 40 µm in the overview images, 10 µm in the enlarged images, and 4 µm in the insets. DIC, differential interference contrast. **(C)** Ultrastructural localization of HA-Nup98-like using immuno-electron microscopy (immunoEM). Ultrathin sections were dual-labeled with mouse anti-HA antibody (10 nm anti-mouse gold particles; red arrowheads) detecting HA-Nup98-like and rabbit anti-laminB1 antibody (5 nm anti-rabbit gold particles; yellow arrowheads) marking the nuclear envelope. ONM, outer nuclear membrane; INM, inner nuclear membrane. Scale bar, 1 µm in low magnification image, 100 nm in high magnification image **(D)** Subcellular fractionation of HA-Nup98-like. Whole lysate, cytoplasmic (CF) and nuclear fractions (NF) were analyzed by immunoblotting. CS1 and Histone H3 serve as cytoplasmic and nuclear fractions, respectively. Uncropped western blots available in S1 Raw Images.

We next examined the subcellular distribution of HA-Nup98-like by indirect immunofluorescence assay (IFA). HA-Nup98-like showed prominent nuclear-associated localization in approximately 70% of the analyzed cells, with the signal frequently concentrated at the nuclear periphery (Fig 2B). A diffuse cytoplasmic signal was also observed, indicating the presence of an additional cytosolic pool of HA-Nup98-like.

To resolve its localization at higher spatial resolution, we performed dual-labeling immuno-electron microscopy (immunoEM) on ultrathin sections of the trophozoites. HA-Nup98-like was detected with a mouse anti-HA primary antibody and followed by a 10-nm gold-conjugated anti-mouse secondary antibody, whereas lamin B1 was detected with a rabbit anti-lamin B1 primary antibody and a 5-nm gold-conjugated anti-rabbit secondary antibody as a positional marker for the nuclear envelope. HA-Nup98-like labeling was enriched at the nuclear membrane and was frequently observed adjacent to lamin B1 signal (Fig 2C). Notably, 10 nm gold particles corresponding to HA-Nup98-like were also detected at electron-dense discontinuities along the nuclear envelope, consistent with the expected position of NPCs. These observations further provide ultrastructural evidence supporting the localization of EhNup98-like with the nuclear periphery and its association with the NPC.

We independently assessed this distribution by subcellular fractionation. Cytoplasmic and nuclear fractions were validated using CS1 and histone H3 as cytoplasmic and nuclear markers, respectively. HA-Nup98-like was detected in both fractions but was clearly enriched in the nuclear fraction (Fig 2D), consistent with its predominant nuclear-peripheral localization observed by microscopy. The presence of HA-Nup98-like in the cytoplasmic fraction was also consistent with the diffuse cytosolic signal detected by immunofluorescence.

To examine whether the conserved autoproteolytic motif contributes to EhNup98-like processing and localization, we generated a mutant in which the threonine residue of the HFT motif was substituted with alanine (HA-Nup98-like^T296A^). Immunoblot analysis showed that the mutant retained a predominant band at approximately 37 kDa, however, the upper band was more prominent than in wild type HA-Nup98-like (S2A Fig). The increased abundance of the upper band is consistent with accumulation of the full-length precursor and suggests that substitution of T296 reduces, but does not completely abolish, processing of EhNup98-like. The T296A substitution also altered the subcellular distribution of EhNup98-like. Whereas wild-type HA-Nup98-like showed prominent enrichment at the nuclear periphery, HA-Nup98-like^T296A^ displayed increased cytoplasmic localization and reduced accumulation at the nuclear rim (S2B Fig). These findings indicate that perturbation of the conserved HFT region affects both the processing pattern and cellular distribution of EhNup98-like, suggesting that the integrity of the autoproteolytic domain contributes to efficient association of the protein with the nuclear periphery.

### Silencing of *EhNup98-like* impairs trophozoite proliferation and promotes nuclear accumulation of poly(A)^+^ RNA

To investigate the functional contribution of EhNup98-like, we generated an *EhNup98-like* gene-silencing strain (Nup98-like-gs) and compared it with the pSAP2 (empty-vector) control strain. Reduction of EhNup98-like expression was confirmed by PCR analysis using EhNup98-like specific primers. Whereas a clear amplification product was detected in the pSAP2 control, the corresponding signal was markedly reduced in Nup98-like-gs cells (Fig3A). Amplification of RNA polymerase II (RNA polII) was comparable between the two strains, supporting the specific reduction of *EhNup98-like* mRNA expression.

We next examined whether depletion of EhNup98-like affected trophozoite proliferation. The pSAP2 control population increased progressively throughout the 96 h observation period, reaching approximately 3 × 10^5^ cells/mL. In contrast, Nup98-like-gs trophozoites exhibited substantially slower growth, with cell numbers remaining markedly lower throughout the experiment and reaching only 5 × 10^4^ cells/mL at 96 h (Fig 3B). These results suggest reduced EhNup98-like expression strongly impaired proliferation of *E. histolytica* trophozoites.

**Fig 3.**
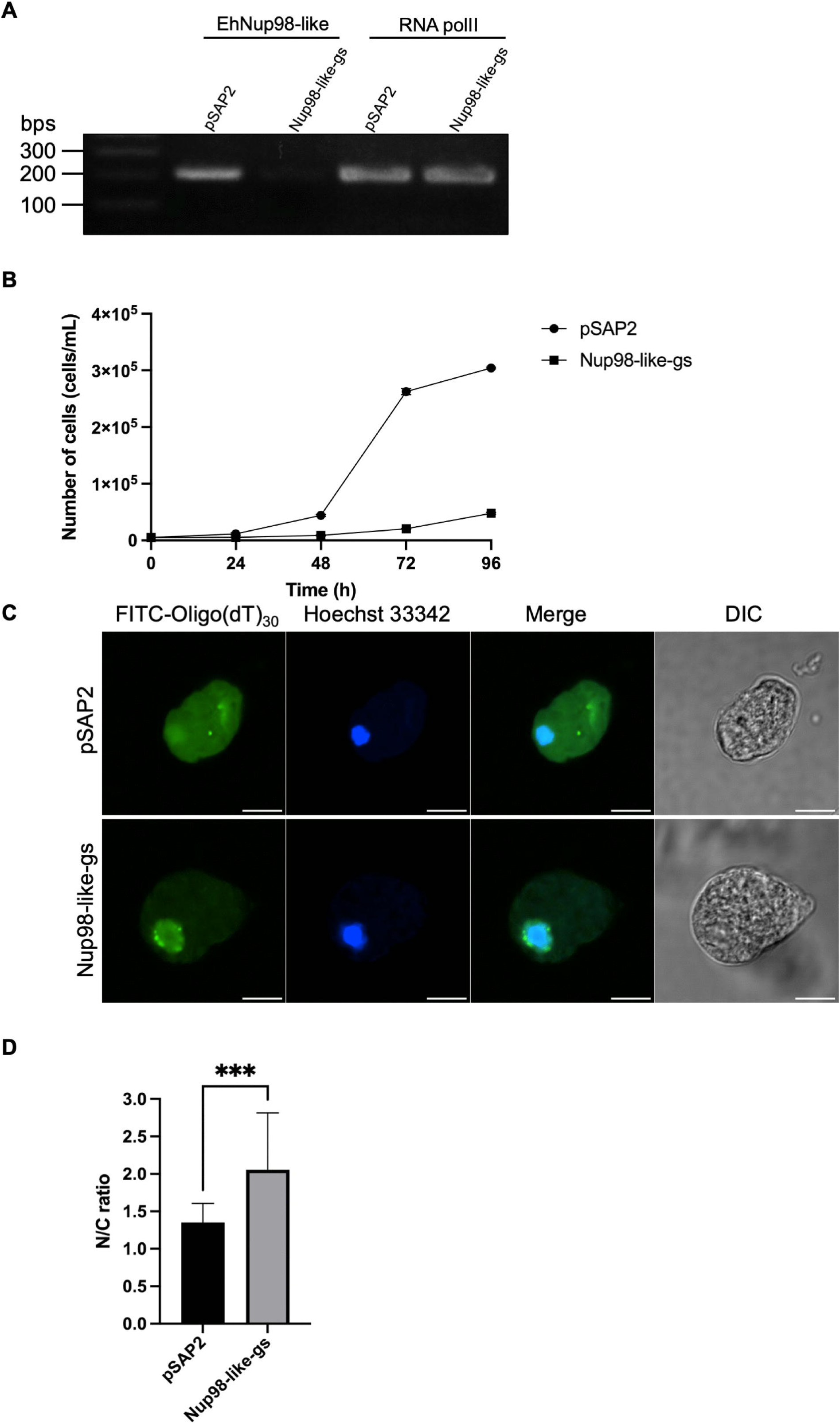
Silencing of EhNup98-like impairs trophozoite proliferation and promotes nuclear accumulation of poly(A)^+^ RNA. **(A)** Confirmation of EhNup98-like gene silencing in the Nup98-like-gs strain by RT-PCR. Uncropped gel available in S1 Raw Images. **(B)** Growth kinetics of pSAP2 control and Nup98-like-gs trophozoites over 96 h. Cell numbers were determined at the indicated time points. **(C)** Representative fluorescence in situ hybridization (RNA-FISH) images showing the intracellular distribution of poly(A)^+^ RNA in pSAP2 and Nup98-like-gs trophozoites. Poly(A)^+^ RNA was detected using FITC-conjugated oligo(dT)_30_ probes (green), and nuclei were stained with Hoechst 33342 (blue). Differential interference contrast (DIC) images are shown on the right. Scale bars, 10 µm. (D) Quantification of the nuclear-to-cytoplasmic (N/C) fluorescence intensity ratio of poly(A)^+^ RNA. A total of 66 cells per group were analyzed. Nup98-like-gs trophozoites showed a significantly increased N/C ratio compared with pSAP2 controls. Error bars indicate standard deviation. ***p < 0.001; statistical analysis was performed as described in the Methods.

Because Nup98 participates in mRNA export in other eukaryotes, we next examined the intracellular distribution of polyadenylated RNA [poly(A)^+^] using fluorescence in situ hybridization (RNA-FISH) with FITC-conjugated oligo(dT)_30_ probes. In pSAP2 control cells, poly(A)^+^ RNA was distributed predominantly throughout the cytoplasm with comparatively limited enrichment within the nuclear region (Fig 3C). In contrast, Nup98-like-gs cells exhibited pronounced accumulation of poly(A)^+^ RNA within the nucleus, producing an intense signal that substantially overlapped with the Hoechst-stained nuclear compartment. Quantification of the nuclear-to-cytoplasmic (N/C) fluorescence ratio confirmed this redistribution. Nup98-like-gs cells showed a significantly higher poly(A)^+^ RNA N/C ratio than pSAP2 control cells, increasing from approximately 1.3 to 2.0 (n=66, unpaired two-tailed t-test, p = 0.0003; Figure 3C). These findings indicate that depletion of EhNup98-like disrupts the normal nucleocytoplasmic distribution of poly(A)^+^ RNA and is consistent with impaired mRNA export.

### EhNup98-like forms heterogeneous macromolecular assemblies and associates with nuclear transport, protein homeostasis, and gene regulation

To determine whether EhNup98-like is incorporated into native macromolecular assemblies, we first analyzed lysates from trophozoites expressing HA-Nup98-like by blue native PAGE (BN-PAGE). Whole-cell lysates were solubilized with increasing concentrations of digitonin and separated under non-denaturing conditions. Although HA-Nup98-like has a predicted molecular weight of approximately 42 kDa, immunoblotting revealed multiple higher molecular weight bands, with prominent signals at approximately 60 kDa, 150-200 kDa, 280 kDa, and 700 kDa (Fig 4A). Apparent molecular masses were estimated from a calibration curve generated using native protein standards. Parallel analysis under denaturing conditions confirmed the presence of HA-Nup98-like in the samples (Fig 4A, lower panel). These results indicate that EhNup98-like is present in multiple native assemblies rather than predominantly as a monomeric protein.

**Fig 4.**
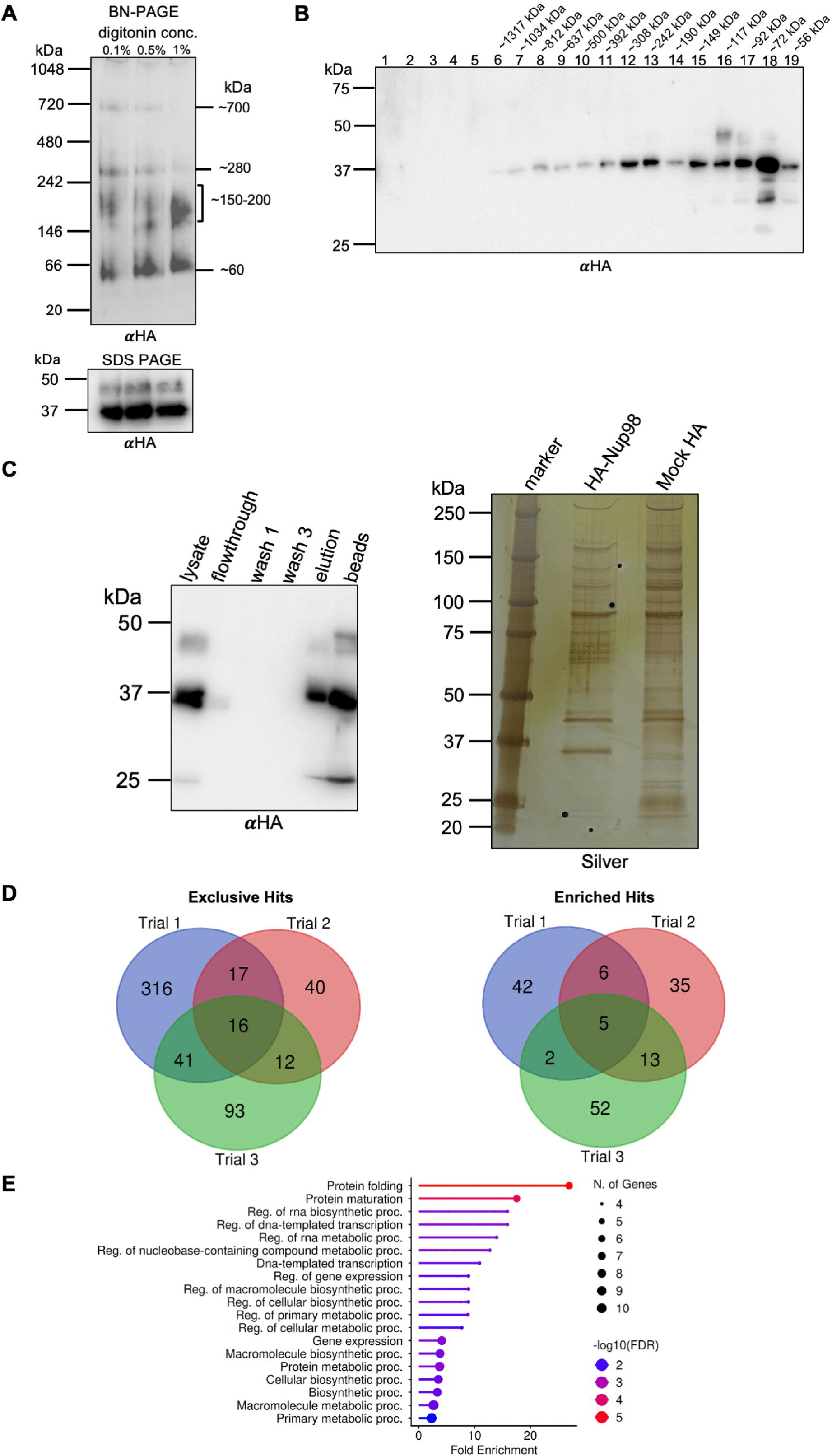
Native complex formation and proteomic identification of the EhNup98-like interactome. **(A)** Blue native PAGE (BN-PAGE) and SDS-PAGE of HA-Nup98-like lysates solubilized in increasing digitonin concentrations (0.1, 0.5, and 1%). **(B)** Size exclusion chromatography (SEC) of HA-Nup98-like lysates. Fractions correspond to approximate molecular weight ranges based on column calibration standards. **(C)** Validation of the co-immunoprecipitation (co-IP) assay by anti-HA immunoblotting. Eluted fractions were analyzed by SDS-PAGE followed by silver staining. Uncropped western blots available in S1 Raw Images. **(D)** Venn diagrams showing overlap of proteins identified across three independent biological replicates of HA–EhNup98-like co-IP/MS. Left, exclusive proteins detected in HA–Nup98-like samples but not in mock controls. Right, enriched proteins detected in both HA–Nup98-like and mock samples with QV ratio ≥ 2.0. **(E)** Gene Ontology (GO) enrichment analysis of the EhNup98-like-associated protein, showing enrichment in protein folding, RNA metabolism, and transcription-related processes.

We independently assessed the size distribution of EhNup98-like assemblies by size-exclusion chromatography (SEC). Consistent with the BN-PAGE results, HA-Nup98-like was detected across a broad series of fractions corresponding to apparent molecular masses ranging from approximately 56 kDa to greater than 1 MDa (Fig 4B). Within this broad distribution, prominent immunoreactive signals were observed in fractions 15–18 (∼72–149 kDa) and fractions 11–13 (∼242–392 kDa), with weaker signals also detected in higher-molecular-mass fractions around ∼637–812 kDa. These SEC profiles broadly corresponded to the major molecular-mass species observed by BN-PAGE. These profiles indicate that EhNup98-like is present in multiple macromolecular assemblies of distinct sizes rather than being restricted to a single stable complex.

To identify proteins associated with these assemblies, we performed co-immunoprecipitation followed by liquid chromatography-tandem mass spectrometry (LC-MS/MS) using lysates from HA-Nup98-like trophozoites and mock HA as controls in three biological replicates. Enrichment of HA-Nup98-like in the elution was confirmed by immunoblotting and silver staining (Fig 4C). Across the combined proteomic datasets from HA-Nup98-like and corresponding mock HA from three biological replicates, a total of 1,842 non-redundant proteins were detected.

To identify reproducible candidate interactors, proteins were classified using two complementary criteria. “Exclusive hits” were defined as proteins detected in HA-Nup98-like but absent from the corresponding mock controls, whereas “enriched hits” were defined as proteins with a quantitative value (QV) ratio of ≥2.0 relative to the mock samples based on normalized spectral counts. Sixteen exclusive proteins were reproducibly detected in all three biological replicates, while five proteins fulfilled the enrichment criteria across all three replicates (Fig4D; Tables 1 and 2).

**Table 1.** Proteins identified as exclusive interactors of HA-Nup98-like by immunoprecipitation–mass spectrometry.

| Accession Number | Annotation | Average QV<br>HA-Nup98-like |
| --- | --- | --- |
| EHI_179400 | Nuclear pore protein, putative | 36.79 |
| EHI_138180 | U-box domain-containing protein | 7.93 |
| EHI_063340 | Prefoldin, alpha subunit | 5.77 |
| EHI_081270 | Prefoldin subunit 3 | 5.64 |
| EHI_026310 | Hypothetical protein | 4.97 |
| EHI_079720 | DnaJ family protein | 4.46 |
| EHI_098400 | Importin beta-3 family protein, putative | 4.12 |
| EHI_155420 | Prefoldin subunit 6 | 2.87 |
| EHI_199060 | Prefoldin subunit beta | 2.82 |
| EHI_193390 | Heat shock protein, Hsp20 family, putative | 2.63 |
| EHI_087320 | MRH domain-containing protein | 2.63 |
| EHI_064590 | PX domain-containing protein | 2.62 |
| EHI_119550 | CCR4/NOT transcription complex subunit 3 | 2.56 |
| EHI_141870 | Protein kinase domain containing protein | 2.27 |
| EHI_039600 | Secreted protein | 1.96 |
| EHI_191910 | Programmed cell death protein 5 | 1.71 |

Proteins were identified by immunoprecipitation–mass spectrometry (IP–MS). Exclusive hits are defined as proteins detected in HA–Nup98-like samples but not in mock controls across three biological replicates. Average QV represents the mean normalized abundance value of HA–Nup98-like. Proteins are ranked by decreasing enrichment. Annotations are based on UniProt.

**Table 2.** Proteins identified as enriched interactors of HA-Nup98-like by immunoprecipitation–mass spectrometry.

| Accession Number | Annotation | QV Ratio<br>(HA–Nup98-like/Mock HA) |
| --- | --- | --- |
| EHI_133920 | TPR repeat protein | 6.2 |
| EHI_178720 | Ubiquitin-like domain-containing protein | 4.8 |
| EHI_008710 | TPR repeat protein | 4.7 |
| EHI_146520 | HEAT repeat domain containing protein | 3.0 |
| EHI_093570 | CCR4–NOT transcription complex subunit 9 | 2.8 |

Proteins were identified by immunoprecipitation–mass spectrometry (IP–MS). Enriched hits are defined as proteins exhibiting an average QV ratio $\geq 2.0$ in HA–EhNup98 samples relative to mock controls across three biological replicates. QV ratio represents the normalized abundance ratio (HA–EhNup98/mock). Proteins are ranked by decreasing enrichment. Annotations are based on UniProt.

Because highly divergent NPC components may not necessarily satisfy these enrichment thresholds, we additionally examined the complete proteomic dataset for proteins annotated or predicted to function as nucleoporins of NPC-associated factors. This analysis recovered eleven candidate components of the *E. histolytica* nucleoporins (Table 3). In addition to the EhNup98-like bait, proteins with predicted relationships to Rae1 (EHI_118780), Gle1 (EHI_082590), Nup62 (EHI_173410), Sec13 (EHI_001050), Gp210 (EHI_183510), Nup205 (EHI_098830), Nup155 (EHI_138330), Nup93 (EHI_021460), Nup54 (EHI_010010), and Nup58 (EHI_00670) were detected (17). Most of these proteins were recovered in all three biological replicates, although their relative abundance compared with the mock control varied. Thus, rather than representing uniformly enriched interactors, this broader dataset provides evidence that HA-Nup98-like co-purifies with multiple proteins predicted to occupy different functional or architectural regions of the NPC.

**Table 3.**
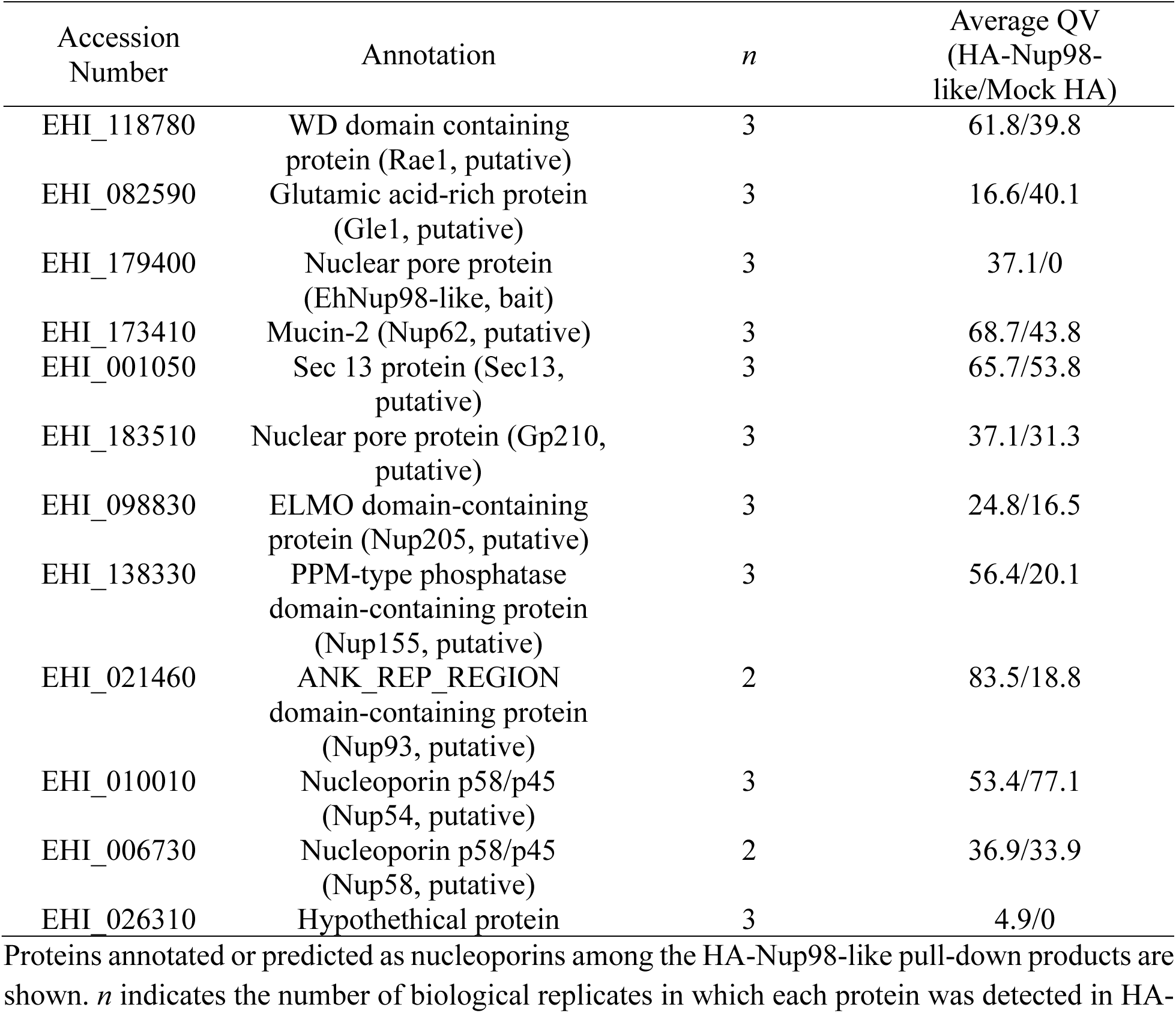
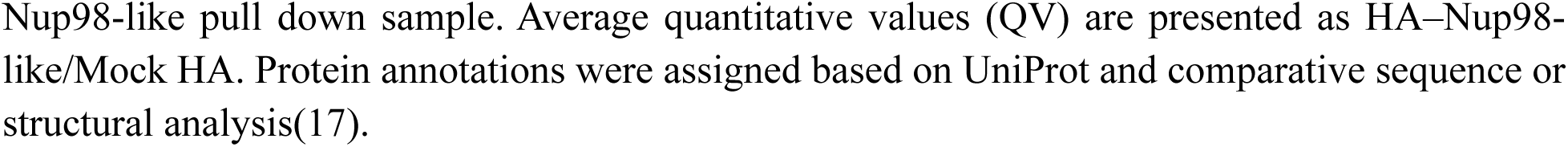
Putative nucleoporins co-immunoprecipitated with HA-Nup98-like.

| Accession Number | Annotation | *n* | Average QV (HA-Nup98-like/Mock HA) |
| --- | --- | --- | --- |
| EHI_118780 | WD domain containing protein (Rae1, putative) | 3 | 61.8/39.8 |
| EHI_082590 | Glutamic acid-rich protein (Gle1, putative) | 3 | 16.6/40.1 |
| EHI_179400 | Nuclear pore protein (EhNup98-like, bait) | 3 | 37.1/0 |
| EHI_173410 | Mucin-2 (Nup62, putative) | 3 | 68.7/43.8 |
| EHI_001050 | Sec 13 protein (Sec13, putative) | 3 | 65.7/53.8 |
| EHI_183510 | Nuclear pore protein (Gp210, putative) | 3 | 37.1/31.3 |
| EHI_098830 | ELMO domain-containing protein (Nup205, putative) | 3 | 24.8/16.5 |
| EHI_138330 | PPM-type phosphatase domain-containing protein (Nup155, putative) | 3 | 56.4/20.1 |
| EHI_021460 | ANK_REP_REGION domain-containing protein (Nup93, putative) | 2 | 83.5/18.8 |
| EHI_010010 | Nucleoporin p58/p45 (Nup54, putative) | 3 | 53.4/77.1 |
| EHI_006730 | Nucleoporin p58/p45 (Nup58, putative) | 2 | 36.9/33.9 |
| EHI_026310 | Hypothetical protein | 3 | 4.9/0 |

Proteins annotated or predicted as nucleoporins among the HA-Nup98-like pull-down products are shown. *n* indicates the number of biological replicates in which each protein was detected in HA-
Nup98-like pull down sample. Average quantitative values (QV) are presented as HA–Nup98-like/Mock HA. Protein annotations were assigned based on UniProt and comparative sequence or structural analysis(17).

Within the stringent exclusive dataset, several components of the prefoldin chaperone machinery were also reproducibly recovered, together with a putative importin β-family protein (EHI_098400) (Table 1). One previously uncharacterized protein (EHI_026310) was also detected exclusively in HA-Nup98-like. Since its lacked functional annotation in AmoebaDB, yet its reproducible and specific association with EhNup98-like prompted us to prioritize EHI_026310 for further characterization. Meanwhile, the enriched dataset additionally contained proteins with TPR- and HEAT-repeat domains and components associated with the CCR4-NOT complex (Table 2), suggesting that the EhNup98-like proteome extends beyond structural NPC components to proteins implicated in nuclear transport, RNA metabolism, and gene regulation A similarly broad functional association of NPCs with RNA processing and gene regulation has been observed in opisthokonts and divergent protists, including trypanosomes and *Plasmodium*(21–23).

To examine the broader functional composition of the EhNup98-like proteome, we performed Gene Ontology enrichment analysis using Biological Process annotations. Protein folding showed the strongest enrichment, accompanied by enrichment of processes related to protein maturation (Fig 4E). Several processes associated with RNA and gene regulation of RNA biosynthesis, regulation of DNA-templated transcription, regulation of RNA metabolic processes, DNA-templated transcription, and regulation of gene expression were enriched. Additional enriched categories encompassed broader cellular and macromolecular biosynthetic and metabolic processes.

### EHI_026310 retains Nup53/Nup35-like structural features despite extensive evolutionary divergence

Among the high-confidence proteins associated with EhNup98-like, we focused on EHI_026310, a previously uncharacterized protein that was reproducibly detected in all three replicates and was absent from the corresponding mock controls (Table 1). Despite its reproducible association with EhNup98-like, EHI_026310 had not been identified as a candidate nucleoporin in a previous profile hidden Markov model-based survey of the *E. histolytica* NPC(17). We therefore investigated whether EHI_026310 might represent a highly divergent NPC component that is poorly recognizable by sequence-based approaches.

Initial domain analysis identified a predicted RNA recognition motif (RRM)-like domain within the C-terminal region of EHI_026310. Because an RRM domain is a characteristic structural feature of Nup53/Nup35-family nucleoporins, we compared the domain architecture of EHI_026310 with those of human Nup35 (HsNup35) and *S. cerevisiae* Nup53 (ScNup53) (Figure 5A). Canonical Nup53/Nup35 proteins contain an RRM domain together with several defined interaction regions, including R1-R3 and a C-terminal membrane-interacting region (M), as in HsNup35, although ScNup53 lacks recognizable R1-R3. In contrast, EHI_026310 displayed a markedly different domain organization. Recognizable counterparts of the R1-R3 regions and M were not apparent, whereas a prominent repeat-rich region occupied much of the N-terminal portion of the protein and a predicted RRM-like domain was retained near the C-terminus (Fig 5A). Based on its reproducible association with EhNup98-ike and the presence of this Nup53/Nup35-associated structural feature, we provisionally designated EHI_026310 as EhNup53-like.

**Fig 5.**
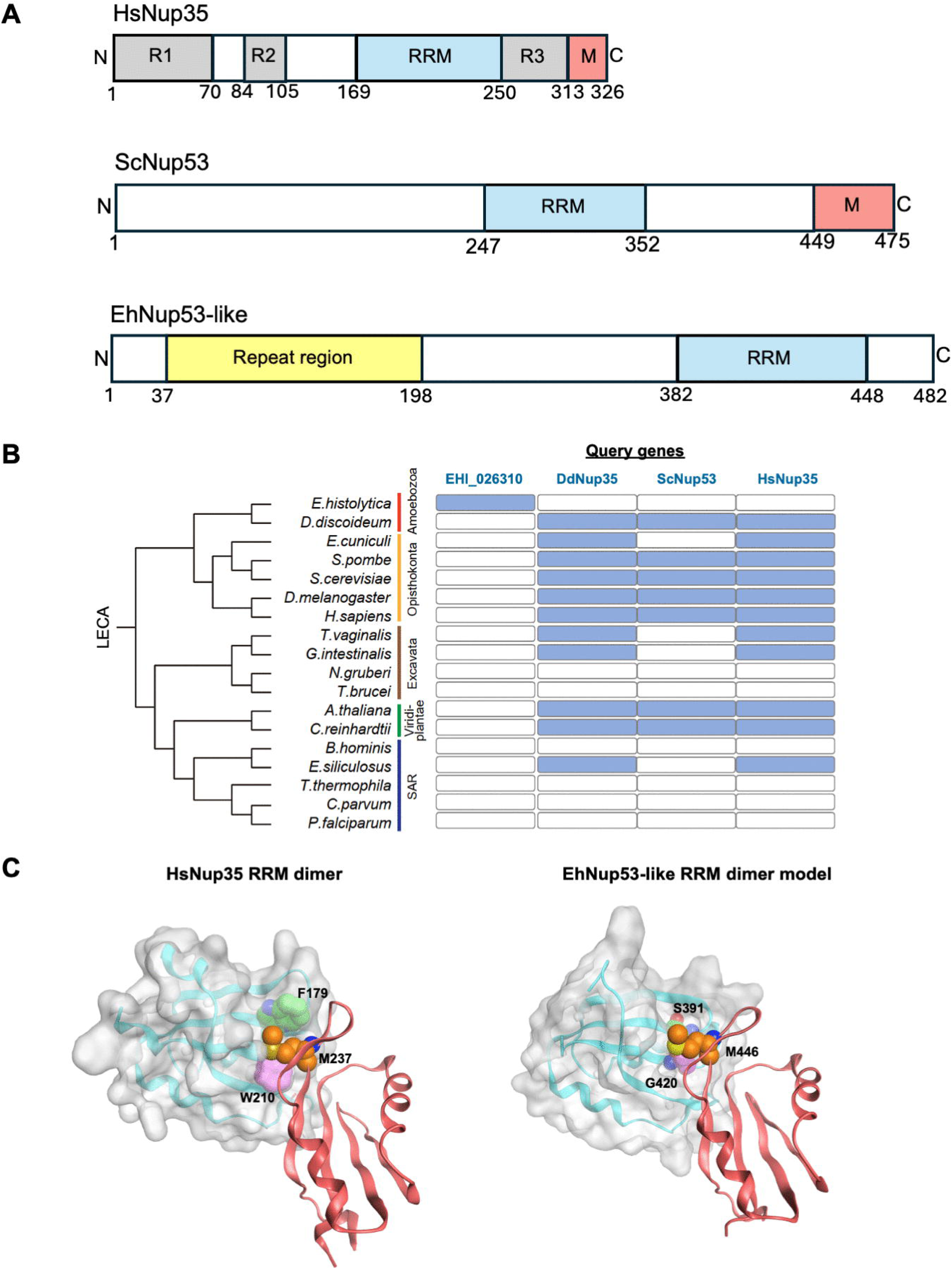
Structural and evolutionary characterization of the divergent amoebic nucleoporin EhNup53-like. **(A)** Schematic comparison of the domain architectures of human Nup35 (HsNup35), *Saccharomyces cerevisae* Nup53 (ScNup53), and *E. histolytica* Nup53-like (EhNup53-like; EHI_026310). R1-R3, interaction regions; RRM, RNA recognition motif; M, membrane-interacting region. **(B)** Profile hidden Markov model (HMM)-based remote homology analysis comparing EhNup53-like with representative Nup53/Nup35 proteins from diverse eukaryotic lineages. Blue shading indicates detectable homology to the indicated query, respectively. **(C)** Structural comparison of the RRM-like region of EhNup53-like with experimentally determined human Nup35 (PDB ID: 4LIR). The RRM dimer of human Nup35 and *E. histolytica* Nup53-like are shown in surface and ribbon representations. The RRM dimer structure of Eh Nup53-like was constructed by homology modeling based on the RRM dimer of human Nup35. Residues involved in the key hydrophobic interaction essential for RRM dimerization are shown as sphere models. In EhNup53-like, Phe and Trp residues that form the aromatic pocket are replaced by Ser and Gly residues, respectively.

We next examined whether EhNup53-like could be linked to canonical Nup53/Nup35 proteins by profile-based sequence comparisons across representative eukaryotic lineages. Cross-profile searches using RRM domains of EhNup53-like demonstrated that canonical Nup53/Nup35 proteins from *Dictyostelium discoideum*, *S. cerevisiae*, and *H. sapiens* revealed little or no detectable reciprocal sequence similarity to EhNup53-like under the applied search criteria (Fig5B). In contrast, relationships among several established Nup53/Nup35 proteins from other eukaryotic lineages were readily detected. These results indicate that EhNup53-like has undergone extensive primary-sequence divergence, sufficient to obscure its relationship with canonical Nup53/Nup35 proteins in conventional profile-based searches.

Conservation within the *Entamoeba* lineage was examined separately by multiple sequence alignment of EhNup53-like-related proteins from representative *Entamoeba* species (S3 Fig). Homologs from *E. histolytica, E. dispar* and *E. nuttalli* showed substantial conservation, particularly within the C-terminal region encompassing the predicted RRM-like domain. In contrast, the N-terminal repeat-rich region was considerably more variable in both sequence and repeat organization, with *E. invadens* and *E. moshkovskii* showing marked divergence from the more closely conserved patterns observed in *E. histolytica*, *E. dispar*, and *E. nuttalli*.

Since sequence-based comparisons provided limited evidence for a relationship with canonical Nup53/Nup35 proteins, we next examined the predicted 3D structure of RRM in EhNup53-like and asked whether the retained RRM-like module itself is the preserved structural feature characteristic of canonical Nup53/Nup35 proteins. Structural comparison with experimentally determined human Nup53/Nup35 RRM structures revealed conservation of the overall RRM-like fold but substantial differences in surface properties (Fig 5C). Canonical Nup53/Nup35 proteins contain a key hydrophobic interaction essential for RRM-mediated dimerization, in which a methionine residue is inserted into an aromatic pocket formed by phenylalanine and tryptophan residues(24). Although the corresponding region of predicted EhNup53-like retained the general fold, an equivalent aromatic pocket was not readily apparent. The EhNup53-like RRM appears to preserve the underlying structural framework while exhibiting substantial remodeling of a surface implicated in intermolecular interactions in canonical Nup53/Nup35 proteins.

### EhNup53-like localizes to the nuclear periphery and associates with nuclear pore structures

To validate the expression and subcellular localization of the candidate interactor identified in the HA-EhNup98-like pulldown, we generated a transgenic *E. histolytica* line expressing N-terminal HA-tagged EhNup53-like (HA-Nup53-like). Immunoblot analysis using an anti-HA antibody detected a prominent band at approximately 54 kDa, consistent with the predicted molecular weight (Fig 6A).

**Fig 6.**
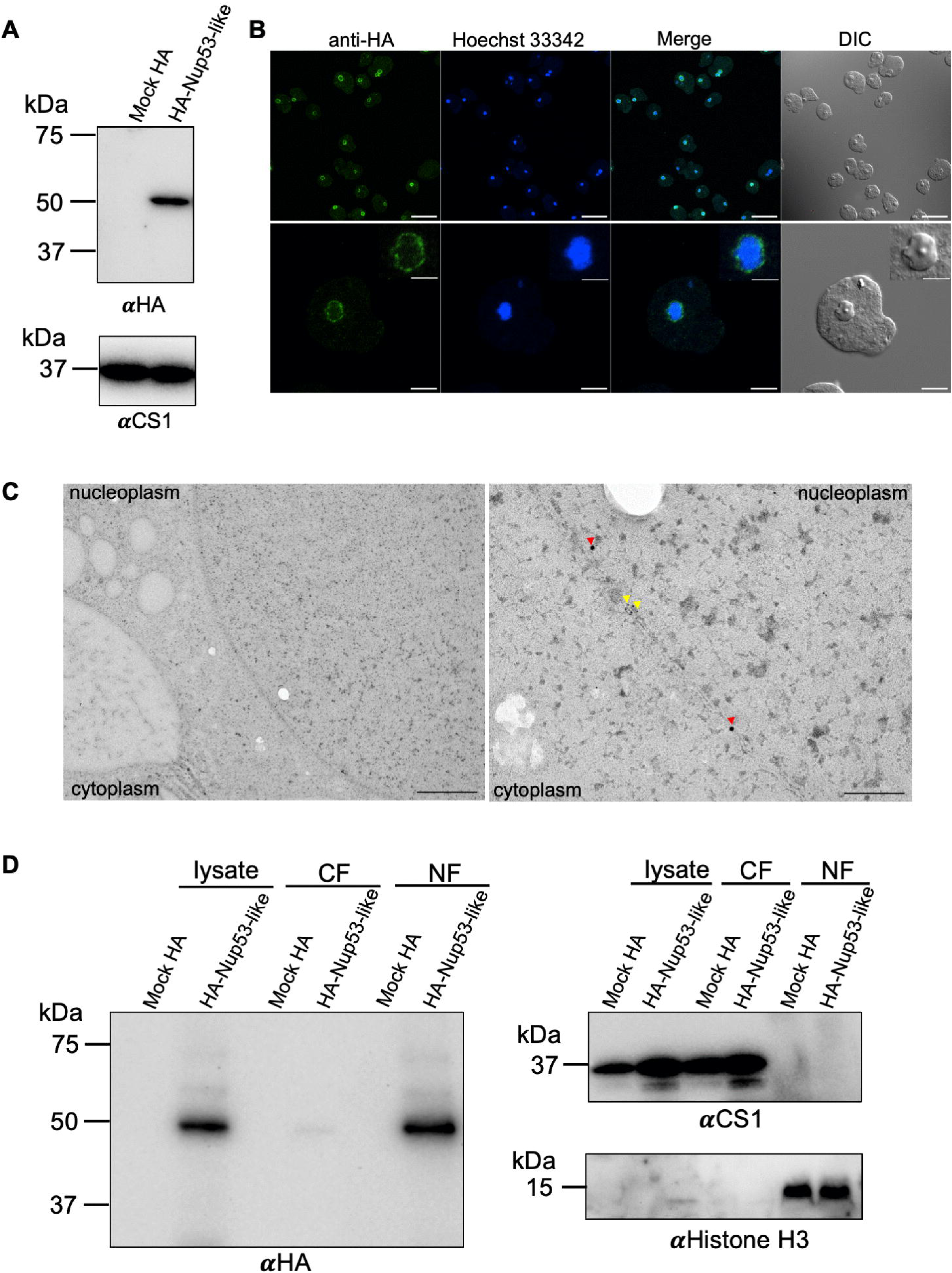
Localization of HA-Nup53-like to the nuclear membrane. **(A)** Immunoblot analysis of *E. histolytica* lysates expressing HA-Nup53-like. Anti-HA detection reveals a band at ∼54 kDa, consistent with the predicted molecular weight. CS1 was used as a loading control **(B)** Indirect immunofluorescence microscopy of HA-Nup53-like. The protein localizes predominantly to the nuclear periphery (anti-HA, green), with nuclei visualized by Hoechst 33342 (blue). Scale bar, 40 µm in the overview images, 10 µm in the enlarged images, and 4 µm in the insets. DIC, differential interference contrast. **(C)** Immuno-electron microscopy of HA-Nup53-like. Immuno-gold particles are enriched at the nuclear membrane and frequently localize to electron-dense regions consistent with NPC. Ultrathin sections were dual-labeled with mouse anti-HA antibody (10 nm anti-mouse gold particles; red arrowheads) detecting HA-Nup53-like and rabbit anti-laminB1 antibody (5 nm anti-rabbit gold particles; yellow arrowheads) marking the nuclear envelope. Scale bar, 1 µm in low magnification image, 100 nm in high magnification image **(D)** Subcellular fractionation of HA-Nup53-like. The protein is enriched in the nuclear fraction (NF) with minimal detection in the cytoplasmic fraction (CF). CS1 and Histone H3 serve as cytoplasmic and nuclear markers, respectively. Uncropped western blots available in S1 Raw Images.

We next examined the subcellular distribution of HA-Nup53-like by indirect immunofluorescence microscopy. Anti-HA staining revealed prominent enrichment around the nucleus, with a distinct rim-like pattern observed in approximately 85% of analyzed trophozoites (Fig 6B). To examine this localization at higher spatial resolution, we performed immuno-electron microscopy. HA-Nup53-like-associated gold particles were enriched along the nuclear membrane (Fig 6C).

To further validate these observations biochemically, we performed subcellular fractionation followed by immunoblotting. HA-Nup53-like was strongly enriched in the nuclear fraction (NF), with minimal signal detected in the cytoplasmic fraction (CF) (Fig 6D). Fraction quality was assessed using CS1 as cytoplasmic marker and histone H3 as nuclear marker. This result is consistent with the imaging data and indicate that EhNup53-like is predominantly associated with nuclear structures.

### Reciprocal proteomic analysis of EhNup53-like identifies EhNup98-like and additional candidate NPC components

To further define the interaction network of EhNup53-like and independently assess its association with the *E. histolytica* NPC, we performed reciprocal co-immunoprecipitation followed by LC-MS/MS using HA-Nup53-like as bait in three independent biological replicates (Fig 7A).

**Fig 7.**
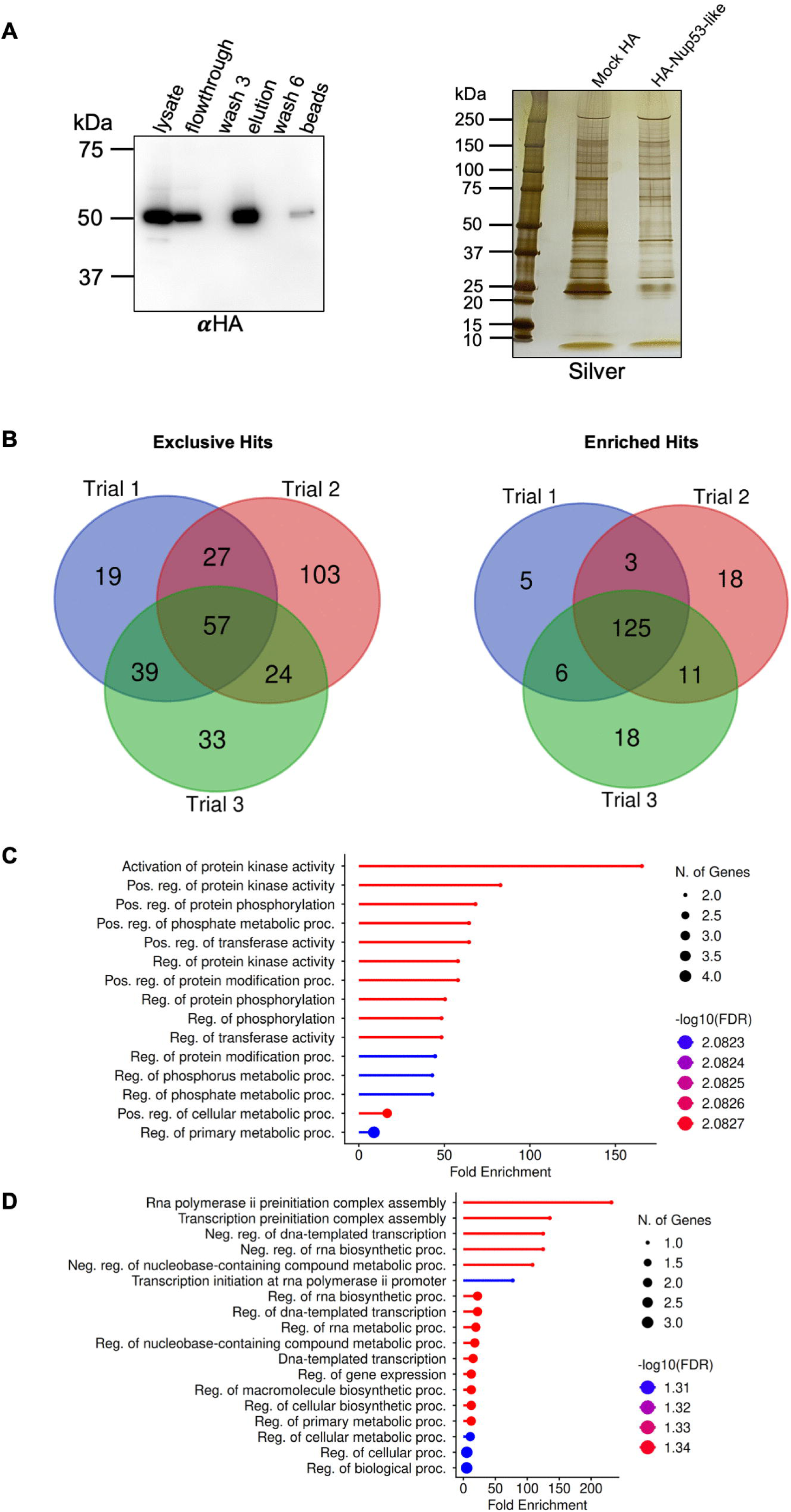
Proteomic landscape of the EhNup53-like interactome. **(A)** Validation of HA-Nup53-like co-immunoprecipitation by immunoblotting with anti-HA antibody. Eluted fractions were analyzed by SDS-PAGE followed by silver staining. Uncropped western blots available in S1 Raw Images. **(B)** Venn diagrams showing overlap of identified proteins across three independent HA-Nup53-like co-IP replicates. Left: exclusive hits (57 proteins) detected in all three replicates and absent in mock control. Right: enriched hits (125 proteins) with abundance ratio ≥ 5.0 and p-value < 0.05 across all three replicates. **(C)** Gene Ontology (GO) enrichment analysis of exclusive interactors, highlighting strong enrichment in kinase activation and phosphorylation-related processes. **(D)** GO enrichment analysis of enriched interactors, showing representation of protein degradation and macromolecular catabolic pathways.

We first identified proteins detected exclusively in HA-Nup53-like and absent from the corresponding mock controls. Although the total number of exclusive proteins varied among replicates, 57 proteins were reproducibly detected in all three experiments (Fig 7B, left; Table 4; the remaining proteins are listed in S1 Table).

**Table 4.** Proteins identified as exclusive interactors of HA-Nup53-like by immunoprecipitation–mass spectrometry.

| Accession Number | Annotation | Average Abundances<br>HA-Nup53-like | p-value |
| --- | --- | --- | --- |
| EHI_026310 | Hypothetical protein (Nup53-like) | 4.E+09 | 0.000 |
| EHI_147500 | Serine/threonine-protein phosphatase | 6.E+06 | 0.049 |
| EHI_131510 | Ras modification protein ERF4 | 4.E+06 | 0.014 |
| EHI_092530 | Protein kinase domain containing<br>protein | 3.E+06 | 0.048 |
| EHI_133920 | TPR repeat protein | 3.E+06 | 0.002 |
| EHI_082900 | Uncharacterized protein | 2.E+06 | 0.032 |
| EHI_159570 | Mob1/phocein family protein | 2.E+06 | 0.003 |
| EHI_151800 | Ganglioside activator protein,<br>putative | 2.E+06 | 0.047 |
| EHI_133100 | Rab family GTPase | 2.E+06 | 0.042 |
| EHI_098010 | WD_REPEATS_REGION domain-containing protein | 2.E+06 | 0.047 |
| EHI_140240 | Deoxycytidine triphosphate<br>deaminase, putative | 2.E+06 | 0.001 |
| EHI_111790 | LIM zinc finger domain containing<br>protein | 2.E+06 | 0.018 |
| EHI_143110 | Acetyltransferase, putative | 1.E+06 | 0.008 |
| EHI_045310 | PH domain-containing protein | 1.E+06 | 0.019 |
| EHI_126890 | Transmembrane protein | 1.E+06 | 0.011 |
| EHI_129340 | FHA domain-containing protein | 1.E+06 | 0.010 |
| EHI_065830 | Density-regulated protein, putative | 1.E+06 | 0.043 |
| EHI_151310 | LSM domain containing protein | 1.E+06 | 0.027 |

Proteins were identified by immunoprecipitation–mass spectrometry (IP–MS). High-confidence interactors were defined as proteins detected in HA-Nup53-like samples with a $p$ -value $< 0.05$ and an average abundance $\geq 1.0 \times 10^6$ . Average abundance represents normalized signal intensity in HA-Nup53-like samples across three biological replicates. Proteins are ranked by decreasing abundance. Functional annotations are based on UniProt.

We next examined proteins detected in both HA-Nup53-like and mock samples but enriched in the HA-Nup53-like samples. Using an abundance ratio of ≥5.0 and a *p*-value <0.05 as selection criteria, 125 proteins were consistently enriched across all three biological replicates (Fig 7B, right; Table 5; the remaining proteins are listed in S2 Table). Notably, EhNup98-like (EHI_179400), which had initially identified EhNup53-like, was reproducibly recovered in the HA-Nup53-like reciprocal pull-down dataset with an abundance ratio of 5.8. This reciprocal recovery independently supports the association between EhNup98-like and EhNup53-like.

**Table 5.**
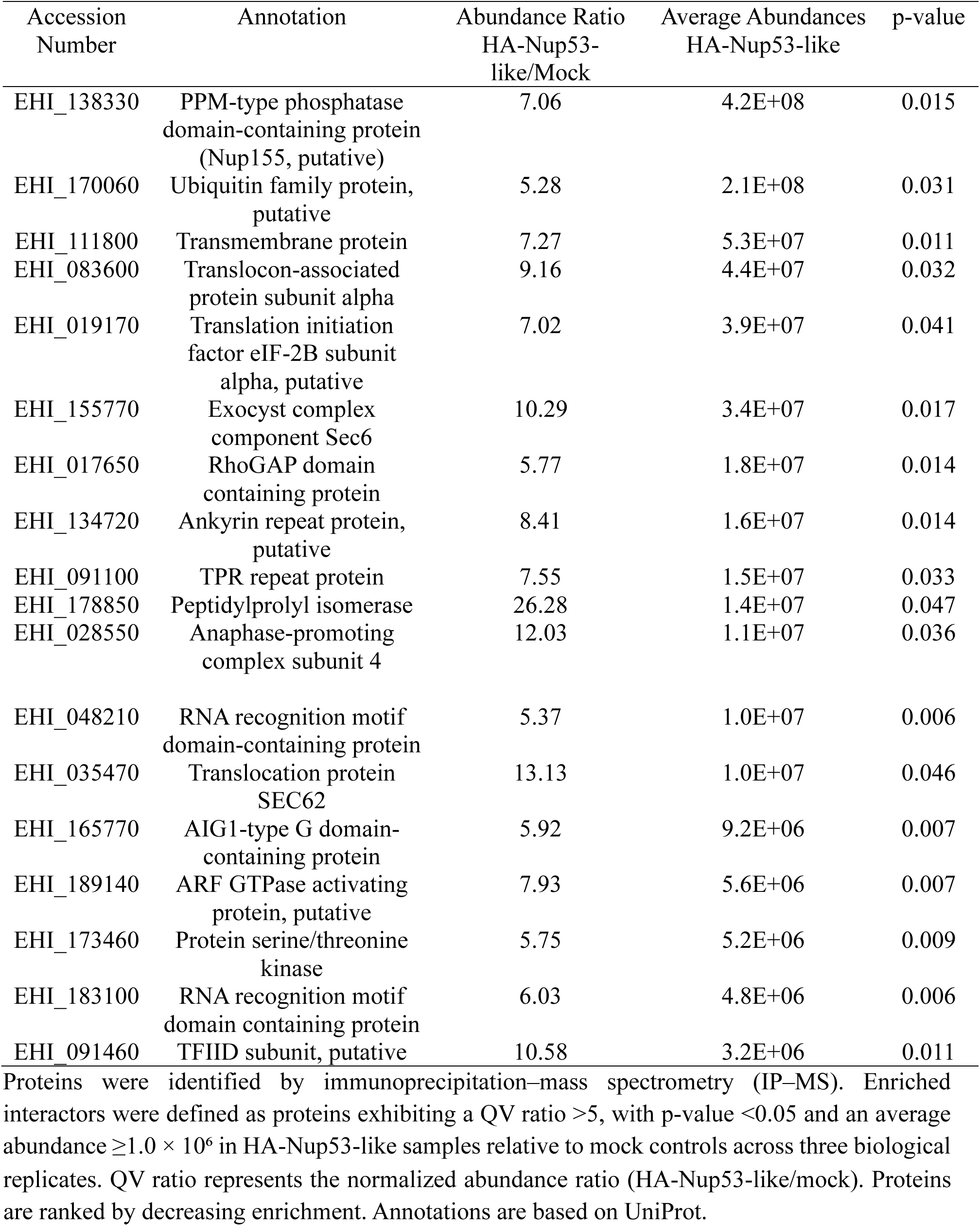
Proteins identified as enriched interactors of HA-Nup53-like by immunoprecipitation–mass spectrometry.

Because individual NPC-associated proteins may not necessarily satisfy the enrichment threshold, we additionally surveyed the complete HA-Nup53-like proteomic dataset for proteins annotated or predicted to function as nucleoporins or NPC-associated factors. This analysis recovered multiple candidate components of the *E. histolytica* nucleoporins (Table 6). In addition to EhNup98-like, proteins assigned as putative Rae1 (EHI_118780), Gle1 (EHI_082590), Nup116 (EHI_193440), Nup100 (EHI_108620), Nup62 (EHI_173410), Sec13 (EHI_001050), Gp210 (EHI_183510), Nup205 (EHI_098830), Nup155 (EHI_138330), Nup93 (EHI_021460), Nup54 (EHI_010010), and Nup58 (EHI_006730) were recovered. Several additional proteins annotated more generally as nucleoporins or nuclear pore complex proteins were also detected (Table 6). Most of these candidates were observed in all three biological replicates, although their abundance relative to the mock control varied.

**Table 6.**
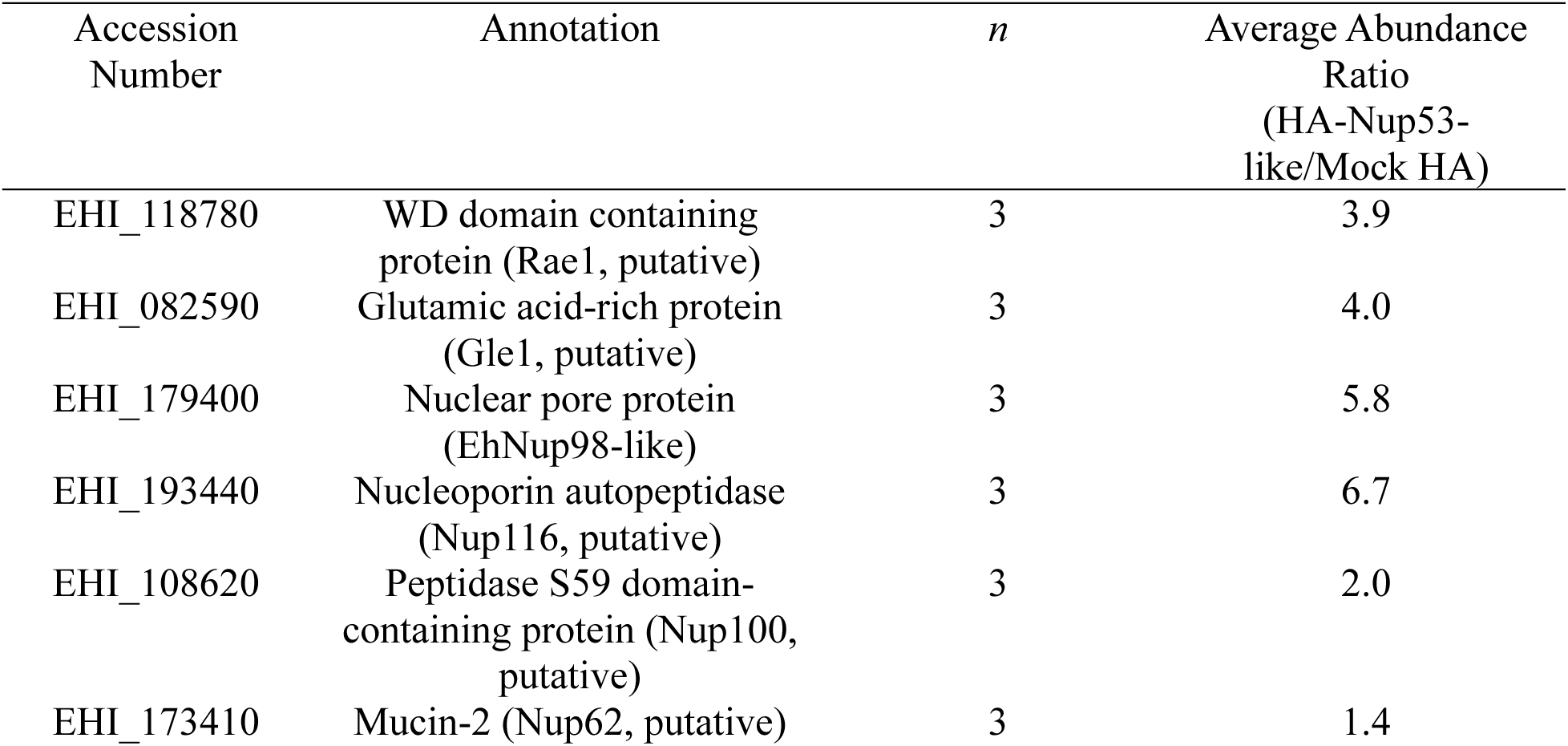
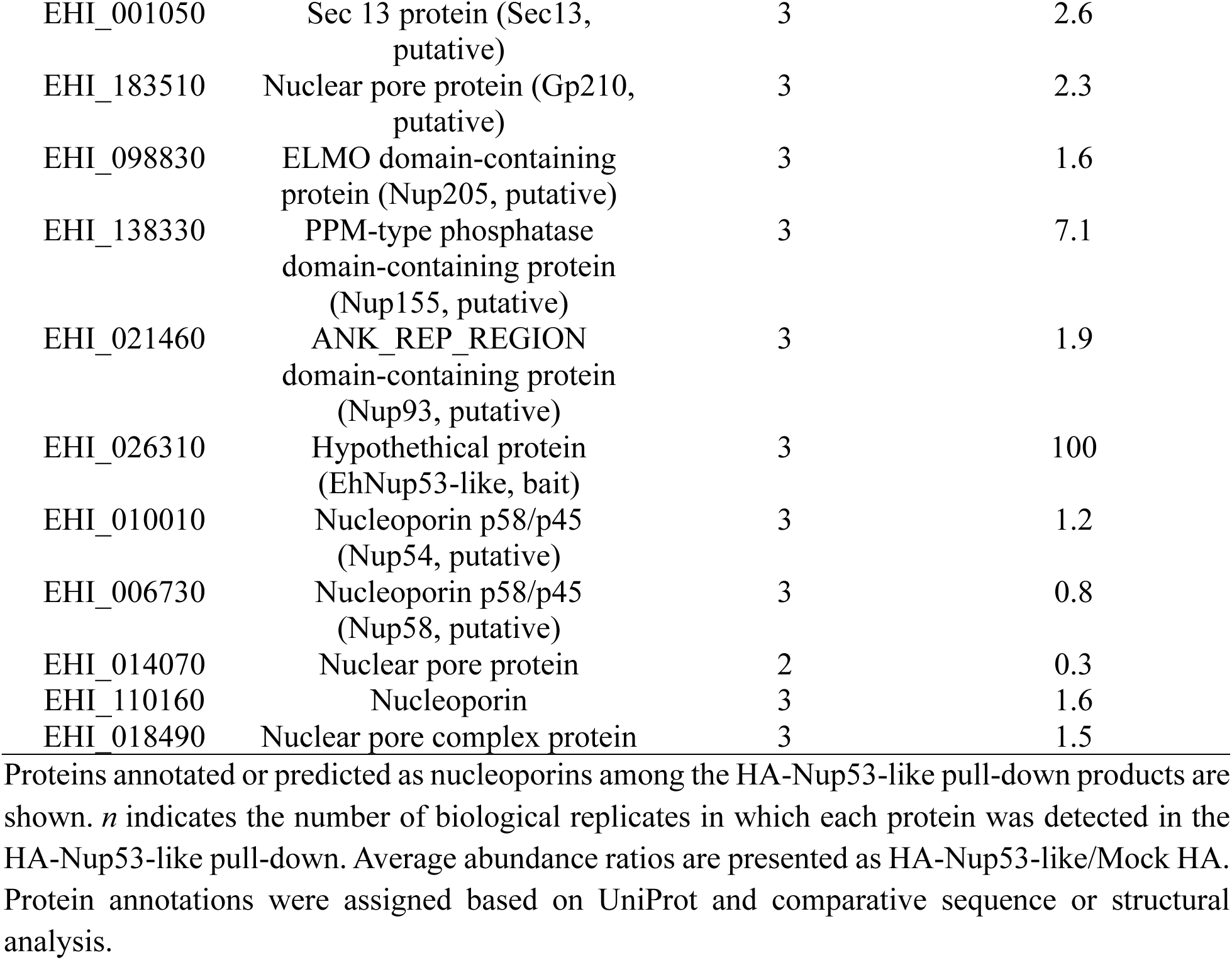
Putative nucleoporins co-immunoprecipitated with HA-Nup53-like.

Among these proteins, the putative Nup155 (EHI_138330; EhNup155-like) was enriched approximately 7.1-fold relative to the mock control and was detected in all three replicates (Table 5). Given the conserved interaction between Nup53/Nup35 and Nup155/170 in other eukaryotes(25), we further examined the domain organization of EhNup155-like. In contrast to human Nup155, no recognizable conserved domains were detected in EhNup155-like (S4 Fig). EhNup98-like was similarly recovered in all three replicates with an abundance ratio of approximately 5.8. A putative Nup116 protein (EHI_193440) was also enriched approximately 6.7-fold. Other candidate NPC components, including putative Nup205, Nup93, Nup54, Nup58, Nup62, Sec13, and Gp210 proteins, were reproducibly detected but showed more modest enrichment (Table 6).

To characterize the broader functional composition of the EhNup53-like proteome, we performed Gene Ontology enrichment analysis using Biological Process annotations. Analysis of the reproducible exclusive dataset revealed enrichment of biological processes associated with protein kinase regulation and phosphorylation-related pathways (Fig 7C). Gene Ontology analysis of the enriched dataset identified an additional set of Biological Process categories related to protein catabolism and proteolysis, including processes associated with proteasome-dependent and ubiquitin-related protein degradation (Fig 7D). Together, the two proteomic datasets indicate that proteins associated with EhNup53-like extend beyond candidate NPC components and encompass proteins participating in phosphorylation-related regulation and cellular protein homeostasis.

### The RRM-like domain contributes to nuclear-peripheral localization of EhNup53-like

To examine the functional significance of the major predicted regions of EhNup53-like, we generated HA-tagged deletion constructs lacking either the N-terminal repeat-rich region (HA-Nup53-like^Δ37–198^) or the predicted RRM-like domain (HA-Nup53-like^Δ382–448^). Immunoblotting with an anti-HA antibody confirmed expression of both deletion constructs at molecular masses consistent with their predicted sizes (Fig 8A). We then examined the localization of the deletion variants by IFA. Deletion of the N-terminal repeat-rich region did not abolish the pattern of rim-like enrichment around the nucleus, observed for wild-type HA-Nup53 like, and HA-Nup53-like^Δ37–198^ remained enriched at the nuclear periphery (Fig 8B). In contrast, deletion of the predicted RRM-like domain markedly altered the distribution of EhNup53-like. HA-Nup53-like^Δ382–448^ no longer displayed the distinct nuclear-rim enrichment and instead showed a more diffuse intracellular distribution accompanied by punctate accumulations (Fig 8B). These observations indicate that the repeat-rich region dispensable for nuclear-peripheral localization of EhNup53-like whereas the RRM-like domain is essential for it.

**Fig 8.**
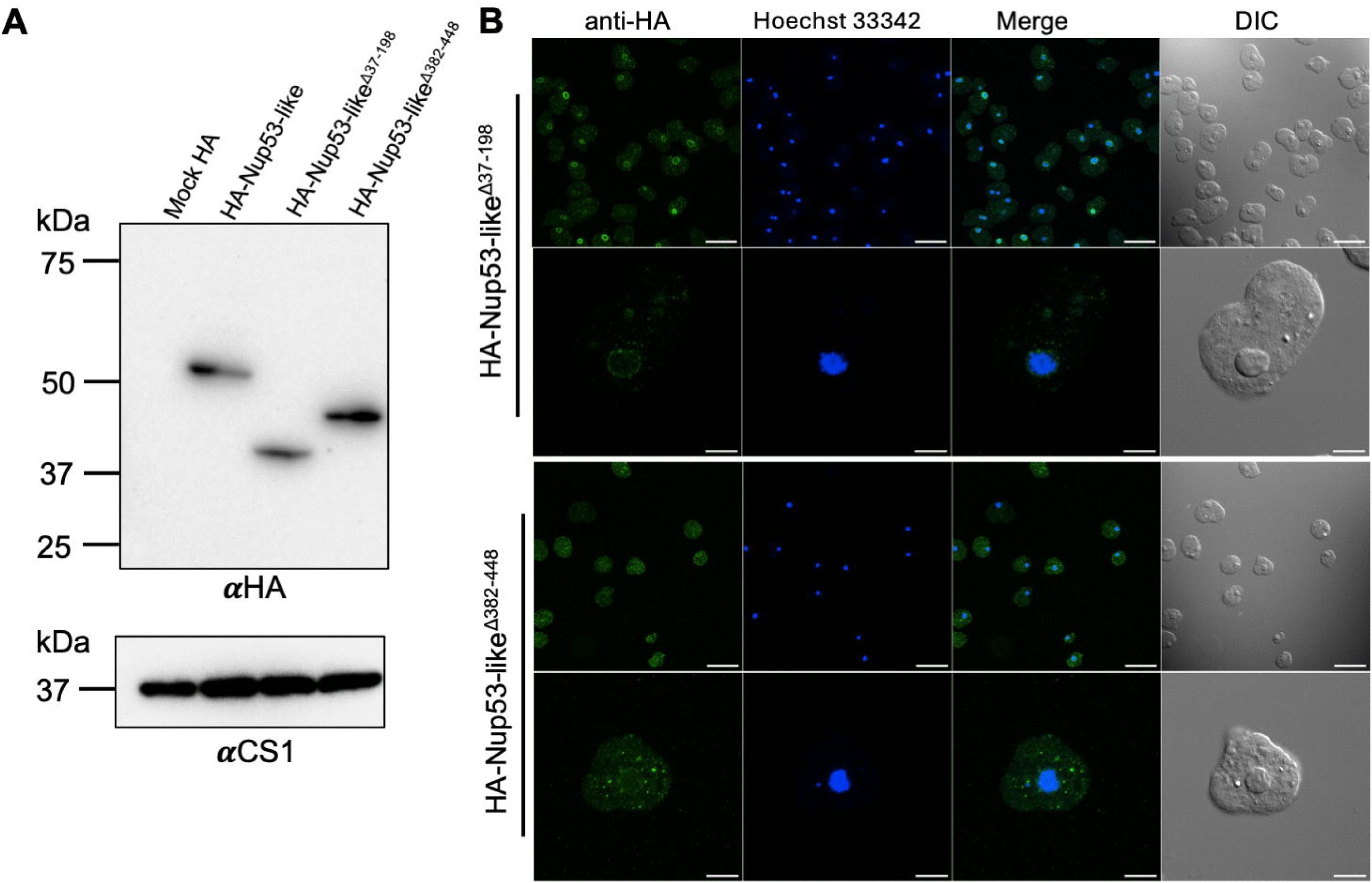
Domain-dependent localization of EhNup53-like revealed by targeted mutation. **(A)** Expression and localization of HA-Nup53-like and deletion mutants lacking the repeat region (HA-Nup53-like^Δ37-198^) or RRM domain (HA-Nup53-like^Δ382-488^). Immunoblot analysis of HA-tagged proteins using anti-HA antibody, with CS1 as a loading control. Uncropped western blots available in S1 Raw Images. **(B)** Immunofluorescence microscopy showing the localization patterns of wild-type and mutant proteins. HA signal (green), Hoechst 33342 nuclear staining (blue), and DIC images are shown. Scale bar, 40 µm in the overview images, 10 µm in the enlarged images.

To determine whether disruption of the major domains of EhNup53-like affects mRNA export, we examined the intracellular distribution of poly(A)^+^ RNA in trophozoites expressing HA-Nup53-like, HA-Nup53-like^Δ37–198^, or HA-Nup53-like^Δ382–448^. RNA FISH was performed using FITC-conjugated oligo(dT)_30_ together with immunofluorescence detection of the HA-tagged proteins (S5A Fig). Poly(A)^+^ RNA was distributed throughout the cytoplasm and nucleus in cells expressing HA-Nup53-like. A comparable distribution was observed in cells expressing either the repeat-region deletion mutant HA-Nup53-like^Δ37–198^ or the RRM-deletion mutant HA-Nup53-like^Δ382–448^, despite the altered subcellular localization of the latter.

Quantification of the nuclear-to-cytoplasmic (N/C) poly(A)^+^ RNA fluorescence ratio showed similar values among HA-Nup53-like and both deletion mutants, with no significant differences detected between the groups (One way ANOVA, p = 0.5884; S5B Fig). Thus, deletion of either the N-terminal repeat-rich region or the C-terminal RRM-like domain of HA-Nup53-like did not produce detectable nuclear accumulation of bulk poly(A)^+^ RNA under the conditions examined.

## Discussion

The NPC is an ancient eukaryotic assembly whose core organizational features are broadly conserved despite extensive divergence in the sequences and domain architectures of individual nucleoporins(26,27). Our study extends this evolutionary framework to Amoebozoa by characterizing an FG-repeat Nup98-like protein and identifying a previously unrecognized, highly divergent RRM-containing Nup53-like protein in *Entamoeba histolytica*. Together, the localization, biochemical, proteomic, and domain-deletion data support a model in which the amoebic NPC retains recognizable features of canonical NPC organization, including an FG-rich transport-barrier component connected to an inner-ring-like scaffold, while the proteins implementing these roles have undergone substantial lineage-specific remodeling.

### EhNup98-like retains core FG-nucleoporin features despite lineage-specific remodeling

EhNup98-like preserves two defining properties of canonical Nup98-family proteins: an N-terminal low-complexity FG-repeat region and a C-terminal peptidase S59 domain(28–30). However, its FG-repeat region is reduced and differently organized compared with well-characterized opisthokont Nup98 proteins. Remodeling of FG-nucleoporins has similarly been reported in other divergent eukaryotes, including trypanosomes, in which the NPC retains the basic classes of scaffold and FG-repeat proteins despite marked differences in individual components(26). EhNup98-like is therefore consistent with a broader evolutionary pattern in which the properties of FG-rich, intrinsically disordered regions can be retained despite considerable variation in repeat number and primary sequence. Whether the reduced FG content of EhNup98-like is compensated or complemented by other FG- or low-complexity nucleoporins remains unknown, but these differences suggest that the permeability barrier of the *E. histolytica* NPC may differ compositionally from that of opisthokonts.

The peptidase S59 region provides a second point of conservation. EhNup98-like contains an HFT sequence at the predicted autoproteolytic site rather than the HFS motif commonly described in vertebrate Nup98, while otherwise retaining the predicted catalytic architecture of the domain(28,31–33). Immunoblotting consistently detected a predominant band of approximately 37 kDa together with a weaker full-length band of approximately 42 kDa, a pattern compatible with proteolytic processing of EhNup98-like. The conservation of this feature is notable because autoproteolytic activity is not universal among divergent protist Nup98-family proteins. In *Trypanosoma brucei*, the Nup98/Nup96-related protein retains the corresponding structural module but lacks catalytic residues required for cleavage(26). Thus, *E. histolytica* appears to have retained a Nup98-like processing mechanism despite substantial divergence elsewhere in the protein. Because cleavage was inferred from migration behavior rather than directly mapped, its precise site and functional consequences remain to be established.

Multiple independent observations support incorporation of EhNup98-like into nuclear pore-associated assemblies. Immunofluorescence microscopy, nuclear fractionation, and immuno-electron microscopy consistently localized EhNup98-like predominantly at the nuclear periphery, while blue-native PAGE and size-exclusion chromatography detected the protein in multiple higher-molecular-mass species. These native complexes are most conservatively interpreted as evidence that EhNup98-like participates in heterogeneous native protein assemblies rather than as proof of particular stoichiometric NPC subcomplexes. The cytosolic pool detected by imaging and fractionation is also consistent with the comparatively dynamic behavior of FG-nucleoporins, although mobility and exchange at the pore were not measured directly in *E. histolytica*(34–37).

Functional perturbation further supports a conserved role for EhNup98-like in nucleocytoplasmic transport. Silencing of EhNup98-like reduced trophozoite proliferation and increased nuclear accumulation of poly(A)^+^ RNA relative to control cells, consistent with impaired mRNA export. This phenotype agrees with studies in vertebrate cells in which perturbation or depletion of Nup98 results in nuclear retention of poly(A)^+^ RNA, supporting a conserved contribution of Nup98-family nucleoporins to RNA export(14,38,39). In the divergent apicomplexan parasite *Toxoplasma gondii*, depletion of the Nup98/Nup96-related TgNup302 similarly causes a pronounced proliferation defect and disrupts nuclear transport, although its reported RNA-export phenotype predominantly affects 18S rRNA rather than bulk poly(A)^+^ RNA(40). These observations suggest that despite substantial divergence in its FG-repeat organization, EhNup98-like retains a physiologically important role in nuclear transport. The accompanying growth defect may therefore reflect broader consequences of impaired nucleocytoplasmic exchange and gene expression, although the present data do not establish that defective mRNA export alone is responsible for reduced proliferation.

The EhNup98-like proteome further places the protein within an expected nuclear transport and NPC-associated molecular environment. Recovery of an importin-β-like protein is consistent with the canonical interaction of transport receptors with FG-repeat domains(41–44). More importantly for NPC architecture, EhNup53-like was reproducibly recovered with EhNup98-like. This association should not be taken to establish a conserved direct Nup98–Nup53 interaction, but it links the amoebic FG-nucleoporin to a candidate inner-ring scaffold protein and therefore supports preservation of the broader architectural principle by which the permeability barrier is anchored to the NPC scaffold. Other enriched proteins, including prefoldin, CCR4–NOT components, and ubiquitin-related factors, indicate that EhNup98-like resides in a broader protein environment connected to protein homeostasis and RNA regulation(45–50). These associations are potentially informative but their relationships to the NPC remain to be determined.

### EhNup53-like illustrates evolutionary remodeling of the NPC inner ring

EhNup53-like provides a more striking example of NPC divergence. Canonical Nup53/Nup35 proteins contain an RRM-like domain together with several interaction segments that connect them to inner-ring partners such as Nup93/Nic96, Nup155/Nup170, and Nup205/Nup192(25). EhNup53-like lacks detectable overall sequence homology to canonical Nup53 proteins and lacks several recognizable interaction motifs found in opisthokont homologues, yet structural prediction identifies an RRM-like fold characteristic of this nucleoporin family. Combined with its localization and interaction profile, these features support its assignment as a highly divergent Nup53-like NPC component despite the absence of conventional sequence-based evidence for orthology. Although its precise evolutionary relationship to canonical Nup53/Nup35 proteins cannot be resolved from the present data, EhNup53-like is consistent with extensive divergence from an ancestral inner ring component.

The contrasting conservation of structured and low-complexity regions provides further insight into this divergence. Comparative analysis of known Nup53 orthologs indicates that the RRM-containing core is more conserved than several peripheral partner-binding regions, and the corresponding region is also the most recognizable structured feature of EhNup53-like. In contrast, the N-terminal portion of EhNup53-like contains an expanded glycine/asparagine-rich low-complexity repeat region that is not comparably developed in other examined *Entamoeba* species. Such a pattern is consistent with the greater evolutionary plasticity of intrinsically disordered regions relative to structured domains, which are generally subject to stronger constraints on sequence and fold conservation(51). The deletion data revealed that removal of the N-terminal repeat region did not abolish nuclear-rim localization, whereas deletion of the predicted RRM-containing region caused loss of normal peripheral localization and prominent cytoplasmic aggregation. These results indicate that the repeat expansion is dispensable for targeting under the conditions tested, whereas the RRM-containing region is required for normal localization and/or protein behavior. Because the ΔRRM protein aggregates, however, the phenotype cannot yet be assigned specifically to loss of an NPC-targeting interface rather than altered folding or stability.

Despite the pronounced localization defect of the ΔRRM mutant, neither deletion caused a detectable increase in the nuclear-to-cytoplasmic ratio of poly(A)^+^ RNA. The absence of mRNA-export phenotype in the EhNup53-like mutants suggests that the major functional consequence of disrupting the RRM-containing region is not a generalized block in mRNA export. Instead, EhNup53-like may contribute primarily to NPC organization, structural stability, or protein–protein interactions. This interpretation is consistent with studies of canonical Nup53/Nup35 proteins, in which the RRM-like domain contributes to dimerization, membrane association, and NPC assembly rather than acting primarily as a conventional RNA-binding module(52,53). The requirement of the EhNup53-like RRM-containing region for proper localization therefore supports conservation of a structural role despite extensive sequence divergence. At the same time, the lack of a measurable bulk poly(A)^+^ RNA-export defect suggests that disruption of EhNup53-like does not cause transport-associated consequences like EhNup98-like depletion.

The EhNup53-like interaction network also supports an inner-ring-like assignment. EhNup53-like localizes predominantly to the nuclear rim by immunofluorescence and expansion microscopy, is enriched at the nuclear envelope by immuno-EM, and is recovered mainly in the nuclear fraction. In contrast to the broader cellular distribution of EhNup98-like, this relatively stable peripheral localization is consistent with a scaffold-associated role. Proteomic analysis additionally recovered a putative EhNup155-protein, providing an independent connection to the inner-ring architecture. Canonical Nup53/Nup35–Nup155/Nup170 interfaces are important elements of the inner ring(25,54), making recovery of EhNup155-like particularly notable. This association raises the possibility that the Nup53-Nup155 architectural relationship is retained in *E. histolytica*, although direct biochemical or structural characterization will be required to establish the molecular basis of the interaction.

Similar observations in other divergent eukaryotes indicate that the conservation of NPC function does not require preservation of every canonical interaction mechanism. In trypanosomes, inner-ring components can retain their architectural positions while using lineage-specific membrane-anchoring or partner-interaction features(9,27). *E. histolytica* may represent another example of this evolutionary mechanism, in which a conserved scaffold function is implemented through a substantially remodeled domain architecture. The identification of EhNup53-like therefore broaden the range of molecular architectures known to support inner ring organization in divergent eukaryotes.

### Distinct nucleoporin-associated proteomes reveal different molecular environments at the amoebic nuclear periphery

The EhNup98-like and EhNup53-like interactomes showed clearly different compositions. EhNup98-like preferentially recovered proteins associated with nucleocytoplasmic transport-, RNA-, and chaperone-related proteins, whereas the EhNup53-like dataset contained more phosphorylation-related factors and, among the broader set of enriched proteins, components associated with ubiquitin-dependent degradation and the proteasome. These differences are consistent with the two proteins occupying distinct molecular environments, with EhNup98-like functioning as a more dynamic FG-nucleoporin associated with transport factors and EhNup53-like as a more stably positioned scaffold-associated protein. Together, these datasets suggest that different regions of the *E. histolytica* NPC engage with partially distinct sets of cellular proteins.

NPCs in opisthokonts provide organizational platforms for processes extending beyond nucleocytoplasmic transport, including transcriptional regulation, protein quality control, SUMO/ubiquitin pathways, and genome maintenance(55–58). Whether comparable functions are associated with the *E. histolytica* NPC remains unknown. In particular, enrichment of kinase- or proteasome-related proteins does not establish catalytic functions for EhNup53-like, nor do CCR4–NOT or prefoldin associations establish direct roles for EhNup98-like in mRNA regulation or chaperone activity. The most conservative interpretation is that EhNup98-like and EhNup53-like are embedded in partially distinct protein networks at the amoebic nuclear periphery. Establishing the biological significance of these associations will require validation of selected interactions and functional analysis of their components.

### Evolutionary implications for nuclear organization in divergent eukaryotes

Taken together, our findings support a model in which the *E. histolytica* NPC is evolutionarily conservative at the level of organizational logic but highly plastic in the molecular features of individual components. EhNup98-like retains an FG-rich transport-barrier module, a conserved S59 catalytic fold, nuclear-envelope localization, and a role in nucleocytoplasmic transport. EhNup53-like represents a more extreme case of divergence, while retaining RRM-like structure, nuclear peripheral localization, and connections to candidate NPC components. These two proteins illustrate different degrees and modes of diversification within the same NPC.

This pattern is consistent with comparative studies suggesting that the basic organization of the NPC was already established in the last eukaryotic common ancestor, followed by substantial lineage-specific diversification of individual nucleoporins and architectural features(59,60). The present data extend that principle to Amoebozoa and further suggest that low-complexity regions and protein-interaction interfaces may tolerate greater evolutionary change than structured cores. In EhNup53-like, the lineage-expanded N-terminal repeat region appears dispensable for nuclear-rim targeting, whereas the conserved RRM-containing region is required for normal localization. In the putative EhNup53-like–EhNup155-like pair, divergence on both sides of a predicted scaffold interface raises the possibility of coordinated evolutionary remodeling, although establishing such co-evolution will require direct structural and comparative evidence.

Accordingly, the broader significance of the *E. histolytica* NPC extends beyond defining its molecular composition. It provides an experimentally accessible example of how an ancient macromolecular machinery accommodates extensive molecular diversification. A more complete inventory and structural map of the *E. histolytica* NPC will be required to determine how far this remodeling extends, particularly across the inner ring, membrane anchors, and transport barrier. Comparative analysis across Amoebozoa should further help distinguish features that are ancestral from those that arose specifically within *Entamoeba* lineage.

## Materials and Methods

### Plasmid Construction

To generate N-terminal HA-tagged constructs, the coding sequence of EhNup98-like and EhNup53-like genes (AmoebaDB accession number: EHI_179400 and EHI_026310, respectively) from *Entamoeba histolytica* were amplified by PCR using cDNA as template. All primer sequences are listed in S3 Table. PCR products were digested with *Xma*I and *Xho*I, purified, and ligated into the pEhEx-HA episomal expression vector pre-digested with the same restriction enzymes, generating pEhEx-HA-Nup98-like and pEhEx-HA-Nup53-like, which encode proteins fused to an N-terminal HA epitope tag.

Site-directed mutagenesis was performed using the PrimeSTAR Mutagenesis Basal Kit (Takara Bio) according to the manufacturer’s instructions. The following variants were generated: HA-Nup98-like^T296A^, targeting the autoproteolytic motif; HA-Nup53-like^Δ37-198^ construct lacking the N-terminal repeat region of; and HA-Nup53-like^Δ382-488^ construct lacking the predicted RNA recognition motif domain. All constructs were verified by Sanger sequencing.

### Cell Culture and Transfection

Trophozoites of *E. histolytica* clonal strain HM-1:IMSS cl6 were cultured axenically in Diamond’s BI-S-33 (BIS) medium at 35.5°C. Plasmids encoding HA-tagged nucleoporin, generated as described above, were introduced into trophozoites using liposome-mediated transfection(61). Stable transformants were selected and maintained in the presence of G418 (Invitrogen, USA) at a final concentration of 10 μg/ml. Expression of HA-tagged proteins was confirmed by immunoblotting as described below.

### Immunofluorescence Assays (IFA)

*E. histolytica* trophozoites were transferred onto 8 mm round well slide glass (Matsunami Glass Ind, Osaka, Japan) and allowed to adhere for 15 min at 35.5°C. Attached cells were then fixed with 4% paraformaldehyde (PFA) for 10 min at room temperature, followed by permeabilization with 0.2% Triton X-100 in PBS for 10 min and subsequent blocking in PBS containing 1% bovine serum albumin (BSA) for 10 min. Samples were incubated with mouse anti-HA monoclonal antibody (1:500 dilution) for 1 h at room temperature. After washing three times with PBS containing 0.1% BSA, cells were incubated with Alexa Fluor-488 conjugated anti-mouse IgG secondary antibody (1:1,000 dilution) and Hoechst 33342 (1:5,000 dilution) for nuclear staining. Images were acquired using a Nikon Ti2 AX/AX-R confocal microscope and analyzed with NIS-Elements software. Final image processing was performed using Fiji (ImageJ)(62).

### Poly(A)^+^ RNA fluorescence in situ hybridization

Poly(A)^+^ RNA fluorescence in situ hybridization (RNA FISH) was performed based on López-Rosas et al., with modifications(63). Cells were fixed with 4% paraformaldehyde (PFA), washed with phosphate-buffered saline (PBS), and incubated with 25 mM NH_4_Cl for 10 min at room temperature to quench residual aldehyde groups. After washing with PBS, the cells were incubated in 70% deionized formamide at 70°C for 5 min, followed by permeabilization and blocking for 30 min at room temperature in PBS containing 0.5% Triton X-100 and 2% bovine serum albumin (BSA). Cells were prehybridized for 2 h at room temperature in hybridization buffer containing 2% BSA, 4× saline–sodium citrate (SSC), 5% dextran sulfate, 35% deionized formamide, and 10 U RNase inhibitor per 100 µL of buffer. Hybridization was performed overnight at room temperature in a humidified chamber using FITC-conjugated oligo(dT)_30_ at a final concentration of 300 ng/mL in the same hybridization buffer. Slides were subsequently washed once with 4× SSC containing 35% deionized formamide and once with 4× SSC. Nuclear DNA was counterstained with Hoechst 33342 diluted 1:5,000, and the cells were mounted using Fluoro-Long antifade mounting medium (Tokyo Chemical Industry, Japan).

For simultaneous detection of poly(A)^+^ RNA and HA-tagged proteins in cells expressing wild-type or mutant constructs, immunofluorescence staining was performed before RNA FISH. Cells were fixed with 4% PFA, washed three times with PBS, permeabilized with 0.2% Triton X-100 for 10 min, and blocked with 1% BSA in PBS for 10 min. The cells were incubated with an anti-HA primary antibody diluted 1:500 for 1 h at room temperature and then washed three times with PBS. The primary antibody was stabilized by post-fixation with 4% PFA for 10 min, followed by three washes with PBS and incubation with 25 mM NH_4_Cl for 10 min at room temperature. The cells were then subjected to formamide treatment, permeabilization, prehybridization, and overnight oligo(dT)_30_ hybridization as described above. After hybridization, the slides were washed once with 4× SSC containing 35% deionized formamide, once with 4× SSC, once with 2× SSC, and once with PBS. Cells were incubated for 1 h at room temperature with an Alexa Fluor 568-conjugated secondary antibody diluted 1:1,000 and Hoechst 33342 diluted 1:5,000. After three washes with PBS, the cells were mounted using Fluoro-Long antifade mounting medium (Tokyo Chemical Industry, Japan). Images were acquired using Nikon Ti2 AX/AX-R confocal microscope and analyzed with NIS-Elements software. Quantification of poly(A)^+^ RNA distribution was performed using the optical section corresponding to the maximal nuclear cross-sectional area, identified from confocal z-stacks acquired under identical imaging conditions. Nuclear-to-cytoplasmic (N/C) fluorescence intensity ratios were calculated for individual cells and used for statistical comparison between experimental groups. A total of 66 cells per experimental group were analyzed for N/C fluorescence intensity measurements. Statistical significance for comparisons between the pSAP2 control and Nup98-like-gs groups was assessed using a two-tailed unpaired *t*-test with Welch’s correction, whereas comparisons among the three EhNup53-like groups were performed using one-way analysis of variance (ANOVA). A *P* value <0.05 was considered statistically significant.

### Immunoblot analysis

Trophozoites of *E. histolytica* were harvested during exponential growth and washed three times with phosphate-buffered saline (PBS). Where indicated, trophozoites expressing HA-tagged overexpression or mutant EhNup98-like and EhNup53-like constructs were used. Cells were lysed in buffer containing 150 mM NaCl, 50 mM Tris-HCl, pH 7.5, 0.1% Triton X-100, 0.5 mg/ml E-64, and 1x cOmplete Mini protease inhibitor cocktail (Roche, Mannheim, Germany) and incubated on ice for 30 min. Lysates were centrifuged at 16,000 × *g* for 20 min at 4°C. Approximately 30 µg of total protein was resolved by SDS–PAGE (5–20% gradient gels) and transferred onto 0.45 µm PVDF membranes (Millipore Immobilon-P, Massachusetts, USA). Membranes were blocked with 5% skim milk in TBST (50 mM Tris-HCl, pH 8.0, 150 mM NaCl, 0.05% Tween-20) for 30 min at room temperature. Blots were incubated overnight at 4°C with primary antibodies diluted in TBST, including mouse anti-HA (clone 16B12, 1:1,000), rabbit anti-CS1 (1:1,000), or rabbit anti-Histone H3 (1:1,000)(20,64). After washing with TBST, membranes were incubated with HRP-conjugated secondary antibodies (anti-mouse or anti-rabbit IgG, 1:5,000) for 1 h at room temperature. Signals were detected using a chemiluminescent HRP substrate (Millipore, Massachusetts, USA), and visualized with ChemiDoc (Bio-Rad, USA) imaging systems according to the manufacturer’s protocols.

### Nuclear Fractionation

Nuclear and cytoplasmic fractions were prepared using a Nuclear Extraction Kit (Active Motif, Carlsbad, CA, USA) according to the manufacturer’s instructions. Briefly, trophozoites were washed in 10 mL ice-cold PBS supplemented with phosphatase inhibitors and pelleted by centrifugation at 500 × *g* for 5 min at 4°C. The cell pellet was resuspended in 1× hypotonic buffer and incubated on ice for 30 min. Following addition of detergent, the suspension was centrifuged at 14,000 × *g* for 2 min at 4°C. The supernatant was collected as the cytoplasmic fraction, while the pellet containing the nuclei was retained on ice.

The nuclear pellet was resuspended in complete digestion buffer (Digestion Buffer supplemented with protease inhibitor cocktail and 100 mM PMSF) and incubated with enzymatic shearing cocktail for 90 min at 4°C with gentle mixing. The reaction was terminated by addition of 0.5 M EDTA, followed by incubation on ice for 5 min. The resulting nuclear extract was stored at −80°C until further use.

### Blue native PAGE

Blue native polyacrylamide gel electrophoresis (BN-PAGE) was performed using the NativePAGE Novex Bis-Tris Gel System (Thermo Fisher Scientific) according to the manufacturer’s instructions. Trophozoites were harvested and mechanically homogenized using a Dounce homogenizer until approximately 20% cell disruption was achieved. Digitonin then added to final concentrations of 0.1%, 0.5%, or 1% for membrane solubilization and mixed with NativePAGE sample buffer supplemented with Coomassie Brilliant Blue G-250. Samples were resolved using an XCell SureLock Mini-Cell system (Thermo Fisher Scientific) and subsequently transferred onto PVDF membranes for immunoblot analysis.

### Size Exclusion Chromatography

Approximately 1×10^6^ trophozoites expressing HA-tagged constructs were cultured in 10-cm dishes containing BIS medium under low-oxygen conditions using Anaerocult (Merck, USA) at 35.5°C for 48 h. Trophozoites were harvested and mechanically disrupted using a Dounce homogenizer until approximately 20% cell lysis was achieved. The cell suspension was then incubated in lysis buffer containing 20 mM HEPES, pH 7.0, 150 mM NaCl, 0.1% digitonin, 0.5 mg/mL E-64, and 1× cOmplete Mini protease inhibitor cocktail (Roche, Mannheim, Germany) for 30 min on ice.

Cell lysates were clarified by centrifugation at 16,000 × *g* for 20 min at 4°C. The resulting supernatants were subjected to size-exclusion chromatography using an ÄKTA go system equipped with a Superdex 200 Increase 10/300 GL column (Cytiva). Eluted fractions were collected, pooled as indicated, and subsequently analyzed by immunoblotting.

### Co-immunoprecipitation (Co-IP)

Approximately 1 × 10⁶ trophozoites expressing HA-tagged constructs were cultured in 10-cm dishes in BIS medium under low-oxygen conditions (Anaerocult; Merck, USA) at 35.5 °C for 48 h. Cells were harvested by gentle detachment with ice-cold PBS, incubated on ice for 10 min, and washed three times with PBS. Protein-protein interactions were stabilized by crosslinking with dithiobis(succinimidyl propionate) (DSP; Thermo Fisher Scientific) according to the manufacturer’s instructions. Cells were lysed in 1 mL lysis buffer, and lysates were clarified by centrifugation at 16,000 × *g* for 5 min at 4 °C. The supernatants were pre-cleared by incubation with Protein G Sepharose beads (GE Healthcare) for 1 h at 4 °C with gentle rotation. After removal of the beads by brief centrifugation, the supernatants were incubated with anti-HA agarose-conjugated monoclonal antibody beads (Sigma Aldrich) for 3.5 h at 4 °C with gentle mixing. Beads were collected by centrifugation and washed three times with lysis buffer to remove non-specifically bound proteins. Bound proteins were eluted by incubation with HA peptide (0.2 mg/mL) in lysis buffer overnight at 4 °C.

### Mass spectrometry

Protein identification by mass spectrometry was performed by the Mass Spectrometry and Proteomics Core Facility at Johns Hopkins University. Lyophilized protein samples were resuspended in 40 μL of 20 mM triethylammonium bicarbonate (TEAB; pH 8.0) and reduced with 50 mM dithiothreitol at 60 °C for 1 h. After cooling to room temperature, samples were alkylated with 100 mM chloroacetamide for 15 min in the dark. Samples were diluted with 9 M urea and processed using 30 kDa molecular weight cutoff (MWCO) spin filters pre-equilibrated with distilled water. Following buffer exchange with urea and TEAB, proteins were digested with sequencing-grade protease (400 ng per sample) in 300 μL TEAB overnight at 37 °C. Peptides were collected by centrifugation, and the filters were rinsed with TEAB to recover residual peptides. Combined eluates were acidified, desalted using an Oasis HLB microelution plate (Waters), and dried by vacuum centrifugation. Peptides were reconstituted in 2% acetonitrile containing 0.1% formic acid prior to analysis. Liquid chromatography-tandem mass spectrometry (LC-MS/MS) was performed using a Q Exactive Plus Hybrid Quadrupole-Orbitrap mass spectrometer (Thermo Fisher Scientific).

### Immuno-electron Microscopy

Trophozoites expressing HA-Nup98 and HA-Nup53 were cultured in Diamond’s BI-S-33 (BIS) medium and allowed to adhere to gold disks for 1 h at 35.5°C. Attached cells were rapidly vitrified by plunge-freezing in liquid propane at −175°C. Freeze substitution was performed at −80°C for 48 h in ethanol containing 2% (w/v) tannic acid and 2% (v/v) water. Samples were gradually warmed to −20°C over 4 h and subsequently to 4°C over 1 h. Dehydration was completed by three 30-min washes in absolute ethanol at 4°C. Samples were infiltrated with a 50:50 mixture of ethanol and LR White resin (London Resin Co. Ltd., UK) at 4°C for 30 minutes, followed by three exchanges of 100% LR White resin for 30 minutes each. Polymerization was carried out at 50°C overnight. Ultrathin sections (∼70 nm) were prepared using a diamond knife on an ultramicrotome (Ultracut UCT; Leica Microsystems, Austria) and mounted on nickel grids.

For immunolabeling, sections were incubated overnight at 4°C with mouse anti-HA primary antibody (clone 16B12) and rabbit anti-Lamin B1 antibody (ab16048; Abcam) diluted in 1% BSA in PBS. After washing three times in 1% BSA/PBS, grids were incubated for 2 h at room temperature with 10-nm gold-conjugated goat anti-mouse IgG to detect HA and 5-nm gold-conjugated goat anti-rabbit IgG to detect Lamin B1. Following extensive washing, sections were fixed in 2% glutaraldehyde in 0.1 M phosphate buffer, stained with 2% uranyl acetate for 10 min and lead stain solution (Sigma-Aldrich, Tokyo, Japan) for 3 minutes at room temperature, and air dried. Visualization was performed using a JEOL JEM-1400Plus transmission electron microscope operating at 100 kV. Images were acquired with a CCD camera (EM-14830RUBY2; JEOL Ltd., Tokyo, Japan) at a resolution of 3296 × 2472 pixels. Sample preparation, data acquisition, and technical optimization were performed with the assistance of Tokai Electron Microscopy, Nagoya, Japan.

### Homology search for Nup53/35 RRM domains

Homology searches for Nup53/35 RRM domains across representative eukaryotes were performed using jackhmmer, an iterative profile HMM–based sequence search method(65). The RRM domain sequences of *E. histolytica* Nup53-like and Nup53/35 from *D. discoideum, S. cerevisiae*, and *H. sapiens* were used as queries. Searches were conducted against the UniProt database (release UniProt2023_03) using the default parameters for three iterations. We then examined whether annotated Nup53/35 were recovered among the search hits. An E-value cutoff of < 1 × 10⁻⁵ was used to define significant hits.

### Homology modelling

Homology models of the autoproteolytic domain of EhNup98-like and the RRM domain of *E. histolytica* Nup53-like were generated using the Homology Model application in the Molecular Operating Environment (MOE)(66). The autoproteolytic domain of human Nup98 (PDB ID: 2Q5X) and the RRM domain of human Nup35 (PDB ID: 4LIR) were used as template structures, respectively(67).

## Supporting information

S1 Raw Images

## Acknowledgments

This research is funded by Grants-in-Aid for Scientific Research (C) (JP26K10004 to H.J.S.) from the Japan Society for the Promotion of Science and by the Japanese Government (MEXT) Scholarship from the Ministry of Education, Culture, Sports, Science and Technology (MEXT), Japan (to H.A.). The authors want to thank all members of the Nozaki Lab at the University of Tokyo for their valuable discussion.

## Supporting Information

**S1 Fig. Sequence conservation of EhNup98-like and Nup98-like proteins across *Entamoeba* species. (A)** Full-length amino acid sequence alignment of human Nup98 (HsNup98) and *E. histolytica* Nup98-like (EhNup98-like). Conserved residues are shown as white letters on a red background, while residues with similar physicochemical properties are highlighted in yellow. **(B)** Multiple sequence alignment of Nup98-like proteins from representative *Entamoeba* species, including *E. invadens*, *E. moshkovskii*, *E. dispar*, *E. histolytica*, and *E. nuttalli*. Conserved residues are shown as white letters on a red background, while residues with similar physicochemical properties are highlighted in yellow. The C-terminal peptidase S59 domain is indicated, and the conserved HFT autoproteolytic motif is highlighted in black box. K240 and N248 as catalytic residues and FG residues in N-terminal region are indicated in blue and green lines, respectively. Sequence alignment and visualization are performed using Clustal Omega and ESPript 3.2, respectively(19).

**S2 Fig. The T296A substitution alters the processing and subcellular localization of EhNup98-like. (A)** Immunoblot analysis of trophozoites expressing wild-type HA-Nup98-like or the HA-Nup98-like^T296A^ mutant. Cysteine synthase 1 (CS1) was used as a loading control. **(B)** Representative immunofluorescence images of trophozoites expressing wild-type HA-Nup98-like or HA-Nup98-like^T296A^. HA-tagged proteins were detected with anti-HA antibody (green), and nuclei were stained with Hoechst 33342 (blue). Scale bars, 10 µm

**S3 Fig. Sequence comparison of Nup53-like proteins across *Entamoeba* species.** Multiple sequence alignment of Nup53-like proteins from *E. invadens*, *E. moshkovskii*, *E. dispar*, *E. histolytica*, and *E. nuttalli*. Deep blue and light blue boxes indicate low-complexity repeat regions containing TN[x]H_y_GGN[G/T]H_y_G (H_y_: hydrophobic residue, x: any residue), which show substantial variation in length and organization among *Entamoeba* species. Black boxes indicate the more conserved region corresponding to the predicted RRM-like domain. Conserved residues are shown as white letters on a red background, while residues with similar physicochemical properties are highlighted in yellow. Sequence alignment and visualization are performed using Clustal Omega and ESPript 3.2, respectively(19,67).

**S4 Fig. Comparison of the domain organization of human Nup155 and putative *E. histolytica* Nup155.** Schematic representation of human Nup155 (HsNup155), comprising an N-terminal domain (blue box; residues 1–886) and C-terminal domain (orange box; residues 887–1391), and the putative *E. histolytica* Nup155 (EhNup155-like). Grey box indicates no conserved domains were detected by NCBI Conserved Domain Search.

**S5 Fig. Deletion of the EhNup53-like repeat-rich or RRM-like region does not alter bulk poly(A)^+^ RNA distribution. (A)** Representative RNA fluorescence in situ hybridization (RNA-FISH) and immunofluorescence images of trophozoites expressing wild-type HA-Nup53-like, HA-Nup53-like^Δ37–198^, or HA-Nup53-like^Δ382–448^. Poly(A)^+^ RNA was detected using FITC-conjugated oligo(dT)_30_ probes (green), HA-tagged proteins were detected using anti-HA antibody (red), and nuclei were stained with Hoechst 33342 (blue). Merged fluorescence and differential interference contrast (DIC) images are shown. Scale bars, 20 µm. **(B)** Quantification of the nuclear-to-cytoplasmic (N/C) fluorescence intensity ratio of poly(A)^+^ RNA in the indicated strains. No significant differences were detected among the three groups by one-way ANOVA. Error bars indicate SD.

## References

1. Wilson KL, Dawson SC. Functional evolution of nuclear structure. Journal of Cell Biology. 2011 Oct 17;195(2):171–81. doi:10.1083/jcb.201103171

2. Field MC, Sali A, Rout MP. On a bender—BARs, ESCRTs, COPs, and finally getting your coat. Journal of Cell Biology. 2011 Jun 13;193(6):963–72. doi:10.1083/jcb.201102042

3. Raices M, D’Angelo MA. Nuclear pore complex composition: a new regulator of tissue-specific and developmental functions. Nat Rev Mol Cell Biol. 2012 Nov;13(11):687–99. doi:10.1038/nrm3461

4. Lin DH, Hoelz A. The Structure of the Nuclear Pore Complex (An Update). Annu Rev Biochem. 2019 Jun 20;88(1):725–83. doi:10.1146/annurev-biochem-062917-011901

5. Frey S, Görlich D. A Saturated FG-Repeat Hydrogel Can Reproduce the Permeability Properties of Nuclear Pore Complexes. Cell. 2007 Aug;130(3):512–23. doi:10.1016/j.cell.2007.06.024

6. Hülsmann BB, Labokha AA, Görlich D. The Permeability of Reconstituted Nuclear Pores Provides Direct Evidence for the Selective Phase Model. Cell. 2012 Aug;150(4):738–51. doi:10.1016/j.cell.2012.07.019

7. Hampoelz B, Andres-Pons A, Kastritis P, Beck M. Structure and Assembly of the Nuclear Pore Complex. Annu Rev Biophys. 2019 May 6;48(1):515–36. doi:10.1146/annurev-biophys-052118-115308

8. Field MC, Koreny L, Rout MP. Enriching the Pore: Splendid Complexity from Humble Origins. Traffic. 2014 Feb;15(2):141–56. doi:10.1111/tra.12141

9. Padilla-Mejia NE, Field MC. Evolutionary, structural and functional insights in nuclear organisation and nucleocytoplasmic transport in trypanosomes. FEBS Letters. 2023 Oct;597(20):2501–18. doi:10.1002/1873-3468.14747

10. Kehrer J, Kuss C, Andres-Pons A, Reustle A, Dahan N, Devos D, et al. Nuclear Pore Complex Components in the Malaria Parasite Plasmodium berghei. Sci Rep. 2018 Jul 26;8(1):11249. doi:10.1038/s41598-018-29590-5

11. Devos D, Dokudovskaya S, Williams R, Alber F, Eswar N, Chait BT, et al. Simple fold composition and modular architecture of the nuclear pore complex. Proc Natl Acad Sci USA. 2006 Feb 14;103(7):2172–7. doi:10.1073/pnas.0506345103

12. Schmidt HB, Görlich D. Nup98 FG domains from diverse species spontaneously phase-separate into particles with nuclear pore-like permselectivity. eLife. 2015 Jan 6;4:e04251. doi:10.7554/eLife.04251

13. Ren Y, Seo HS, Blobel G, Hoelz A. Structural and functional analysis of the interaction between the nucleoporin Nup98 and the mRNA export factor Rae1. Proc Natl Acad Sci USA. 2010 Jun 8;107(23):10406–11. doi:10.1073/pnas.1005389107

14. Blevins MB, Smith AM, Phillips EM, Powers MA. Complex Formation among the RNA Export Proteins Nup98, Rae1/Gle2, and TAP. Journal of Biological Chemistry. 2003 Jun;278(23):20979–88. doi:10.1074/jbc.M302061200

15. Iwamoto M, Asakawa H, Hiraoka Y, Haraguchi T. Nucleoporin Nup98: a gatekeeper in the eukaryotic kingdoms. Genes to Cells. 2010 Jun;15(7):661–9. doi:10.1111/j.1365-2443.2010.01415.x

16. Gwairgi MA, Ghildyal R. Nuclear transport in *Entamoeba histolytica*: knowledge gap and therapeutic potential. Parasitology. 2018 Sep;145(11):1378–87. doi:10.1017/S0031182018000252

17. Žárský V, Klimeš V, Pačes J, Vlček Č, Hradilová M, Beneš V, et al. The *Mastigamoeba balamuthi* Genome and the Nature of the Free-Living Ancestor of *Entamoeba*. Barlow M, editor. Molecular Biology and Evolution. 2021 May 19;38(6):2240–59. doi:10.1093/molbev/msab020

18. The UniProt Consortium, Bateman A, Martin MJ, Orchard S, Magrane M, Adesina A, et al. UniProt: the Universal Protein Knowledgebase in 2025. Nucleic Acids Research. 2025 Jan 6;53(D1):D609–17. doi:10.1093/nar/gkae1010

19. Sievers F, Wilm A, Dineen D, Gibson TJ, Karplus K, Li W, et al. Fast, scalable generation of high-quality protein multiple sequence alignments using Clustal Omega. Mol Syst Biol. 2011 Oct 11;7(1):MSB201175. doi:10.1038/msb.2011.75

20. Nozaki T, Asai T, Kobayashi S, Ikegami F, Noji M, Saito K, et al. Molecular cloning and characterization of the genes encoding two isoforms of cysteine synthase in the enteric protozoan parasite Entamoeba histolytica. Molecular and Biochemical Parasitology. 1998 Nov;97(1–2):33–44. doi:10.1016/S0166-6851(98)00129-7

21. Kerr SC, Azzouz N, Fuchs SM, Collart MA, Strahl BD, Corbett AH, et al. The Ccr4-Not Complex Interacts with the mRNA Export Machinery. Rossi J, editor. PLoS ONE. 2011 Mar 28;6(3):e18302. doi:10.1371/journal.pone.0018302

22. Gabiatti BP, Krenzer J, Braune S, Krüger T, Zoltner M, Kramer S. Detailed characterisation of the trypanosome nuclear pore architecture reveals conserved asymmetrical functional hubs that drive mRNA export. Schneider A, editor. PLoS Biol. 2025 Feb 3;23(2):e3003024. doi:10.1371/journal.pbio.3003024

23. Ambekar SV, Beck JR, Mair GR. TurboID Identification of Evolutionarily Divergent Components of the Nuclear Pore Complex in the Malaria Model Plasmodium berghei. Miller LH, editor. mBio. 2022 Oct 26;13(5):e01815–22. doi:10.1128/mbio.01815-22

24. Handa N, Kukimoto-Niino M, Akasaka R, Kishishita S, Murayama K, Terada T, et al. The Crystal Structure of Mouse Nup35 Reveals Atypical RNP Motifs and Novel Homodimerization of the RRM Domain. Journal of Molecular Biology. 2006 Oct;363(1):114–24. doi:10.1016/j.jmb.2006.07.089

25. Petrovic S, Samanta D, Perriches T, Bley CJ, Thierbach K, Brown B, et al. Architecture of the linker-scaffold in the nuclear pore. Science. 2022 Jun 10;376(6598):eabm9798. doi:10.1126/science.abm9798

26. DeGrasse JA, DuBois KN, Devos D, Siegel TN, Sali A, Field MC, et al. Evidence for a Shared Nuclear Pore Complex Architecture That Is Conserved from the Last Common Eukaryotic Ancestor*□S.

27. Obado SO, Brillantes M, Uryu K, Zhang W, Ketaren NE, Chait BT, et al. Interactome Mapping Reveals the Evolutionary History of the Nuclear Pore Complex. Schwartz TU, editor. PLoS Biol. 2016 Feb 18;14(2):e1002365. doi:10.1371/journal.pbio.1002365

28. Iwamoto M, Asakawa H, Hiraoka Y, Haraguchi T. Nucleoporin Nup98: a gatekeeper in the eukaryotic kingdoms. Genes to Cells. 2010 Jun;15(7):661–9. doi:10.1111/j.1365-2443.2010.01415.x

29. Ibáñez De Opakua A, Pantoja CF, Cima-Omori MS, Dienemann C, Zweckstetter M. Impact of distinct FG nucleoporin repeats on Nup98 self-association. Nat Commun. 2024 May 7;15(1):3797. doi:10.1038/s41467-024-48194-4

30. Denning DP, Rexach MF. Rapid Evolution Exposes the Boundaries of Domain Structure and Function in Natively Unfolded FG Nucleoporins. Molecular & Cellular Proteomics. 2007 Feb;6(2):272–82. doi:10.1074/mcp.M600309-MCP200

31. Sun Y, Guo H. Structural constraints on autoprocessing of the human nucleoporin Nup98. Protein Science. 2008 Mar;17(3):494–505. doi:10.1110/ps.073311808

32. Rosenblum JS, Blobel G. Autoproteolysis in nucleoporin biogenesis. Proc Natl Acad Sci USA. 1999 Sep 28;96(20):11370–5. doi:10.1073/pnas.96.20.11370

33. Pohl F, Seufert F, Chung YK, Volke D, Hoffmann R, Schöneberg T, et al. Structural basis of GAIN domain autoproteolysis and cleavage-resistance in the adhesion G-protein coupled receptors [Internet]. Biochemistry; 2023 [cited 2026 Apr 24]. Available from: http://biorxiv.org/lookup/doi/10.1101/2023.03.12.532270 doi:10.1101/2023.03.12.532270

34. Griffis ER, Xu S, Powers MA. Nup98 Localizes to Both Nuclear and Cytoplasmic Sides of the Nuclear Pore and Binds to Two Distinct Nucleoporin Subcomplexes. Gall J, editor. MBoC. 2003 Feb;14(2):600–10. doi:10.1091/mbc.e02-09-0582

35. Griffis ER, Altan N, Lippincott-Schwartz J, Powers MA. Nup98 Is a Mobile Nucleoporin with Transcription-dependent Dynamics. Gall J, editor. MBoC. 2002 Apr;13(4):1282–97. doi:10.1091/mbc.01-11-0538

36. Oka M, Asally M, Yasuda Y, Ogawa Y, Tachibana T, Yoneda Y. The Mobile FG Nucleoporin Nup98 Is a Cofactor for Crm1-dependent Protein Export. Weis K, editor. MBoC. 2010 Jun;21(11):1885–96. doi:10.1091/mbc.e09-12-1041

37. Rabut G, Doye V, Ellenberg J. Mapping the dynamic organization of the nuclear pore complex inside single living cells. Nat Cell Biol. 2004 Nov;6(11):1114–21. doi:10.1038/ncb1184

38. Labade AS, Karmodiya K, Sengupta K. HOXA repression is mediated by nucleoporin Nup93 assisted by its interactors Nup188 and Nup205. Epigenetics & Chromatin. 2016 Dec;9(1):54. doi:10.1186/s13072-016-0106-0

39. Makio T, Zhang K, Love N, Mast FD, Liu X, Elaish M, et al. SARS-CoV-2 Orf6 is positioned in the nuclear pore complex by Rae1 to inhibit nucleocytoplasmic transport. Corbett A, editor. MBoC. 2024 May 1;35(5):ar62. doi:10.1091/mbc.E23-10-0386

40. Courjol F, Mouveaux T, Lesage K, Saliou JM, Werkmeister E, Bonabaud M, et al. Characterization of a nuclear pore protein sheds light on the roles and composition of the Toxoplasma gondii nuclear pore complex. Cell Mol Life Sci. 2017 Jun;74(11):2107–25. doi:10.1007/s00018-017-2459-3

41. Kimura M, Imamoto N. Biological Significance of the Importin-β Family-Dependent Nucleocytoplasmic Transport Pathways. Traffic. 2014 Jul;15(7):727–48. doi:10.1111/tra.12174

42. Kimura M, Imai K, Morinaka Y, Hosono-Sakuma Y, Horton P, Imamoto N. Distinct mutations in importin-β family nucleocytoplasmic transport receptors transportin-SR and importin-13 affect specific cargo binding. Sci Rep. 2021 Aug 2;11(1):15649. doi:10.1038/s41598-021-94948-1

43. Ström AC, Weis K. Importin-beta-like nuclear transport receptors. Genome Biol. 2001;2(6):REVIEWS3008. doi: 10.1186/gb-2001-2-6-reviews3008

44. Guo Y, Tao T, Wu T, Hou J, Lin W. Nucleoporin Nup98 is an essential factor for ipo4 dependent protein import. J of Cellular Biochemistry. 2024 Jul;125(7):e30573. doi:10.1002/jcb.30573

45. Sarma NJ, Willis K. The new nucleoporin: Regulator of transcriptional repression and beyond. Nucleus. 2012 Nov;3(6):508–15. doi:10.4161/nucl.22427

46. Sumner MC, Brickner J. The Nuclear Pore Complex as a Transcription Regulator. Cold Spring Harb Perspect Biol. 2022 Jan;14(1):a039438. doi:10.1101/cshperspect.a039438

47. Ikegami K, Lieb JD. Nucleoporins and Transcription: New Connections, New Questions. Bickmore WA, editor. PLoS Genet. 2010 Feb 26;6(2):e1000861. doi:10.1371/journal.pgen.1000861

48. Millán-Zambrano G, Chávez S. Nuclear functions of prefoldin. Open Biol. 2014 Jul;4(7):140085. doi:10.1098/rsob.140085

49. Mallik S, Poch D, Burick S, Schlieker C. Protein folding and quality control during nuclear transport. Current Opinion in Cell Biology. 2024 Oct;90:102407. doi:10.1016/j.ceb.2024.102407

50. Collart MA. The Ccr4-Not complex is a key regulator of eukaryotic gene expression. WIREs RNA. 2016 Jul;7(4):438–54. doi:10.1002/wrna.1332

51. Khan T, Douglas GM, Patel P, Nguyen Ba AN, Moses AM. Polymorphism Analysis Reveals Reduced Negative Selection and Elevated Rate of Insertions and Deletions in Intrinsically Disordered Protein Regions. Genome Biol Evol. 2015 Jun;7(6):1815–26. doi:10.1093/gbe/evv105

52. Vollmer B, Schooley A, Sachdev R, Eisenhardt N, Schneider AM, Sieverding C, et al. Dimerization and direct membrane interaction of Nup53 contribute to nuclear pore complex assembly: Nup53 membrane binding promotes NPC assembly. The EMBO Journal. 2012 Oct 17;31(20):4072–84. doi:10.1038/emboj.2012.256

53. Hawryluk-Gara LA, Platani M, Santarella R, Wozniak RW, Mattaj IW. Nup53 Is Required for Nuclear Envelope and Nuclear Pore Complex Assembly. Weis K, editor. MBoC. 2008 Apr;19(4):1753–62. doi:10.1091/mbc.e07-08-0820

54. Stuwe T, Bley CJ, Thierbach K, Petrovic S, Schilbach S, Mayo DJ, et al. Architecture of the fungal nuclear pore inner ring complex. Science. 2015 Oct 2;350(6256):56–64. doi:10.1126/science.aac9176

55. Chatel G, Fahrenkrog B. Dynamics and diverse functions of nuclear pore complex proteins. Nucleus. 2012 Mar;3(2):162–71. doi:10.4161/nucl.19674

56. Palancade B, Doye V. Sumoylating and desumoylating enzymes at nuclear pores: underpinning their unexpected duties? Trends in Cell Biology. 2008 Apr;18(4):174–83. doi:10.1016/j.tcb.2008.02.001

57. Gasser SM, Stutz F. SUMO in the regulation of DNA repair and transcription at nuclear pores. FEBS Letters. 2023 Nov;597(22):2833–50. doi:10.1002/1873-3468.14751

58. Albert S, Schaffer M, Beck F, Mosalaganti S, Asano S, Thomas HF, et al. Proteasomes tether to two distinct sites at the nuclear pore complex. Proc Natl Acad Sci USA. 2017 Dec 26;114(52):13726–31. doi:10.1073/pnas.1716305114

59. Neumann N, Lundin D, Poole AM. Comparative Genomic Evidence for a Complete Nuclear Pore Complex in the Last Eukaryotic Common Ancestor. Fairhead C, editor. PLoS ONE. 2010 Oct 8;5(10):e13241. doi:10.1371/journal.pone.0013241

60. Makarov AA, Padilla-Mejia NE, Field MC. Evolution and diversification of the nuclear pore complex. Biochemical Society Transactions. 2021 Aug 27;49(4):1601–19. doi:10.1042/BST20200570

61. Nozaki T, Asai T, Sanchez LB, Kobayashi S, Nakazawa M, Takeuchi T. Characterization of the Gene Encoding Serine Acetyltransferase, a Regulated Enzyme of Cysteine Biosynthesis from the Protist ParasitesEntamoeba histolyticaand Entamoeba dispar. Journal of Biological Chemistry. 1999 Nov;274(45):32445–52. doi:10.1074/jbc.274.45.32445

62. Schindelin J, Arganda-Carreras I, Frise E, Kaynig V, Longair M, Pietzsch T, et al. Fiji: an open-source platform for biological-image analysis. Nat Methods. 2012 Jul;9(7):676–82. doi:10.1038/nmeth.2019

63. López-Rosas I, Orozco E, Marchat LA, García-Rivera G, Guillen N, Weber C, et al. mRNA Decay Proteins Are Targeted to poly(A)+ RNA and dsRNA-Containing Cytoplasmic Foci That Resemble P-Bodies in Entamoeba histolytica. Stoecklin G, editor. PLoS ONE. 2012 Sep 24;7(9):e45966. doi:10.1371/journal.pone.0045966

64. Lozano-Amado D, Herrera-Solorio AM, Valdés J, Alemán-Lazarini L, Almaraz-Barrera MaDJ, Luna-Rivera E, et al. Identification of repressive and active epigenetic marks and nuclear bodies in Entamoeba histolytica. Parasites Vectors. 2016 Dec;9(1):19. doi:10.1186/s13071-016-1298-7

65. Eddy SR. Accelerated Profile HMM Searches. Pearson WR, editor. PLoS Comput Biol. 2011 Oct 20;7(10):e1002195. doi:10.1371/journal.pcbi.1002195

66. Chemical Computing Group ULC. Molecular Operating Environment (MOE). Montreal, QC, Canada: Chemical Computing Group ULC; 2026.

67. Robert X, Gouet P. Deciphering key features in protein structures with the new ENDscript server. Nucleic Acids Research. 2014 Jul 1;42(W1):W320–4. doi:10.1093/nar/gku316

