## Supplementary figures and images for "Evolutionary diversification of the nuclear pore complex in *Entamoeba histolytica* reveals conserved and lineage-specific nucleoporins"

### S1 Fig

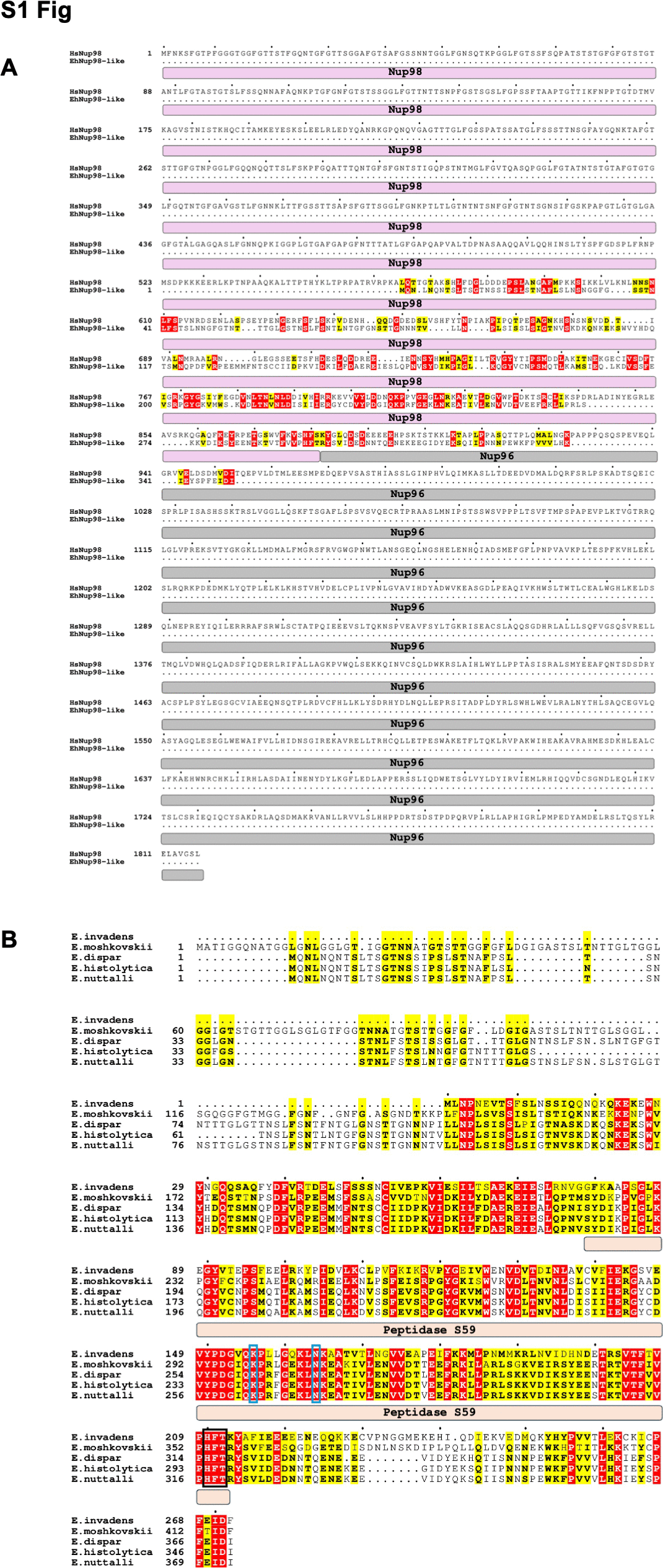

### S2 Fig

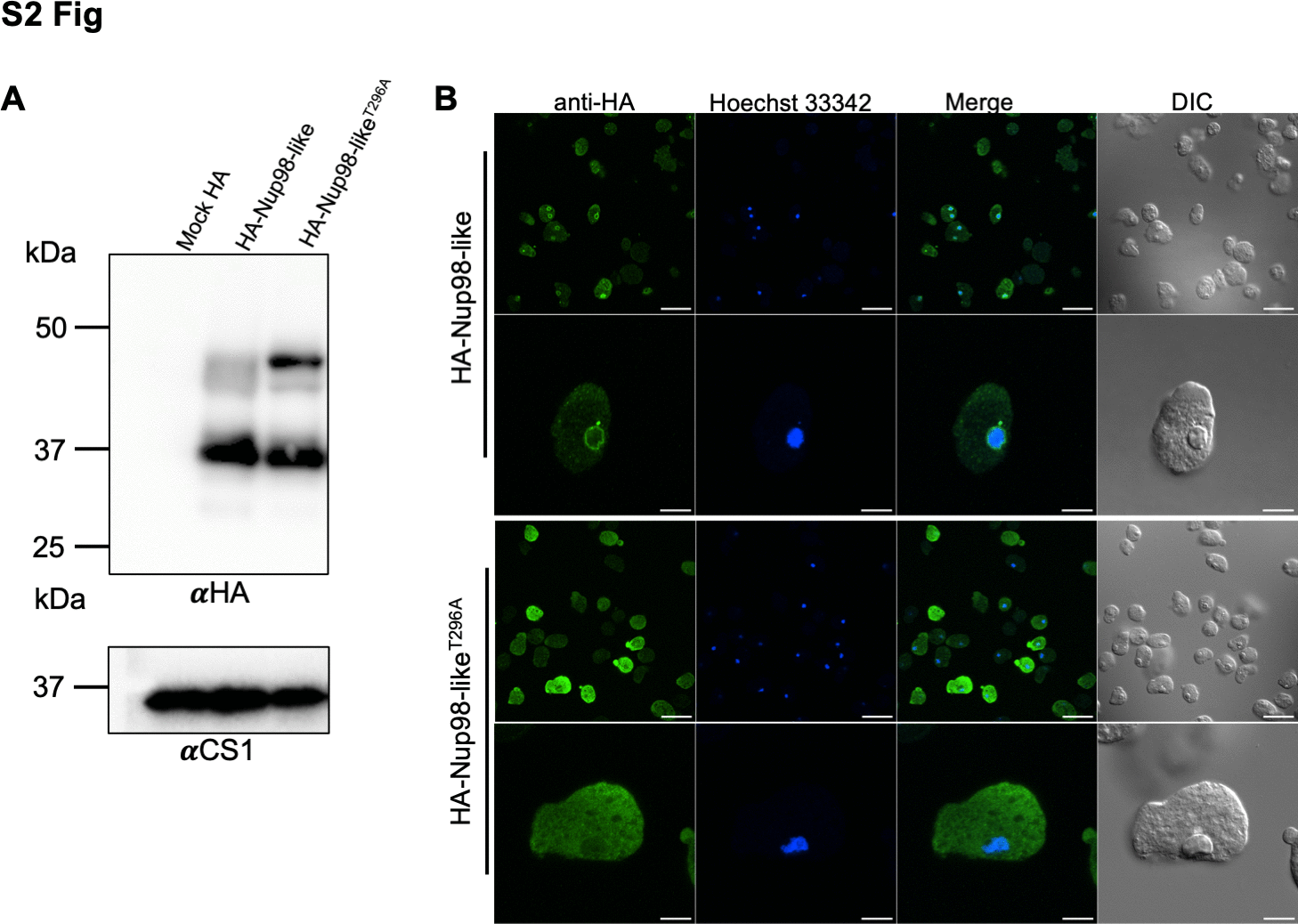

### S3 Fig

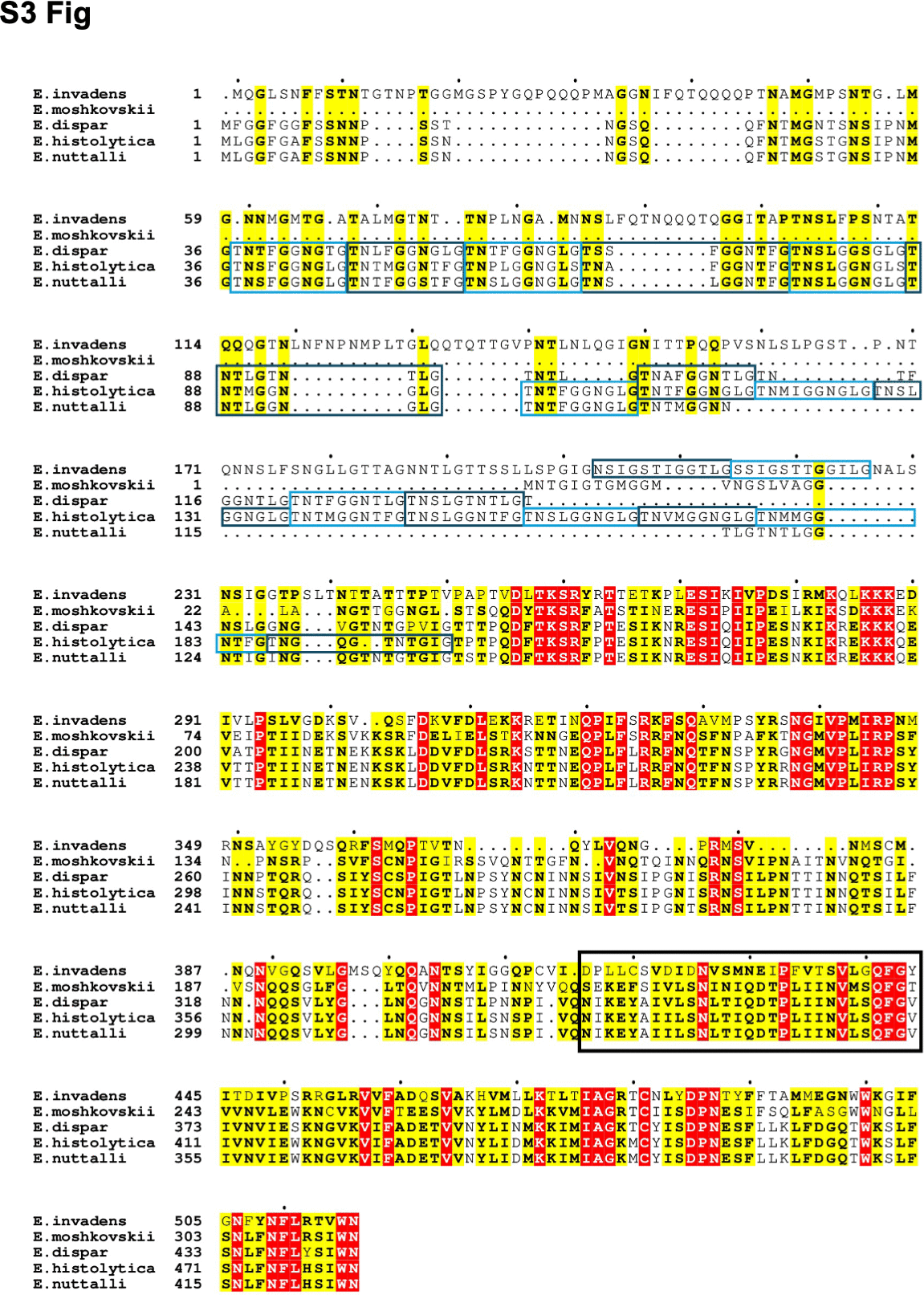

### S4 Fig

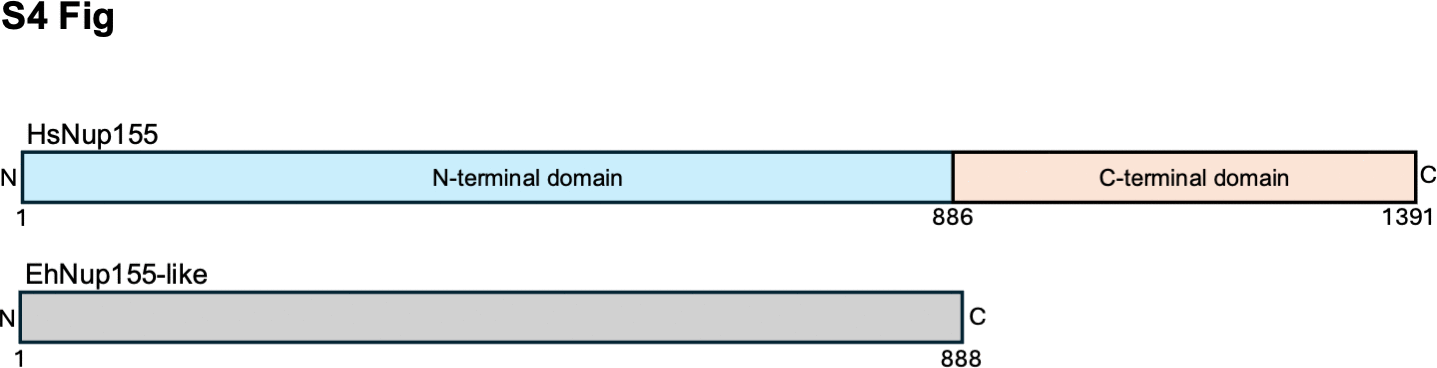

### S5 Fig

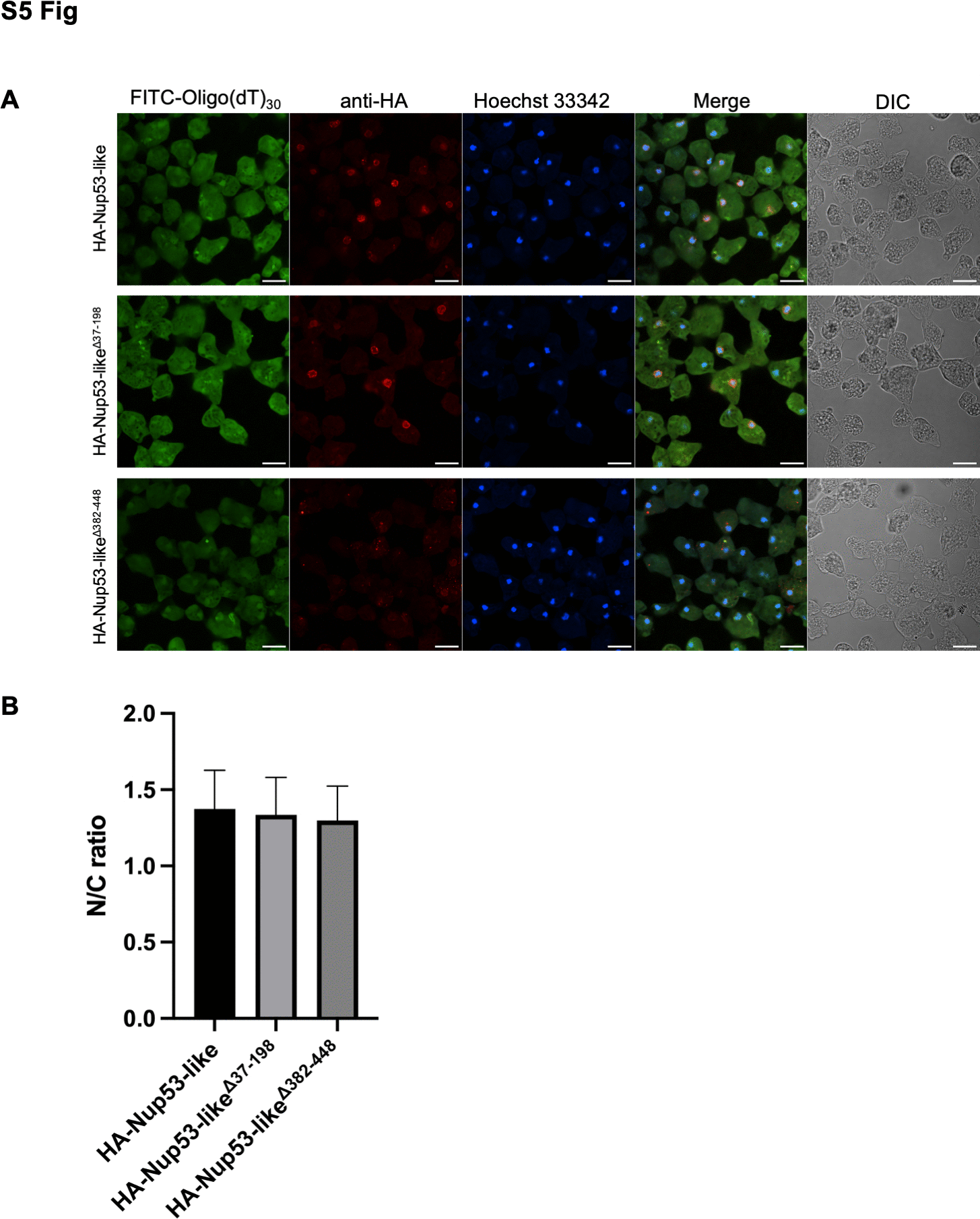
