## Supplementary Tables for "Evolutionary diversification of the nuclear pore complex in *Entamoeba histolytica* reveals conserved and lineage-specific nucleoporins"

**S1 Table. Proteins identified as exclusive interactors of HA–EhNup53-like by immunoprecipitation–mass spectrometry.**

| Accession Number | Annotation | Average Abundances HA-Nup53-like | p-value |
|---|---|---|---|
| EHI_067920 | CDP-alcohol phosphatidyltransferase family protein, putative | 5.E+06 | 0.189 |
| EHI_035190 | Uncharacterized protein | 5.E+06 | 0.135 |
| EHI_075170 | DNA double-strand break repair Rad50 ATPase | 5.E+06 | 0.062 |
| EHI_000700 | NFX1-type zinc finger-containing protein 1 | 4.E+06 | 0.055 |
| EHI_067900 | Serine/threonine-protein phosphatase | 4.E+06 | 0.113 |
| EHI_000240 | Guanine nucleotide-binding protein subunit beta-1, putative | 3.E+06 | 0.149 |
| EHI_194350 | WD domain, G-beta repeat protein | 2.E+06 | 0.275 |
| EHI_179880 | SMP-LTD domain-containing protein | 2.E+06 | 0.187 |
| EHI_009670 | Helicase ATP-binding domain-containing protein | 2.E+06 | 0.092 |
| EHI_110790 | Zinc finger domain containing protein | 2.E+06 | 0.114 |
| EHI_092520 | DUF3340 domain-containing protein | 2.E+06 | 0.140 |
| EHI_054750 | Phosphatidate cytidylyltransferase, putative | 1.E+06 | 0.060 |
| EHI_146350 | Serine/threonine-protein phosphatase | 1.E+06 | 0.119 |
| | Mediator of RNA polymerase II transcription subunit 10 | 1.E+06 | 0.287 |
| EHI_119910 | Lipid phosphate phosphatase, putative | 1.E+06 | 0.070 |
| EHI_006920 | Papain family cysteine protease domain containing protein | 1.E+06 | 0.123 |
| EHI_078200 | Protein kinase | 1.E+06 | 0.201 |
| EHI_110320 | Protein phosphatase, putative | 1.E+06 | 0.248 |
| EHI_107230 | Leucine rich repeat containing protein | 9.E+05 | 0.028 |
| EHI_199640 | Serine/threonine-protein kinase | 9.E+05 | 0.060 |
| EHI_005070 | Acetyltransferase, GNAT family | 9.E+05 | 0.012 |
| EHI_105090 | Importin alpha, putative | 7.E+05 | 0.000 |
| EHI_040280 | Eukaryotic translation initiation factor 4E | 7.E+05 | 0.042 |
| EHI_008370 | Uncharacterized protein | 7.E+05 | 0.040 |
| EHI_053610 | Apyrase, putative | 6.E+05 | 0.000 |
| EHI_007640 | Calponin-homology (CH) domain-containing protein | 6.E+05 | 0.046 |

| | | | |
|---|---|---|---|
| EHI_073670 | WD domain, G-beta repeat-containing protein | 6.E+05 | 0.001 |
| EHI_016410 | AB hydrolase-1 domain-containing protein | 6.E+05 | 0.037 |
| EHI_124570 | Rap-GAP domain-containing protein | 5.E+05 | 0.007 |
| EHI_073310 | MABP domain-containing protein | 5.E+05 | 0.242 |
| EHI_178100 | Protein kinase domain containing protein | 5.E+05 | 0.011 |
| EHI_078180 | Enkurin domain-containing protein | 5.E+05 | 0.109 |
| EHI_030120 | Tyrosine protein kinase, putative | 5.E+05 | 0.011 |
| EHI_060980 | WD domain containing protein | 4.E+05 | 0.002 |
| EHI_200010 | Glycerophosphodiester phosphodiesterase | 4.E+05 | 0.000 |
| EHI_136480 | UEV domain-containing protein | 4.E+05 | 0.000 |
| EHI_104370 | Legume family lectin family protein, membrane-bound, putative | 4.E+05 | 0.019 |
| EHI_074700 | DH domain-containing protein | 3.E+05 | 0.014 |

**S2 Table. Proteins identified as enriched interactors of HA–EhNup53-like by immunoprecipitation–mass spectrometry.**

| Accession Number | Annotation | Abundance Ratio HA-Nup53-like/Mock | Average Abundances HA-Nup53-like | p-value |
|---|---|---|---|---|
| EHI_199660 | PRA1 family protein | 5.08 | 1.2E+09 | 0.699 |
| EHI_125350 | DNA-directed RNA polymerase subunit | 10.43 | 5.0E+08 | 0.224 |
| EHI_179400 | Nucleoporin autopeptidase, putative (Nup98, putative) | 5.83 | 2.7E+08 | 0.097 |
| EHI_193440 | Nucleoporin autopeptidase | 6.73 | 2.1E+08 | 0.576 |
| EHI_059840 | V-type proton ATPase proteolipid subunit | 6.09 | 1.5E+08 | 0.425 |
| EHI_085060 | RanBD1 domain-containing protein | 5.13 | 1.4E+08 | 0.233 |
| EHI_067540 | Uncharacterized protein | 8.37 | 1.3E+08 | 0.467 |
| EHI_065490 | F-BAR domain-containing protein | 5.19 | 1.3E+08 | 0.375 |
| EHI_140310 | Leucine rich repeats family protein | 7.82 | 1.1E+08 | 0.126 |
| EHI_011830 | Uncharacterized protein | 9.54 | 1.0E+08 | 0.130 |
| EHI_080900 | UBA/TS-N domain containing protein | 8.55 | 1.0E+08 | 0.091 |
| EHI_198560 | Palmitoyltransferase | 6.70 | 9.2E+07 | 0.271 |
| EHI_135200 | RRM domain-containing protein | 9.31 | 8.9E+07 | 0.054 |
| EHI_128430 | Protein tyrosine kinase domain-containing protein | 13.44 | 8.7E+07 | 0.637 |
| EHI_109980 | PHR domain-containing protein | 9.52 | 8.1E+07 | 0.138 |
| EHI_078710 | Proteasome subunit beta | 5.42 | 8.0E+07 | 0.521 |
| EHI_068130 | Uncharacterized protein | 6.79 | 7.8E+07 | 0.247 |
| EHI_103610 | Protein kinase, putative | 5.22 | 7.0E+07 | 0.832 |
| EHI_182600 | Proteasome regulatory subunit, putative | 7.09 | 6.1E+07 | 0.091 |
| EHI_155720 | TLDc domain-containing protein | 5.84 | 5.7E+07 | 0.129 |
| EHI_004760 | Proteasome subunit alpha type | 5.21 | 5.6E+07 | 0.106 |
| EHI_098370 | Cleavage stimulation factor, putative | 5.65 | 4.9E+07 | 0.078 |
| EHI_030810 | Malate dehydrogenase, putative | 7.79 | 4.9E+07 | 0.231 |
| EHI_096550 | RNA recognition motif domain containing protein | 10.52 | 4.2E+07 | 0.182 |

| | | | | |
|---|---|---|---|---|
| EHI_062540 | Permease | 6.40 | 4.2E+07 | 0.517 |
| EHI_141040 | ATPase, AAA family protein, putative | 6.31 | 4.1E+07 | 0.305 |
| EHI_049380 | RNA recognition motif domain containing protein | 9.81 | 4.0E+07 | 0.054 |
| EHI_196530 | Ras family protein | 16.57 | 3.9E+07 | 0.105 |
| EHI_048560 | Transcription factor BTF3, putative | 5.15 | 3.9E+07 | 0.185 |
| EHI_177180 | Uncharacterized protein | 5.35 | 3.6E+07 | 0.062 |
| EHI_126140 | 60S ribosomal protein L9, putative | 6.01 | 3.6E+07 | 0.220 |
| EHI_135070 | RNA recognition motif domain containing protein | 5.56 | 3.5E+07 | 0.297 |
| EHI_155590 | Uncharacterized protein | 5.89 | 3.4E+07 | 0.456 |
| EHI_065710 | Uncharacterized protein | 5.56 | 3.4E+07 | 0.570 |
| EHI_073160 | NTR domain-containing protein | 6.49 | 3.4E+07 | 0.513 |
| EHI_170080 | Polyadenylation factor subunit, putative | 7.41 | 3.4E+07 | 0.128 |
| EHI_026470 | Uncharacterized protein | 6.40 | 3.4E+07 | 0.149 |
| EHI_023270 | Ras guanine nucleotide exchange factor, putative | 5.07 | 3.3E+07 | 0.408 |
| EHI_167720 | Proteasome subunit alpha type | 9.87 | 3.3E+07 | 0.301 |
| EHI_051760 | Calponin-homology (CH) domain-containing protein | 19.52 | 3.1E+07 | 0.131 |
| EHI_144610 | Methionine gamma-lyase, putative | 8.83 | 3.1E+07 | 0.145 |
| EHI_088020 | Alcohol dehydrogenase, putative | 6.22 | 3.0E+07 | 0.117 |
| EHI_007000 | Arp2/3 complex-activating protein rickA, putative | 5.35 | 3.0E+07 | 0.209 |
| EHI_004750 | Signal recognition particle 54 kDa protein | 11.82 | 2.8E+07 | 0.892 |
| EHI_141380 | Leucine-rich repeat containing protein | 29.26 | 2.8E+07 | 0.289 |
| EHI_134810 | Rubrerythrin, putative | 21.14 | 2.8E+07 | 0.087 |
| EHI_197340 | Sulfotransferase, putative | 5.06 | 2.7E+07 | 0.539 |
| EHI_097870 | AKAP7_NLS domain-containing protein | 5.32 | 2.5E+07 | 0.322 |
| EHI_098250 | MutT/nudix family protein, putative | 5.09 | 2.4E+07 | 0.209 |
| EHI_194860 | Damaged DNA binding protein, putative | 27.22 | 2.2E+07 | 0.079 |
| EHI_069420 | Uncharacterized protein | 6.55 | 2.1E+07 | 0.113 |
| EHI_169210 | Coatomer subunit delta | 13.02 | 2.1E+07 | 0.075 |

| | | | | |
|---|---|---|---|---|
| EHI_175060 | Tetratricopeptide repeat protein | 6.88 | 2.0E+07 | 0.759 |
| EHI_087570 | Endoribonuclease L-PSP, putative | 15.62 | 2.0E+07 | 0.317 |
| EHI_039060 | HMG (High mobility group) box domain containing protein | 9.20 | 1.9E+07 | 0.944 |
| EHI_192050 | UDP-glucose 4-epimerase | 5.06 | 1.8E+07 | 0.172 |
| EHI_027340 | RNA polymerase II subunit A C-terminal domain phosphatase | 5.14 | 1.7E+07 | 0.126 |
| EHI_145970 | Uncharacterized protein | 6.25 | 1.7E+07 | 0.425 |
| EHI_183420 | Suf domain-containing protein | 12.44 | 1.6E+07 | 0.056 |
| EHI_098430 | VWFA domain-containing protein | 7.42 | 1.6E+07 | 0.939 |
| EHI_009510 | TBC domain containing protein | 9.19 | 1.5E+07 | 0.120 |
| EHI_052140 | Proteasome subunit alpha type | 7.61 | 1.5E+07 | 0.153 |
| EHI_188950 | PH domain-containing protein | 9.37 | 1.4E+07 | 0.057 |
| EHI_146330 | Calpain large subunit domain III containing protein | 5.43 | 1.3E+07 | 0.089 |
| EHI_091940 | Phosphatidylinositol-4,5-bisphosphate 3-kinase catalytic subunit, putative | 6.84 | 1.3E+07 | 0.139 |
| EHI_158160 | LIM zinc finger domain containing protein | 7.18 | 1.2E+07 | 0.698 |
| EHI_042250 | Rab family GTPase | 5.09 | 1.2E+07 | 0.051 |
| EHI_186950 | Protein phosphatase domain-containing protein | 6.68 | 1.2E+07 | 0.051 |
| EHI_004990 | Ankyrin, putative | 15.84 | 1.2E+07 | 0.587 |
| EHI_180770 | EGF-like domain-containing protein | 8.78 | 1.2E+07 | 0.059 |
| EHI_111750 | Importin alpha re-exporter, putative | 16.57 | 1.1E+07 | 0.620 |
| EHI_114330 | Uncharacterized protein | 11.35 | 9.5E+06 | 0.279 |
| EHI_137760 | Leucine rich repeat domain containing protein | 5.66 | 9.0E+06 | 0.569 |
| EHI_087370 | Protein kinase domain containing protein | 6.64 | 8.9E+06 | 0.243 |
| EHI_140590 | Ethanolamine-phosphate cytidylyltransferase | 5.48 | 8.9E+06 | 0.265 |

| | | | | |
|---|---|---|---|---|
| EHI_156430 | t-SNARE coiled-coil homology domain-containing protein | 5.30 | 8.8E+06 | 0.228 |
| EHI_079720 | DnaJ family protein | 5.62 | 8.8E+06 | 0.118 |
| EHI_158560 | Protein arginine N-methyltransferase | 6.58 | 8.7E+06 | 0.308 |
| EHI_092220 | RING-type domain-containing protein | 6.82 | 8.7E+06 | 0.126 |
| EHI_159790 | Ribosome assembly factor | 6.56 | 8.4E+06 | 0.198 |
| EHI_188130 | Sm protein F | 10.64 | 8.0E+06 | 0.709 |
| EHI_067580 | Zinc finger protein, putative | 7.16 | 7.9E+06 | 0.508 |
| EHI_087540 | Phosphoribulokinase/uridine kinase family protein | 9.36 | 7.9E+06 | 0.452 |
| EHI_178670 | Aldose reductase, putative | 5.92 | 7.0E+06 | 0.431 |
| EHI_103300 | Eukaryotic translation initiation factor 4C | 5.01 | 6.8E+06 | 0.144 |
| EHI_135950 | Fip1 domain-containing protein | 32.30 | 6.6E+06 | 0.428 |
| EHI_008160 | Sec1 family protein | 6.97 | 6.6E+06 | 0.176 |
| EHI_126930 | Serine/threonine protein phosphatase type 5, putative | 5.57 | 6.3E+06 | 0.332 |
| EHI_073300 | Pre-mRNA-splicing factor SYF1 | 10.57 | 6.3E+06 | 0.607 |
| EHI_094010 | PWWP domain-containing protein | 8.78 | 6.2E+06 | 0.249 |
| EHI_126150 | Pre-mRNA-splicing factor cwc2, putative | 8.35 | 6.1E+06 | 0.226 |
| EHI_170370 | DUF2726 domain-containing protein | 6.11 | 6.0E+06 | 0.657 |
| EHI_134680 | Leucine-rich repeat containing protein | 5.27 | 5.4E+06 | 0.079 |
| EHI_112010 | ABC transporter, putative | 8.10 | 5.2E+06 | 0.581 |
| EHI_138190 | U-box domain, putative | 5.52 | 5.0E+06 | 0.128 |
| EHI_100110 | Ubiquitin-conjugating enzyme family protein, putative | 6.37 | 5.0E+06 | 0.112 |
| EHI_053450 | DNA-directed RNA polymerase subunit | 7.97 | 4.8E+06 | 0.068 |
| EHI_118140 | Uncharacterized protein | 5.64 | 4.4E+06 | 0.645 |
| EHI_023220 | 60S ribosomal protein L29 | 5.96 | 4.3E+06 | 0.999 |
| EHI_111040 | DEAD/DEAH box helicase, putative | 5.69 | 3.9E+06 | 0.490 |
| EHI_118090 | Uncharacterized protein | 12.79 | 3.4E+06 | 0.487 |
| EHI_103750 | Nucleosome assembly protein, putative | 6.59 | 3.2E+06 | 0.419 |

| | | | | |
|---|---|---|---|---|
| EHI_130690 | Oligo-1,6-glucosidase, putative | 5.29 | 3.0E+06 | 0.606 |
| EHI_194320 | Plus3 domain-containing protein | 7.03 | 2.6E+06 | 0.418 |
| EHI_074780 | Protein kinase domain containing protein | 5.26 | 2.2E+06 | 0.164 |
| EHI_164360 | Zinc ribbon domain-containing protein | 6.27 | 1.1E+06 | 0.139 |

**S3 Table. Primer sequences used in experiments.**

| | | |
|---|---|---|
| HA-Nup98-like | Forward | 5'- (GAACCCGGGATGCAAAACTTAAATCAAAATAC)-3' |
| | Reverse | 5'- (GAACTCGAG TTATATATCAATCTCAAATGGACT)-3' |
| HA-Nup53-like | Forward | 5'- (GAACCCGGGATGTTAGGTGGCTTTGGT)-3' |
| | Reverse | 5'- (GCCCTCGAGTTAATTCCAAATTGAATGAAGGAAAT)-3' |
| HA-Nup98-like$^{T296A}$ | Forward | 5'- ([PHO]CTCGATATTCTGTTATTGATGAAGATAATAACACACAAG)-3' |
| | Reverse | 5'- ([PHO]CAAAATGAGGAACAACAAAAGTAACAGTTTTAGTATTTTC)-3' |
| HA-Nup53-like$^{\Delta 37-198}$ | Forward | 5'- (CATGGGAACACCCACACCA)-3' |
| | Reverse | 5'- (GTGGGTGTTCCCATGTTAGGTATTGA)-3' |
| HA-Nup53-like$^{\Delta 37-198}$ | Forward | 5'- (TCCAATAGATCCAAATGAATCATTCTTATTAAAACTCTTT)-3' |
| | Reverse | 5'- (TTTGGATCTATTGGACTATTTGATAATATAGAATTATTTCCTTG)-3' |
