## Supplementary material for "Evolutionary diversification of the nuclear pore complex in *Entamoeba histolytica* reveals conserved and lineage-specific nucleoporins": S1 Raw Images

### Uncropped Western blots and gels for the figures in:

Each of the full-size, uncropped blot or gel is presented with markers indicated. The red boxes indicate the cropped area shown in the final figure. Asterisks denote lanes not included in the final figure.

**Figure 2A**

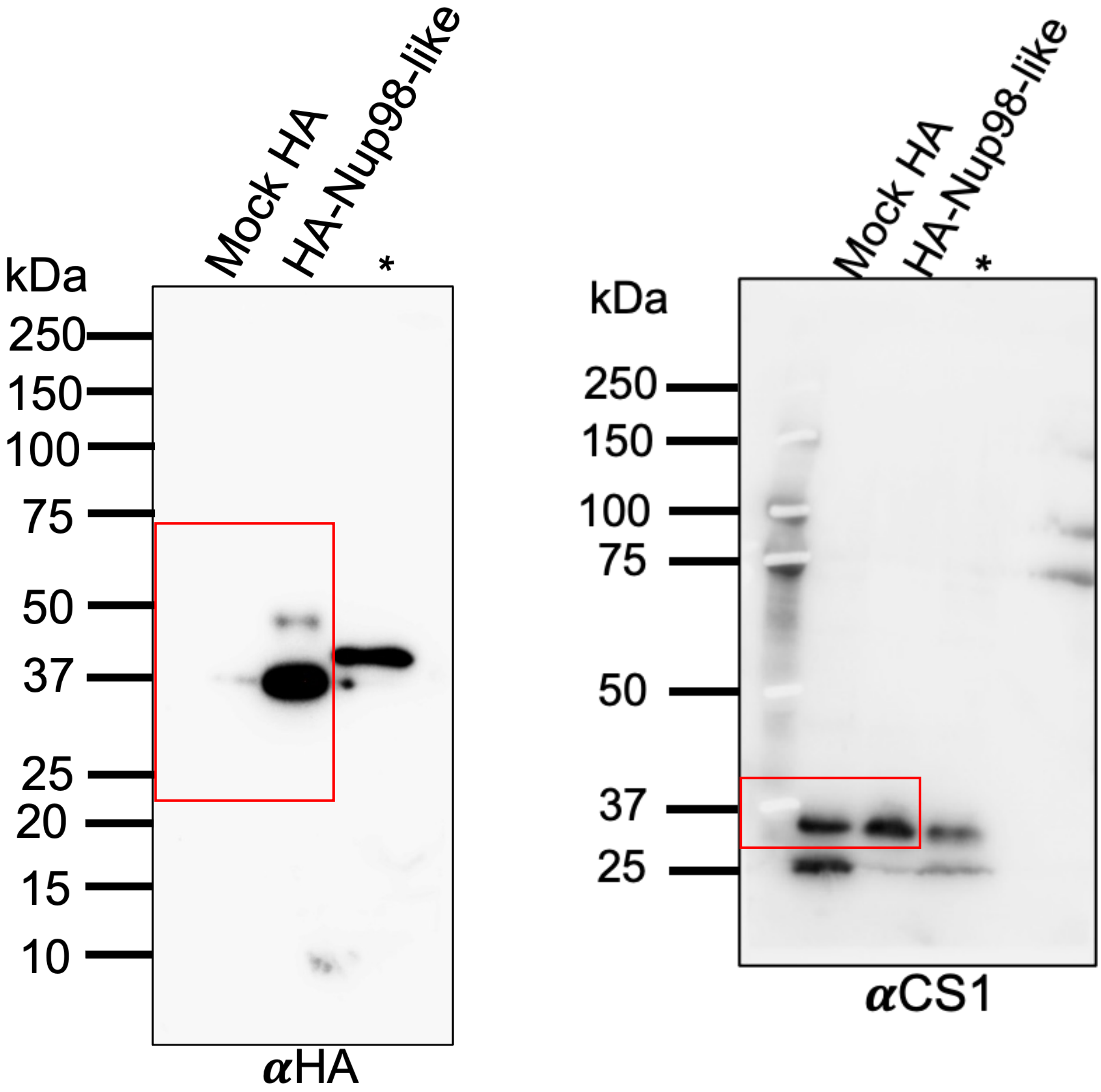
Mock HA
HA-Nup98-like
*
kDa
250
150
100
75
50
37
25
20
15
10
αHA
Mock HA
HA-Nup98-like
*
kDa
250
150
100
75
50
37
25
αCS1

**Figure 2C**

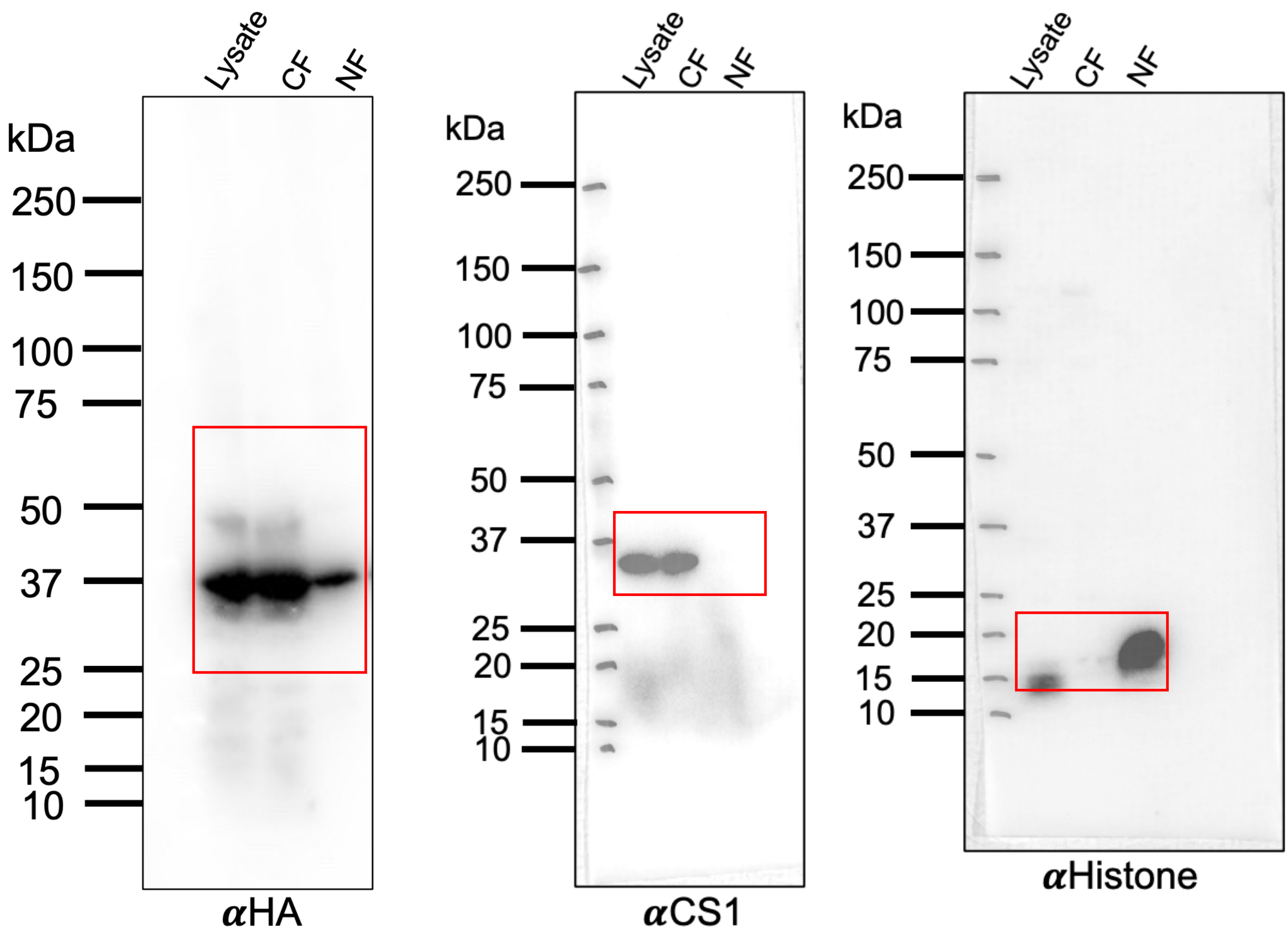
Lysate
CF
NF
kDa
250
150
100
75
50
37
25
20
15
10
αHA
αCS1
αHistone

**Figure 3A**

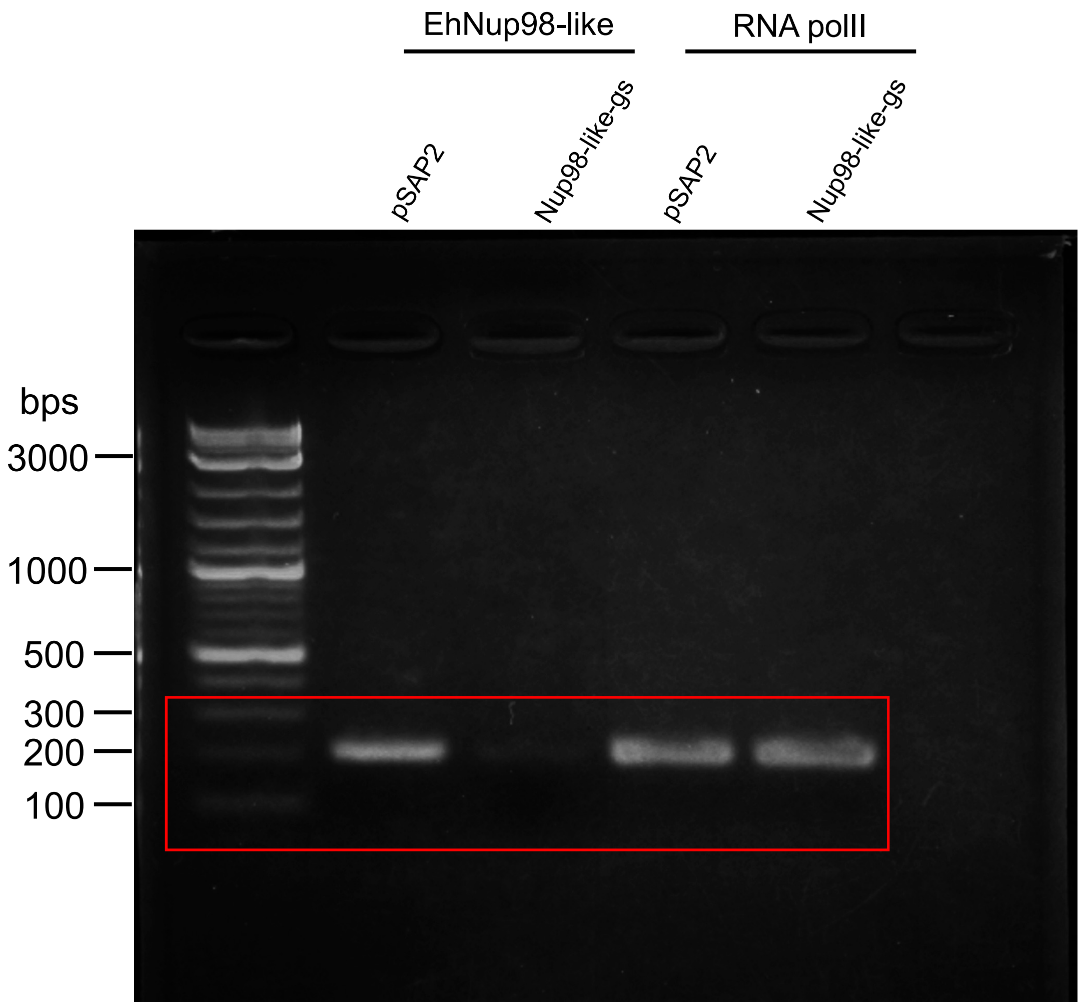
EhNup98-like
RNA polII
pSAP2
Nup98-like-gs
pSAP2
Nup98-like-gs
bps
3000
1000
500
300
200
100

**Figure 4A**

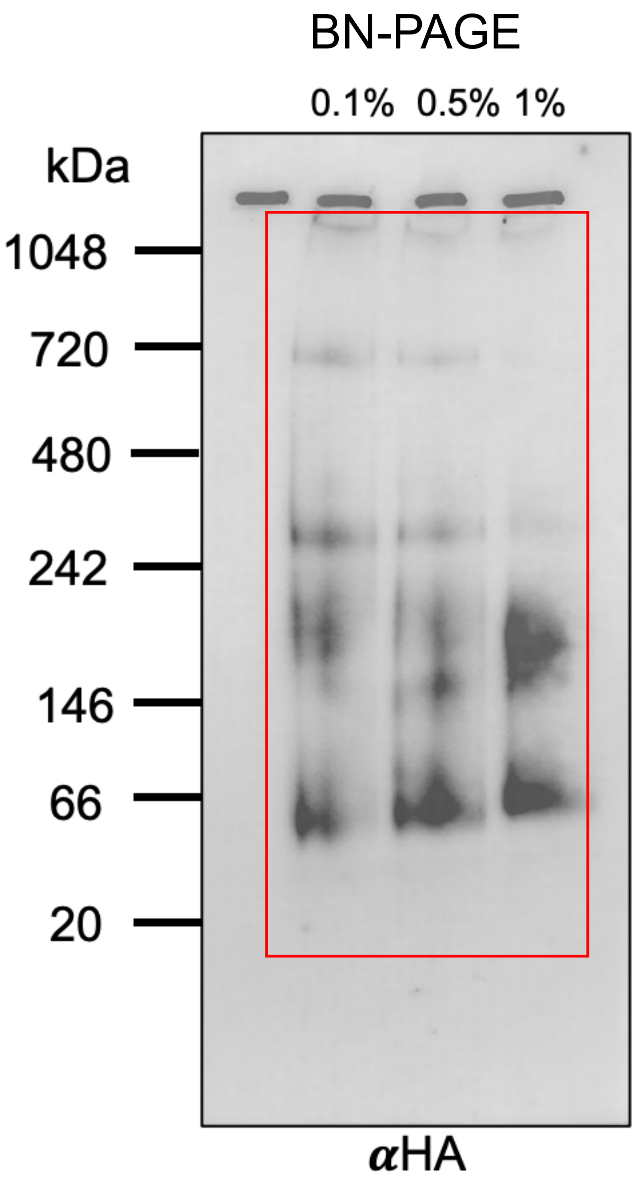
BN-PAGE
0.1% 0.5% 1%
kDa
1048
720
480
242
146
66
20
αHA

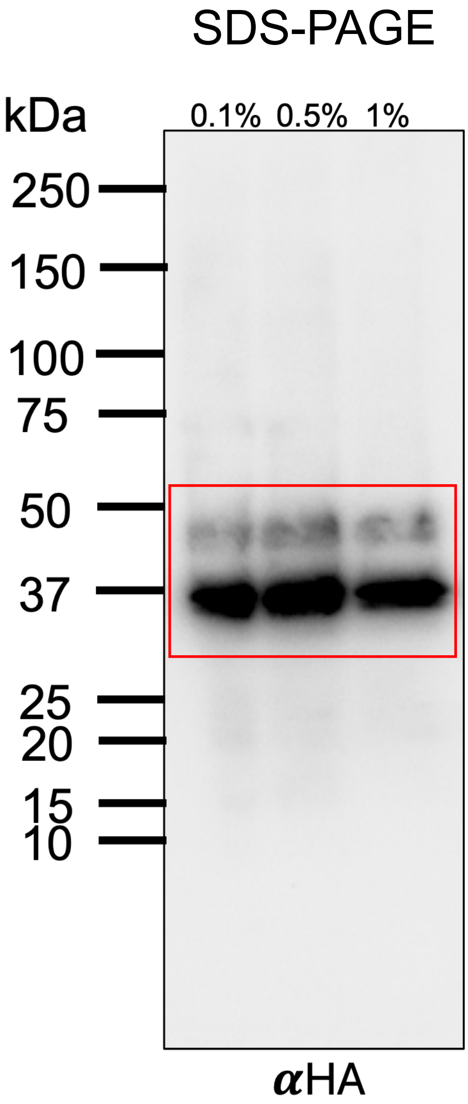
SDS-PAGE
kDa
0.1% 0.5% 1%
250
150
100
75
50
37
25
20
15
10
αHA

**Figure 4B**

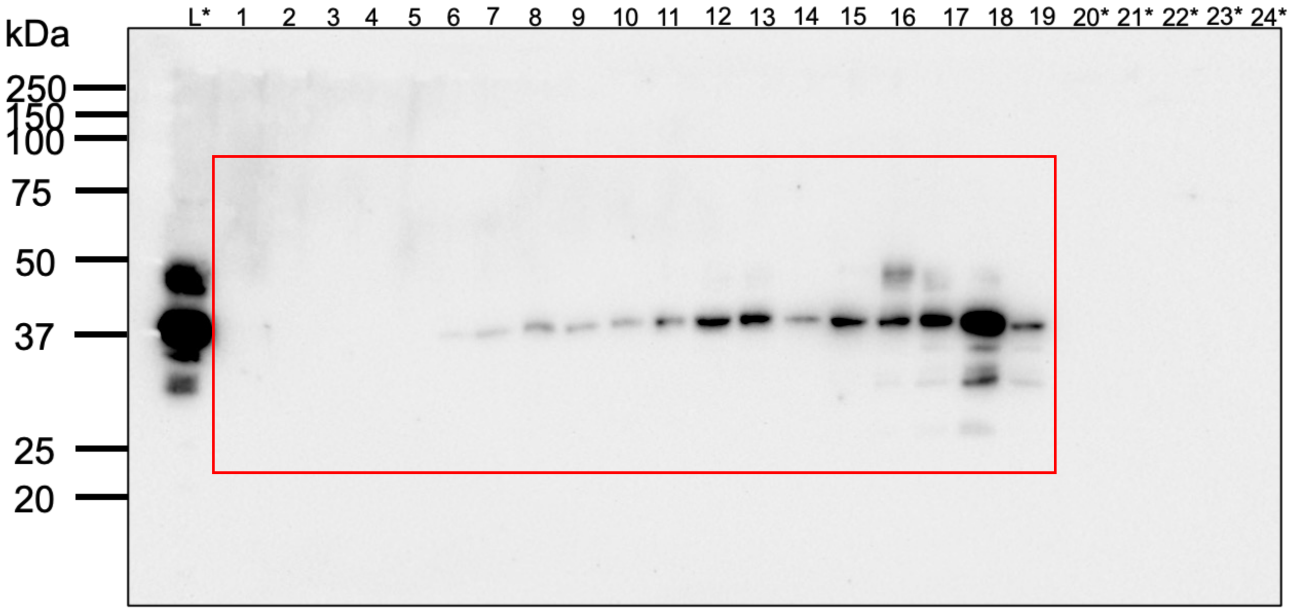
kDa
250
150
100
75
50
37
25
20
L*
20*
21*
22*
23*
24*

Figure 4C

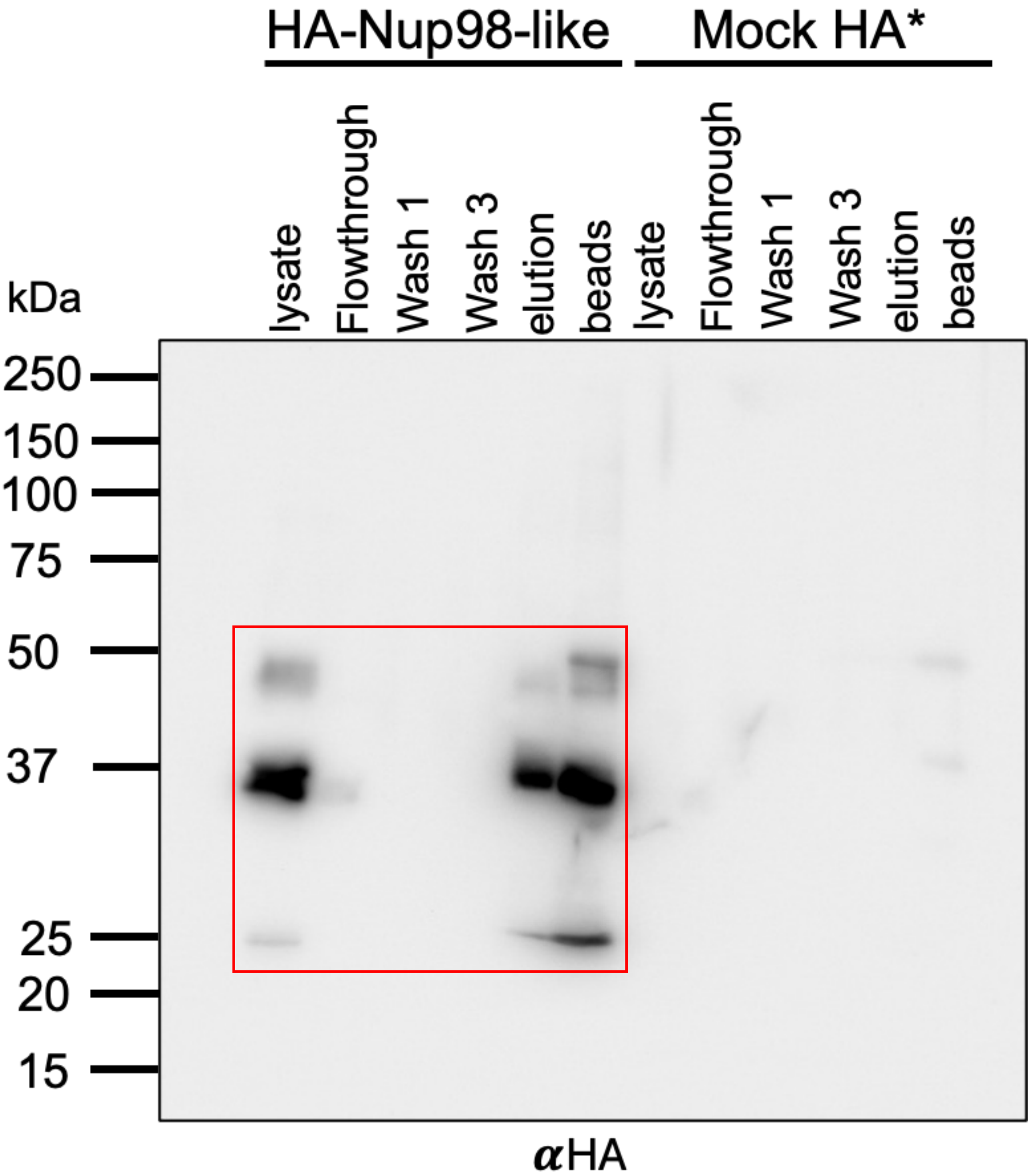
HA-Nup98-like
Mock HA*
lysate
Flowthrough
Wash 1
Wash 3
elution
beads
lysate
Flowthrough
Wash 1
Wash 3
elution
beads
kDa
250
150
100
75
50
37
25
20
15
αHA

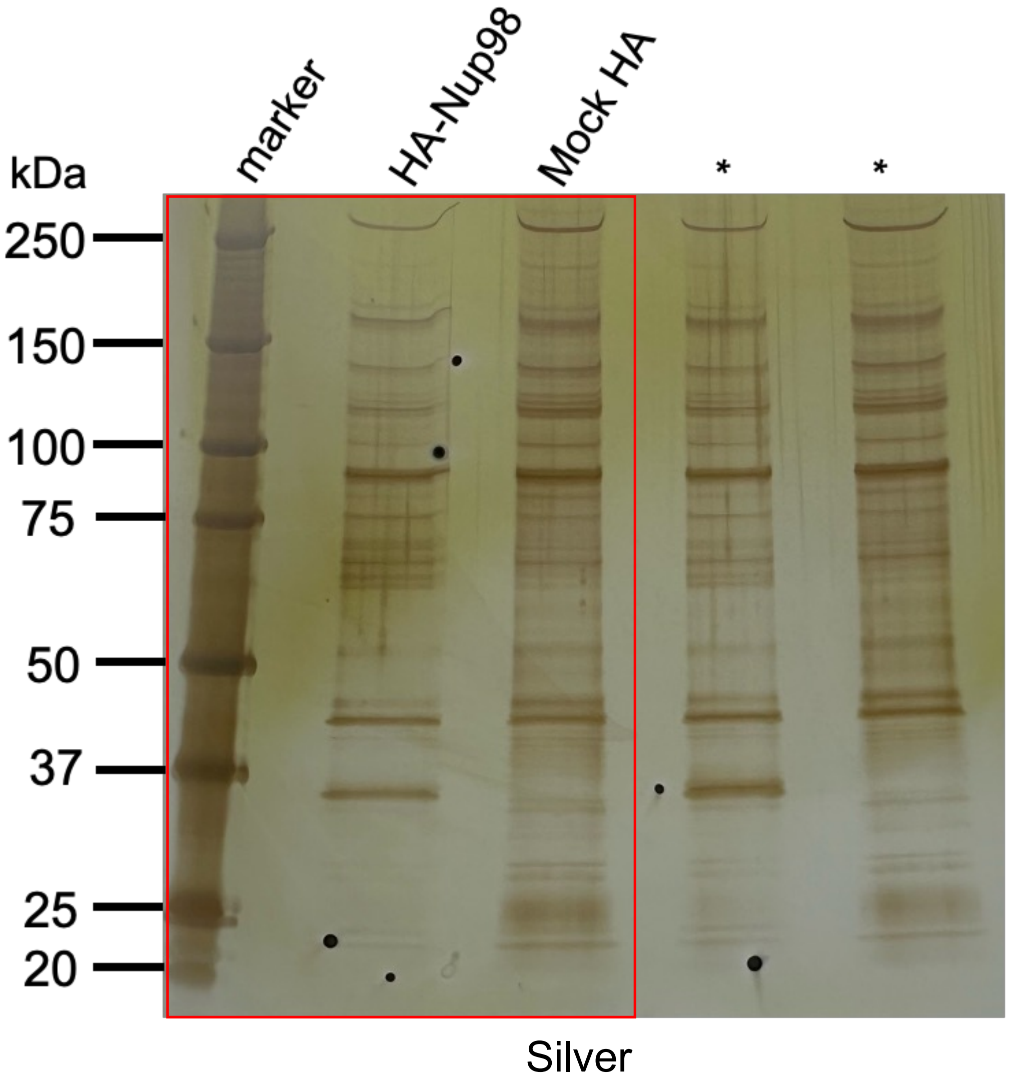
marker
HA-Nup98
Mock HA
*
*
kDa
250
150
100
75
50
37
25
20
Silver

**Figure 6A**

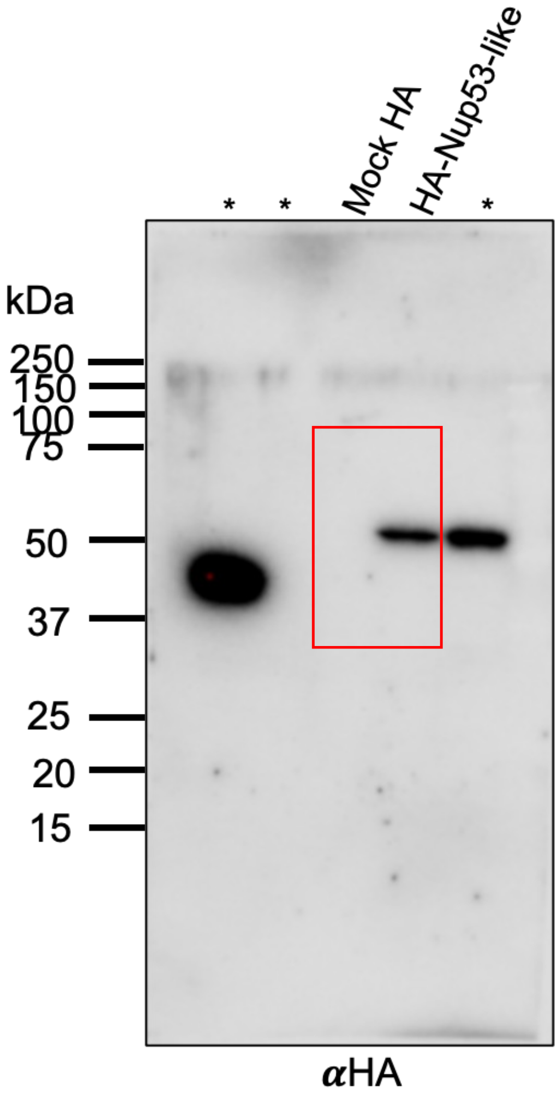

*
*
Mock HA
HA-Nup53-like
*
kDa
250
150
100
75
50
37
25
20
15
αHA

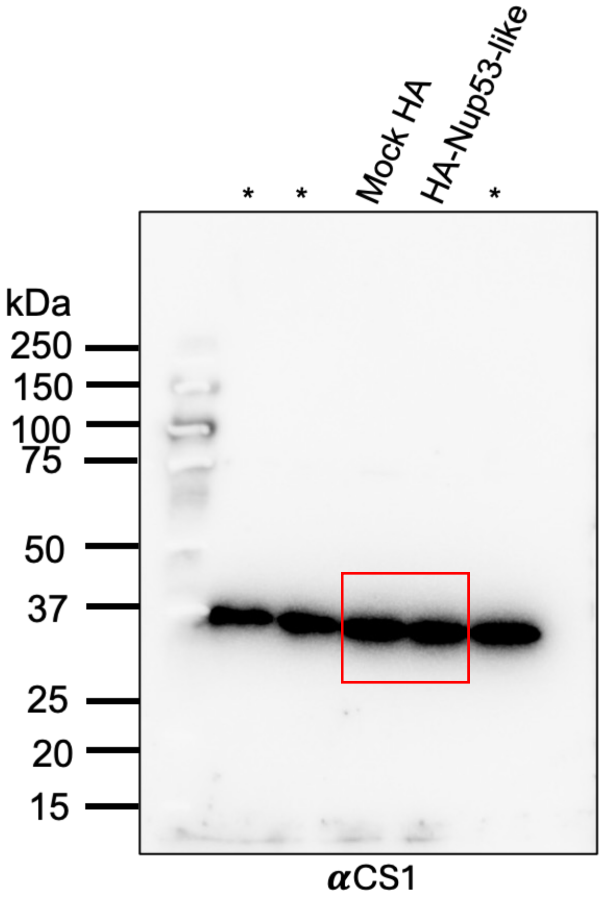

*
*
Mock HA
HA-Nup53-like
*
kDa
250
150
100
75
50
37
25
20
15
αCS1

### Figure 6E

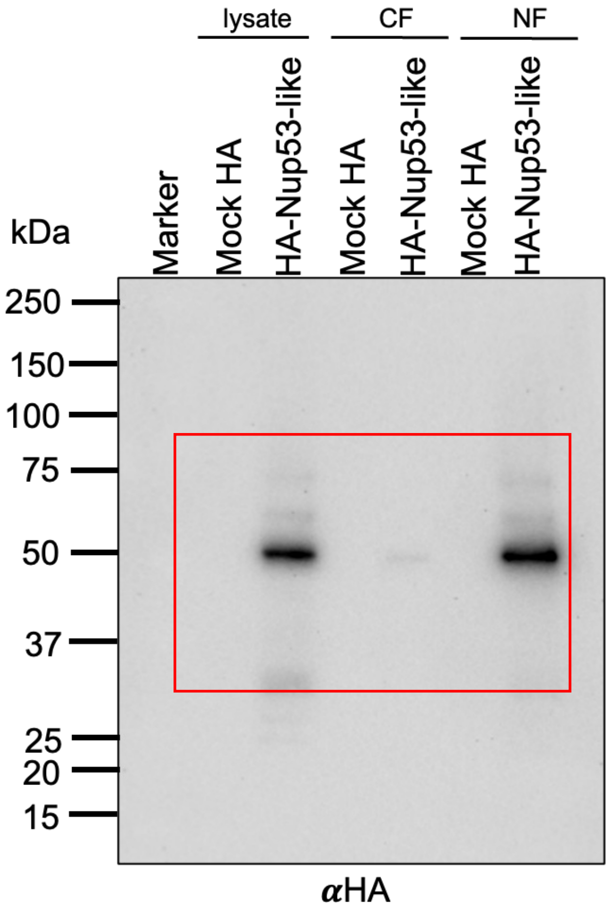

lysate
CF
NF
Marker
Mock HA
HA-Nup53-like
Mock HA
HA-Nup53-like
Mock HA
HA-Nup53-like
kDa
250
150
100
75
50
37
25
20
15
αHA

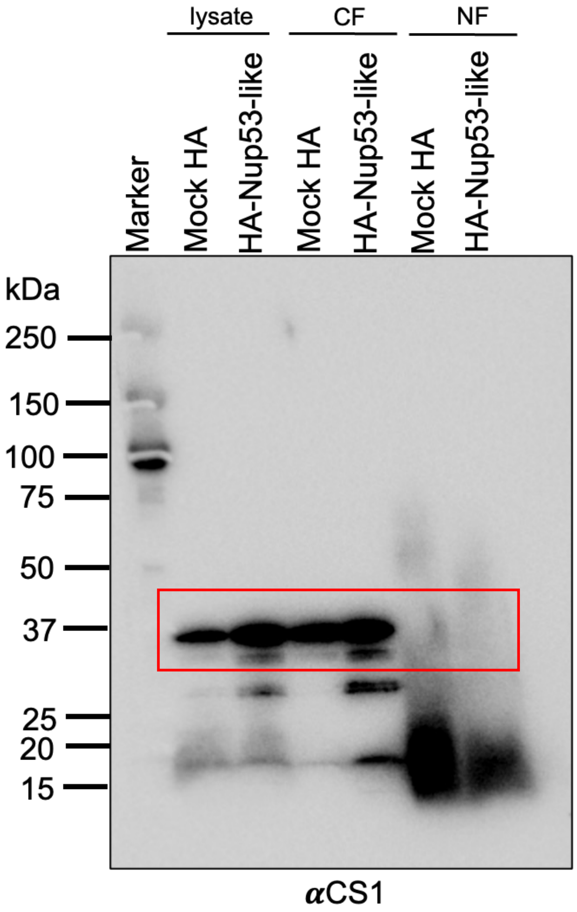

lysate
CF
NF
Marker
Mock HA
HA-Nup53-like
Mock HA
HA-Nup53-like
Mock HA
HA-Nup53-like
kDa
250
150
100
75
50
37
25
20
15
αCS1

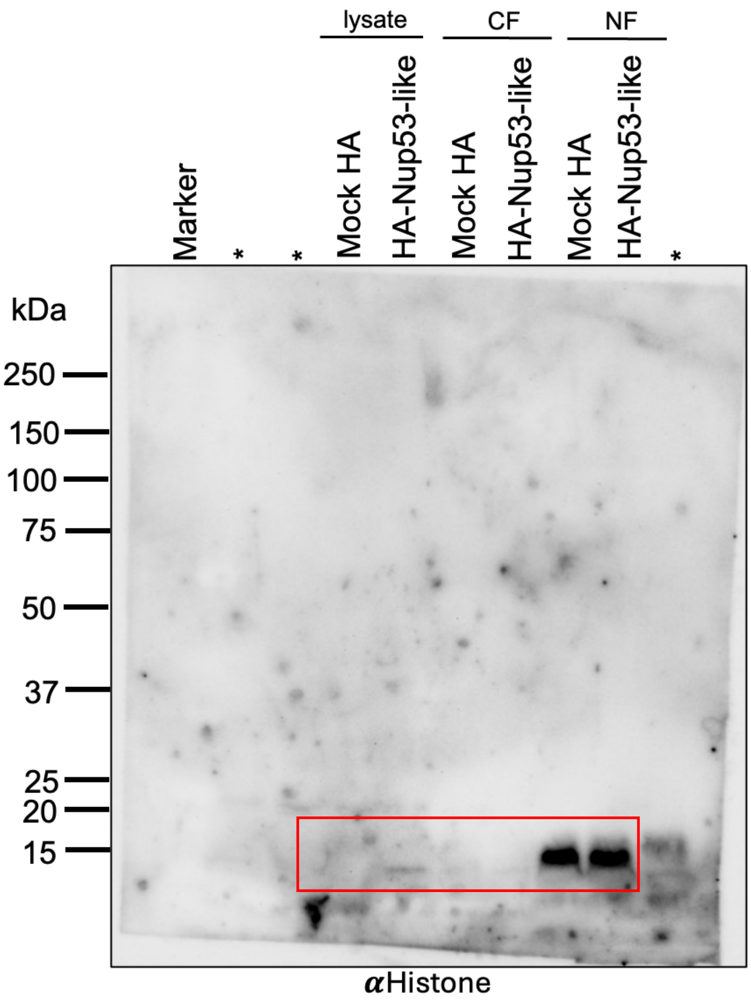

lysate
CF
NF
Marker
*
*
Mock HA
HA-Nup53-like
Mock HA
HA-Nup53-like
Mock HA
HA-Nup53-like
*
kDa
250
150
100
75
50
37
25
20
15
αHistone

Figure 7A

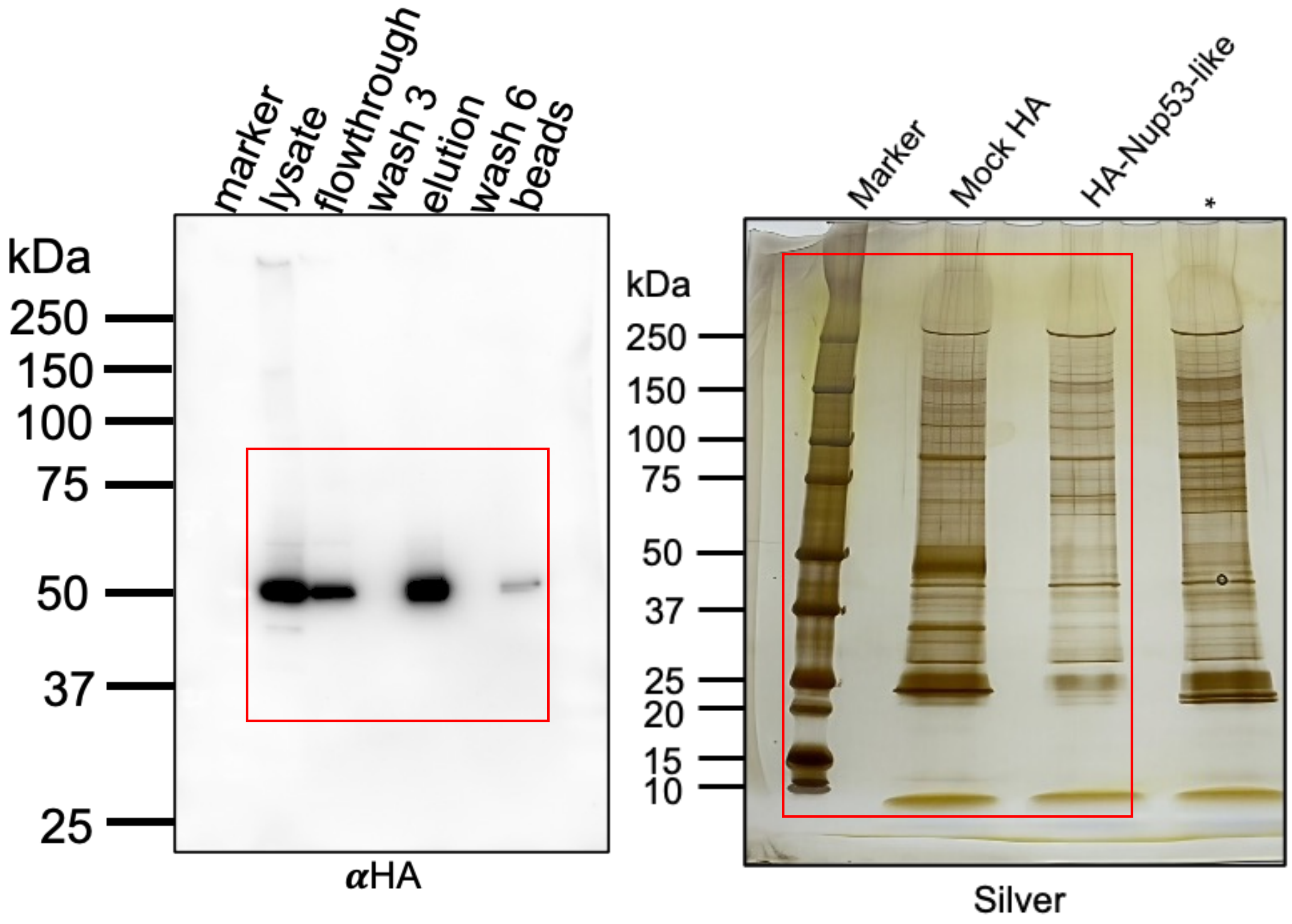
marker
lysate
flowthrough
wash 3
elution
wash 6
beads
kDa
250
150
100
75
50
37
25
αHA
Marker
Mock HA
HA-Nup53-like
*
kDa
250
150
100
75
50
37
25
20
15
10
Silver

**Figure 8A**

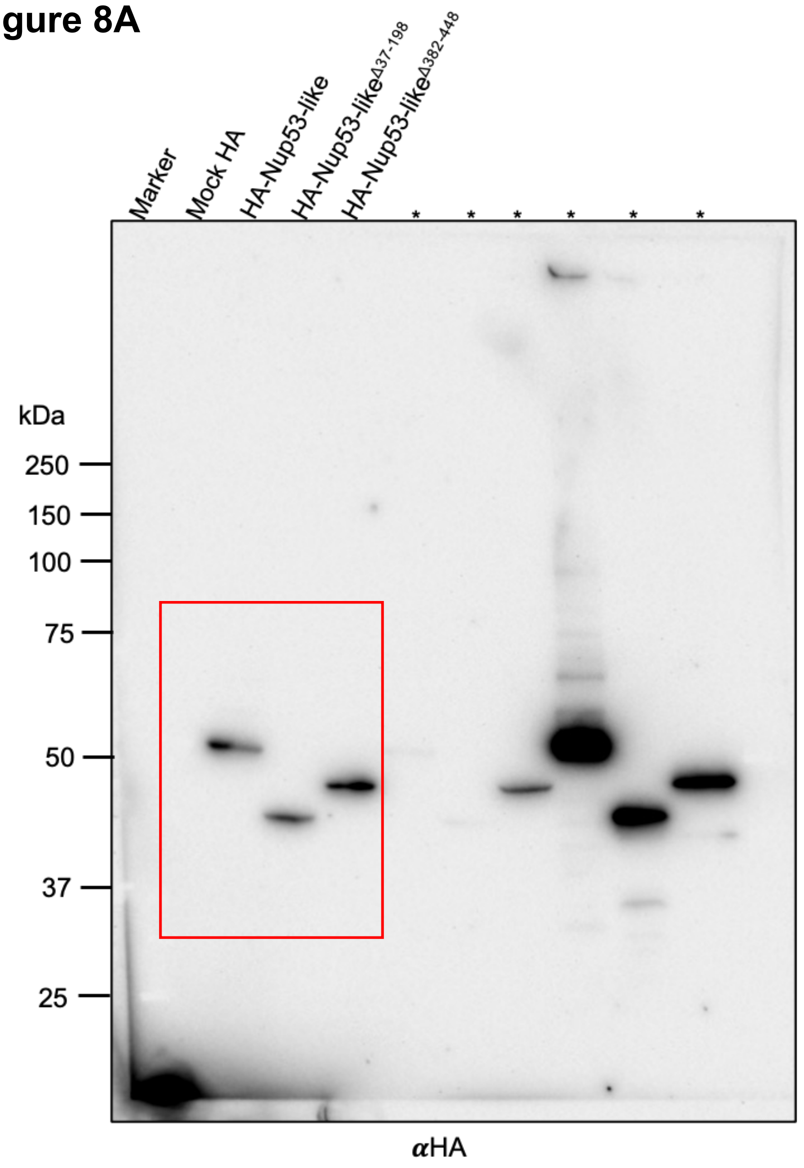
Marker
Mock HA
HA-Nup53-like
HA-Nup53-likeΔ37-198
HA-Nup53-likeΔ382-448
kDa
250
150
100
75
50
37
25
αHA

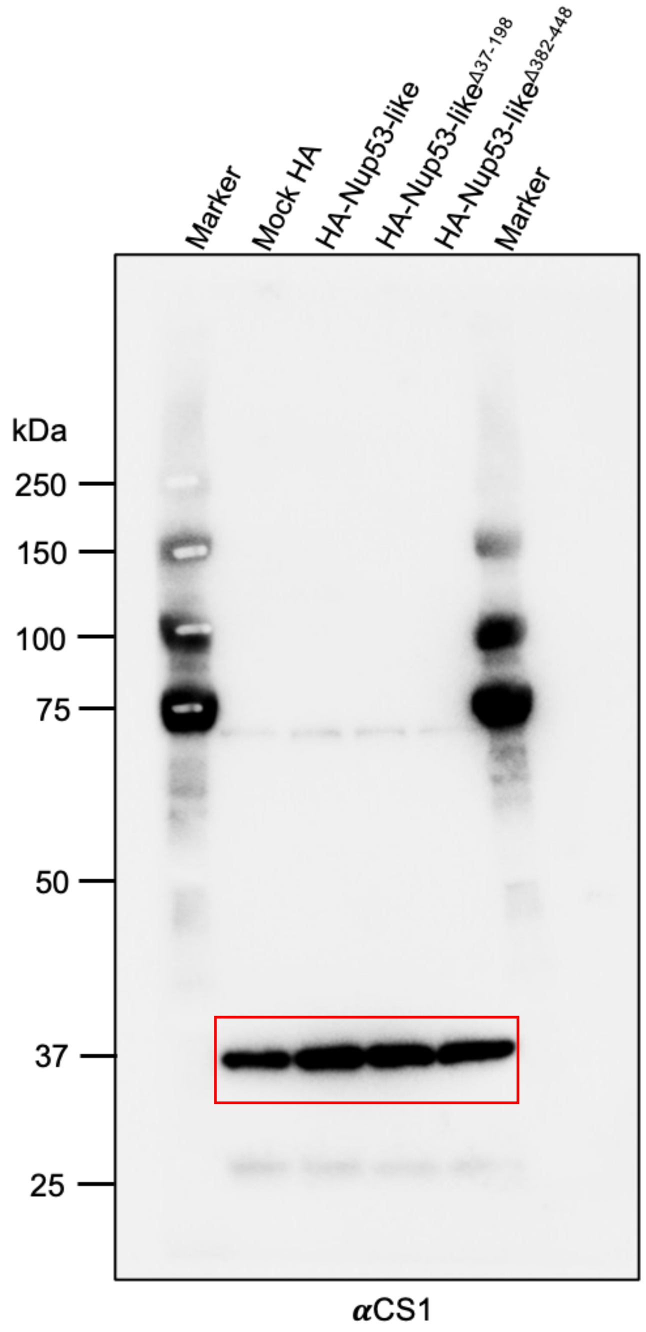
Marker
Mock HA
HA-Nup53-like
HA-Nup53-likeΔ37-198
HA-Nup53-likeΔ382-448
Marker
kDa
250
150
100
75
50
37
25
αCS1

**Supplementary Figure 2A**

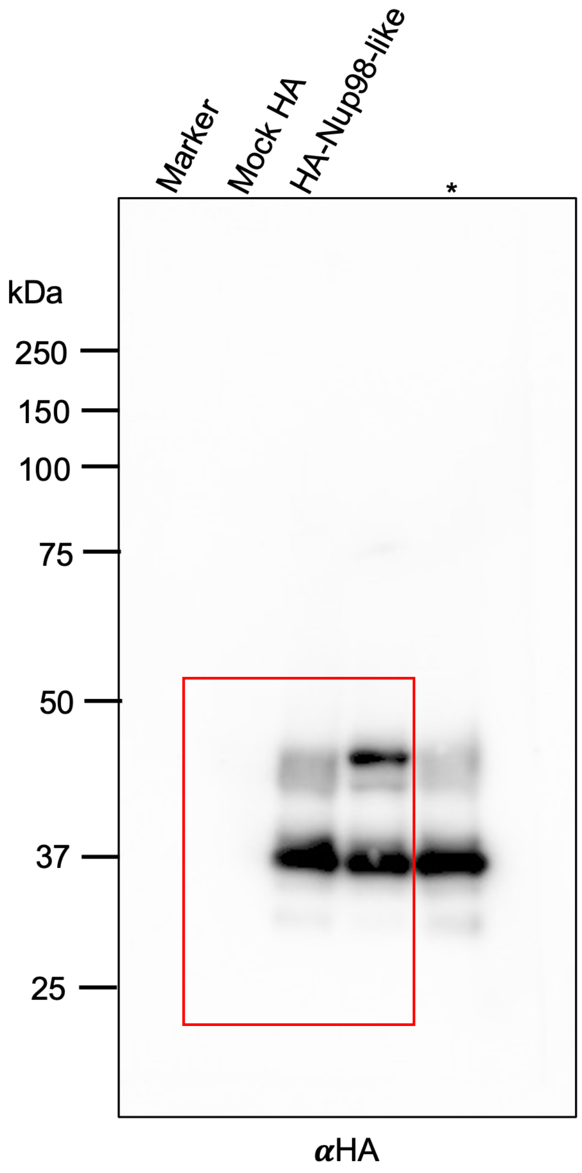

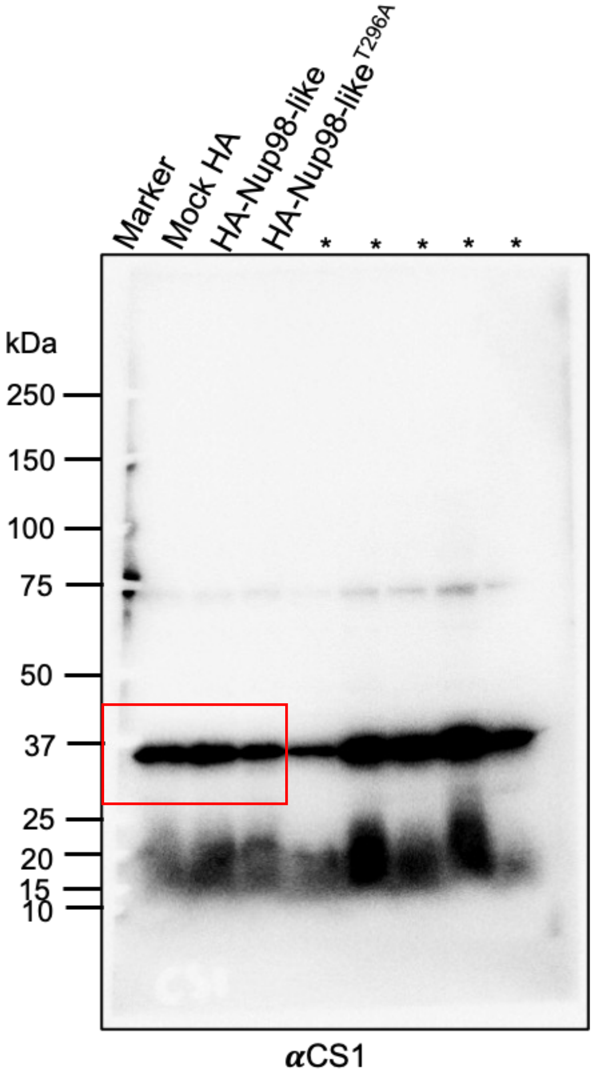
